# Beyond linear dynamic functional connectivity: a vine copula change point model

**DOI:** 10.1101/2021.04.25.441254

**Authors:** Xin Xiong, Ivor Cribben

## Abstract

To estimate dynamic functional connectivity for functional magnetic resonance imaging (fMRI) data, two approaches have dominated: sliding window and change point methods. While computationally feasible, the sliding window approach has several limitations. In addition, the existing change point methods assume a Gaussian distribution for and linear dependencies between the fMRI time series. In this work, we introduce a new methodology called Vine Copula Change Point (VCCP) to estimate change points in the functional connectivity network structure between brain regions. It uses vine copulas, various state-of-the-art segmentation methods to identify multiple change points, and a likelihood ratio test or the stationary bootstrap for inference. The vine copulas allow for various forms of dependence between brain regions including tail, symmetric and asymmetric dependence, which has not been explored before in the dynamic analysis of neuroimaging data. We apply VCCP to various simulation data sets and to two fMRI data sets: a reading task and an anxiety inducing experiment. In particular, for the former data set, we illustrate the complexity of textual changes during the reading of Chapter 9 in *Harry Potter and the Sorcerer’s Stone* and find that change points across subjects are related to changes in more than one type of textual attributes. Further, the graphs created by the vine copulas indicate the importance of working beyond Gaussianity and linear dependence. Finally, the R package **vccp** implementing the methodology from the paper is available from CRAN.

## 1 Introduction

Functional magnetic resonance imaging (fMRI) experiments yield high dimensional data sets that contain complex spatial correlations, often referred to as functional connectivity (FC) networks (see for example, Cribben and Fiecas, 2016, for a review). These FC networks have been previously studied to expose important characteristics of brain function and individual variations in cognition and behavior. In particular, Greicius et al. (2004), Menon (2011), Bakhtiari et al. (2017) and Hart et al. (2018) showed that neurological disorders disrupt the FC structure and Hrybouski et al. (2021) studied changes in FC in healthy aging subjects.

Recently, there has been a surge in the development of new statistical methods for investigating how FC networks change over time. These changes are commonly referred to as time-varying, or dynamic FC in neuroimaging. This reconstruction of dynamic FC has a major impact on the understanding of the functional organization of the brain. Similar to other biological networks, understanding the complex, dynamic organization and characterizations of the brain can lead to profound clinical implications (Bullmore and Sporns, 2009).

To estimate dynamic FC for fMRI data, two approaches have dominated: sliding window and change point methods. For the former, a sliding window of pre-specified length is defined and the correlations between distinct regions of the brain throughout the duration of the window are estimated. While the sliding window approach is computationally feasible, it also has limitations (Hutchison et al., 2013). For example, the choice of window size is crucial and sensitive, as different window sizes can lead to quite different FC patterns. Another disadvantage is that equal weight is given to all *k* neighbouring time points and 0 weight to all the others. Hence, researchers considered change point methods that partition the time series into optimal windows. Determining change points may also reveal properties of brain networks as they relate to experimental stimuli and disease processes. There exists an extensive literature and a long history on change point detection. The most widely discussed problems have been concerned with finding multiple change points in univariate time series (Inclan and Tiao, 1994; Chen and Gupta, 1997). Recently, the multiple change point detection problem in multivariate time series has received some attention especially in non-stationary practical problems. To detect changes in the covariance matrix of a multivariate time series, Aue et al. (2009) introduced a method using a nonparametric CUSUM type test, Dette and Wied (2016) proposed a test where the dimension of the data is fixed while Kao et al. (2018) considered the case where the dimension of the data increases with the sample size. Sundararajan and Pourahmadi (2018) proposed a new method for detecting multiple change points in the covariance structure of a multivariate piecewise-stationary process, while Dette and Gösmann (2020) developed a likelihood ratio approach to detect change points for a general class of parameters including changes in correlations.

In other work, Barnett and Onnela (2016) considered a method for detecting change points in correlation networks. Gibberd and Nelson (2014) identifies both change points and the graphical dependency structure in multivariate time series. Li et al. (2019) considered multiple structural breaks in large contemporaneous covariance matrices of high dimensional time series satisfying an approximate factor model. Cho and Fryzlewicz (2015) segmented the multivariate time series into partitions based on the second-order structure.

For neuroscience applications, Cribben et al. (2012, 2013) first introduced the idea of detecting FC change points by introducing Dynamic Connectivity Regression for detecting multiple change points in the precision matrices (undirected graphs) from a multivariate time series. Schröder and Ombao (2019); Kirch et al. (2015); Cribben and Yu (2017); Kundu et al. (2018); Dai et al. (2019); Ofori-Boateng et al. (2021); Ondrus et al. (2021); Anastasiou et al. (2022) among others have since introduced new network change point methods. However, all of these methods have limitations. The most obvious is that they all consider a Gaussian distribution for and linear dependencies between the fMRI time series.

In this paper, we introduce a new methodology, called *Vine Copula Change Point* (VCCP), to estimate multiple change points in the FC structure between brain regions. The new method combines vine copulas, various state-of-the-art segmentation methods to identify multiple change points, and a likelihood ratio test or the stationary bootstrap for inference. The proposed VCCP method has the following unique and significant attributes. First, it is the first statistical method that has applied vine copulas to neuroimaging data. (Fontaine et al., 2020, apply copula models to local field potential data of rats and focus on static dependence). Second, it is the first statistical method that considers change points in vine copulas. Vine copulas split the multivariate distribution into marginal distributions and a dependence measure without a need to assume that the data follows a parametric distribution (e.g., Gaussian). Hence, VCCP is very flexible. Third, VCCP is the first method that allows us to describe various forms of dependence structures such as tail, symmetric and asymmetric dependence between time series in a dynamic fashion, which has not been explored before in the analysis of neuroimaging data. Therefore, VCCP is capable of modeling a wider range of dependence patterns and hence allows for the detection of more change points. Fourth, the layers of a vine (the number of trees) can be cut, which leads to a simplified and sparse dependence matrix which reduces computation time when the dimension of the data expands. Fifth, VCCP allows for the exploration of network dynamics during a reading fMRI experiment (Chapter 9 in *Harry Potter and the Sorcerer’s Stone*) and an anxiety inducing fMRI experiment. In the former, it detects change points across subjects that coincide to more than one type of textual attributes. Further, the graphs created by the vine copulas indicate the importance of working beyond Gaussianity and linear dependence. Sixth, as VCCP uses various state-of-the-art segmentation methods, the paper can be viewed as an extensive comparison of these methods. Seventh, while motivated by fMRI data, VCCP could also be applicable to electroencephalography (EEG), magnetoencephalography (MEG) and electrocorticography (ECoG) data, and other time series applications where the network structure is changing. Finally, the R package **vccp** implementing the VCCP methodology is available from CRAN (Xiong and Cribben, 2021).

This paper is organized as follows. In Section 2 we provide a background on copulas, vine copulas and explain the setup of our proposed new methodology, Vine Copula Change Point (VCCP). In Section 3, we describe the simulated data sets with known change points locations and the two fMRI data sets. We present the performance of VCCP in Section 4, have a discussion in Section 5 before concluding in Section 6.

## 2 Methods

### 2.1 Copulas

Table 1 provides a summary of the notation used in the paper. We now introduce copulas and vine copulas. A copula “couples” marginal distributions into a joint distribution. Hence, copulas allow for the independent construction of joint distributions and marginal distributions. This is convenient as marginal distributions in many cases can be adequately estimated from data, whereas dependence information involves summary indicators and judgment. More specifically, let **X** = (*X*_1_, …, *X_p_*) be a *p*-dimensional random variable or a multivariate time series from *p* regions of interest (ROIs) with cumulative distribution function (cdf) *F* (*x*_1_, …, *x_p_*). A copula, which is also a multivariate cdf, serves as the link that connects the marginal distributions of **X** to its multivariate cdf, *F* (*x*_1_, …, *x_p_*). A formal definition of a copula is given by:

**Table 1:** Notation for the Vine Copula Change Point (VCCP) model.

| Index | Meaning |
| --- | --- |
| $p$ | Data dimension |
| $X = (x_1, x_2, \dots, x_p)$ | A $p$-dimensional random variable |
| $C(\cdot, \dots, \cdot)$ | A copula function |
| $c(\cdot, \dots, \cdot)$ | A copula density function |
| $n$ | Sample size |
| $T$ | The length of a time series |
| $\delta$ | The minimum distance between two adjacent candidate change points |

#### Definition 1 (Copula)

*A p-dimensional copula, C, is a multivariate cdf defined on* [0, 1]^*p*^, *C* : [0, 1]^*p*^ → [0, 1], *and its univariate margins have a uniform distribution*.

According to Sklar (1959), assuming the marginals of **X** are continuous, every continuous cdf *F* has a unique copula *C* that satisfies

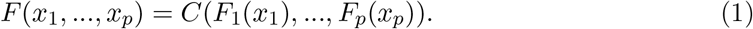

The availability of high-dimensional copula models is limited, but there are several parametric bivariate copula models such as Gaussian, *t*, Clayton, Gumbel, and Frank copula (see the Supplementary Materials for definitions) to name but a few (the accompanying R package **vccp** allows for many more choices of copulae). These have led to the development of hierarchical models, constructed from cascades of bivariate copula models, called pair-copula constructions (PCCs) or vine copulas.

### 2.2 Vine copulas

PCCs is a method for building a multivariate distribution by firstly decomposing a joint density function into a sequence of conditional densities, such as

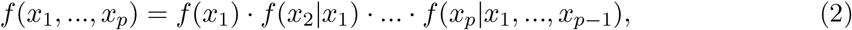

making the construction of a possible complex dependence structure both flexible and tractable. Then using the definition of conditional densities and the derivative of (1), Joe (1996) proved that *f* (*x*_1_, …, *x_p_*) can be further decomposed to:

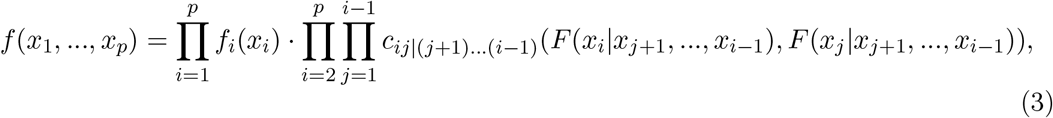

where *f_i_* denotes the marginal density of *x_i_*, *F* (*x_i_*|*x_j_*_+1_, …, *x_i−_*_1_) is its univariate conditional distribution function, and *c*_*ij*|(*j*+1)…(*i*−1)_ is the density of the conditional pair copula associated with the bivariate conditional distribution of *x_i_* and *x_j_* given the subset, (*x*_*j*+1_, …, *x*_*i*−1_). There are many other decompositions of the joint density based on the assigned conditional sets in (2). In fact, Morales Napoles (2016) proved that there are 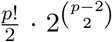 possible decompositions for a *p*-dimensional distribution. Bedford and Cooke (2002) organized the PCCs decomposition as a one-to-one graphical model, a sequence of *p* − 1 nested trees (where edges correspond to the bivariate copulas), called a regular vine, or R-vine (see Kurowicka and Cooke 2006 for more details). For tree *m* (*m* = 1,…, *p* − 1), define *T*_*m*_ = (*V*_*m*_, *E*_*m*_), where *V*_*m*_ and *E*_*m*_ represent the node set and edge set, respectively, then the tree sequence (called the structure of the PCC) is an R-vine if it satisfies a set of conditions guaranteeing that the decomposition is a valid joint density. To specify a *p*-dimensional distribution on an R-vine structure, 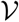, with a node set 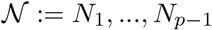 and an edge set *ε* ≔ *E*_1_, …, *E*_*p*−1_, we need to connect each edge *e* = *j*(*e*), *k*(*e*)|*D*(*e*) in *E_i_* to a conditional bivariate copula density *c*_*j*(*e*),*k*(*e*)|*D*(*e*)_, where *j*(*e*) and *k*(*e*) are the <u>conditioned set</u>, while *D*(*e*) is the <u>conditioning set</u>. Finally, the joint copula density can be written as the product of all pair-copula densities 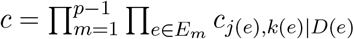. To estimate vine copulas, it is common to follow a sequential approach from higher trees (more nodes) to lower trees (less nodes), since there are 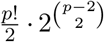 possible R-vine structures for a *p*-dimensional cdf. If the vine structure is known, the pair-copulas of the first tree, *T*_1_, can be estimated directly from the data. This is not possible for the other trees in the sequence, since data from the densities *c*_*j*(*e*),*k*(*e*)|*D*(*e*)_ are not observed. However, it is easy to create “pseudo-observations” using appropriate data transformations, resulting in the following estimation procedure: estimate tree *T*_1_ directly from the data, for each edge in the tree, estimate all pairs, construct pseudo-observations for the next tree and then iterate. As the tree sequence *T*_1_, *T*_2_, …, *T*_*p*−1_ is a regular vine, it guarantees that at each step, all required pseudo-observations are available. In this work, in order to specify every tree’s connecting network, we utilize Algorithm 1 (Dissmann et al., 2013). For more information on vines and its extensions, see Aas (2016); Czado (2019).

#### Algorithm 1: A sequential method to select a vine copula model

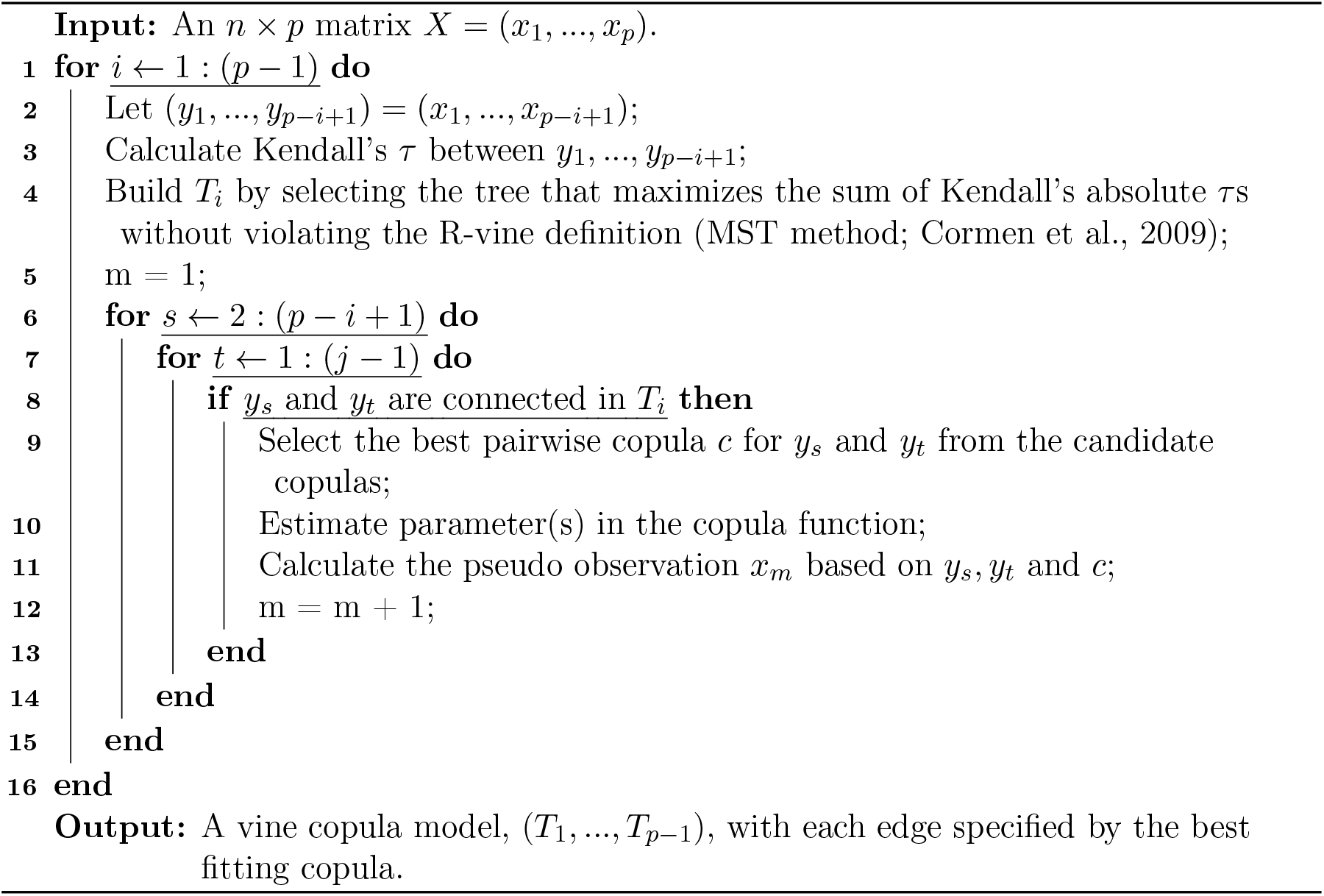

To illustrate how to compute the pseudo observations, suppose we need to fit a vine copula model to a 4-dimensional distribution. The first tree (*T*_1_) is composed of 4 nodes ({*x*_1_, *x*_2_, *x*_3_, *x*_4_}) and three connected edges ({*x*_1_, *x*_2_}, {*x*_2_, *x*_3_}, {*x*_3_, *x*_4_}). Three copulas are fitted to the corresponding three edges which we denote by *c*_12_, *c*_23_, *c*_34_. For the second tree (*T*_2_), we transform edges in the last tree to nodes and then find the best copula between the two transformed nodes. For example, to fit a copula to edge *e* = {*a, b*} where *a* = {*x*_1_, *x*_2_} and *b* = {*x*_2_, *x*_3_}, we first find the conditioning set ({*u*_1_, *u*_2_} = {*x*_1_, *x*_3_}) and conditioned set ({*v*} = {*x*_2_}) of *e* (Dissmann et al., 2013). We obtain 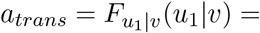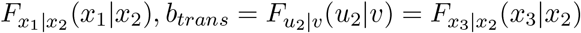 where 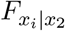 can be derived from 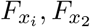 and *c_i2_* (*i* = 1, 3). Then we find the best copula that minimizes a certain criterion of the bivariate model on 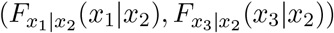. Such a transformation is carried recursively as the layers of the tree increases.

In our work, the candidate set of copulas include Gaussian, *t*, Clayton, Gumbel and Frank copula (see the Supplementary Materials for definitions). The best copula type is selected according to the Bayesian Information Criteria (BIC), which is equal to 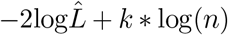, where 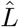 is the maximized value of the likelihood function of the vine copula model, *n* is the number of observations and *k* is the number of parameters estimated in the vine copula. First, all available bivariate copulas are fitted to the (pseudo) observed data using maximum likelihood estimation. Then the BICs are computed for all available copula families and the family with the minimum value is chosen.

Two special cases of an R-vine are the canonical-vine (C-vine) and the drawable-vine (D-vine). An R-vine is called a *C-vine* if there is one unique node (center node) of degree *p* − *i* in each tree *T_i_* (*i* = 1, …, *p* − 1) (in every tree, one node serves as the center that links all the other nodes). An R-vine is called a *D-vine* if the maximal degree of freedom of nodes in each tree is 2 (no node in any tree can be linked with more than 2 other nodes). Figure 1 shows examples of a D-vine and a C-vine that are based on 5 variables (or 5 ROIs). The colored point on each tree of the C-vine represents the center node, which is also indicated by the dashed line on the previous tree.

**Figure 1:**
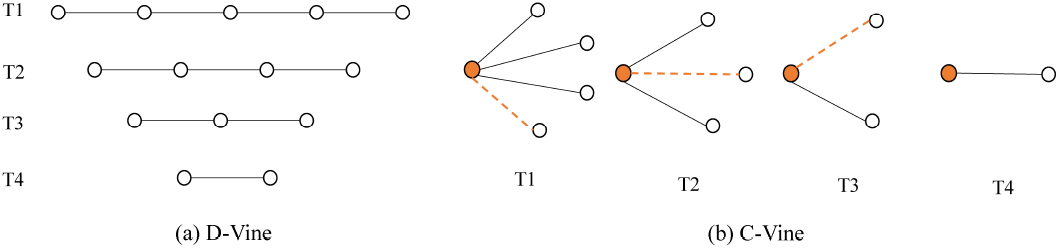
A connected graph for a D-vine (left) and a C-vine (right) that is based on 5 variables (or 5 ROIs).

### 2.3 Segmentation Methods

In this work, we are not only interested in estimating vine copulas but in estimating change points in the vine copula structure, where we assume that those change points are discrete and at some distance from each other. In order to find the change points, VCCP uses the Bayesian Information Criterion (BIC: Schwarz et al., 1978) metric. BIC is equal to 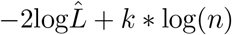, where 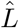 is the maximized value of the likelihood function of the vine copula model, *n* is the number of observations and *k* is the number of parameters estimated in the vine copula. However, a time axis of length *T* and *r* change points can be divided into 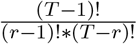 possible sub-partitions (with unknown *r*). It is clear that as *T* increases, the number of sub-partitions explodes. Hence, VCCP uses four segmentation methods to simplify the process, which we now detail.

Binary segmentation (BS) is a generic segmentation method. Assuming an unknown number of change points, BS can be used to find multiple change points with low computational complexity. BS begins by searching the entire time course for one change point. We set an auxiliary time point *i* that shifts from (*δ* + 1) to (*T* − *δ* − 1), where *i* indicates a candidate change point and *δ* is a parameter that represents the minimal distance between two candidate change points. *δ* needs to be sufficiently large enough to estimate a stable R-vine structure, but also small enough in order to not miss candidate change points. Thus, three R-vine copula specifications are estimated on the data: between time points (1 : *i* − 1), (*i* : *T*) and (1 : *T*). The reduced BIC as a result of partitioning the time course into two intervals is calculated as *i* moves across time. We choose time *t*_1_, which corresponds to the largest BIC reduction as the first candidate change point. After inference, if it is deemed a change point, the time course is split into two sub-intervals (where the binary name originates from) and a similar search is repeated on each of them. The recursion does not stop until BIC does not decrease by partitioning the time course, or the candidate point corresponding to the maximum BIC reduction fails the inference test. An example of BS is shown in Figure 2. As ‘arguably the most widely used change point search method’, the BS method is conceptually simple and is very easy to use. Despite this, as it is a ‘greedy’ procedure, each step except the first depends on the previous steps, which are never re-visited. This sequential method leads to inconsistent estimates when the minimum spacing between two adjacent change points is of order less than *T* ^3/4^. We propose an adapted BS method (NBS), which is similar to the old classic BS (OBS), but does not carry out inference on each change point until the segmentation has been exhausted (the stopping criteria for NBS is whether the maximal BIC reduction is larger than 0). Once no further candidate change points reduce the BIC score, NBS orders the candidate change points in sequence and recalculates the reduced BIC:

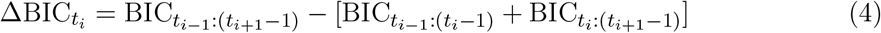

where 1 = *r*_0_ < *r*_1_ < … < *t*_*r*_ < *t*_*r*+1_ = *T* and *t*_1_,…, *t_r_* are the candidate change points detected by NBS. Then, inference is performed on each non-zero ΔBIC in partitions [*t*_*i*−1_, *t*_*i*+1_].

**Figure 2:**
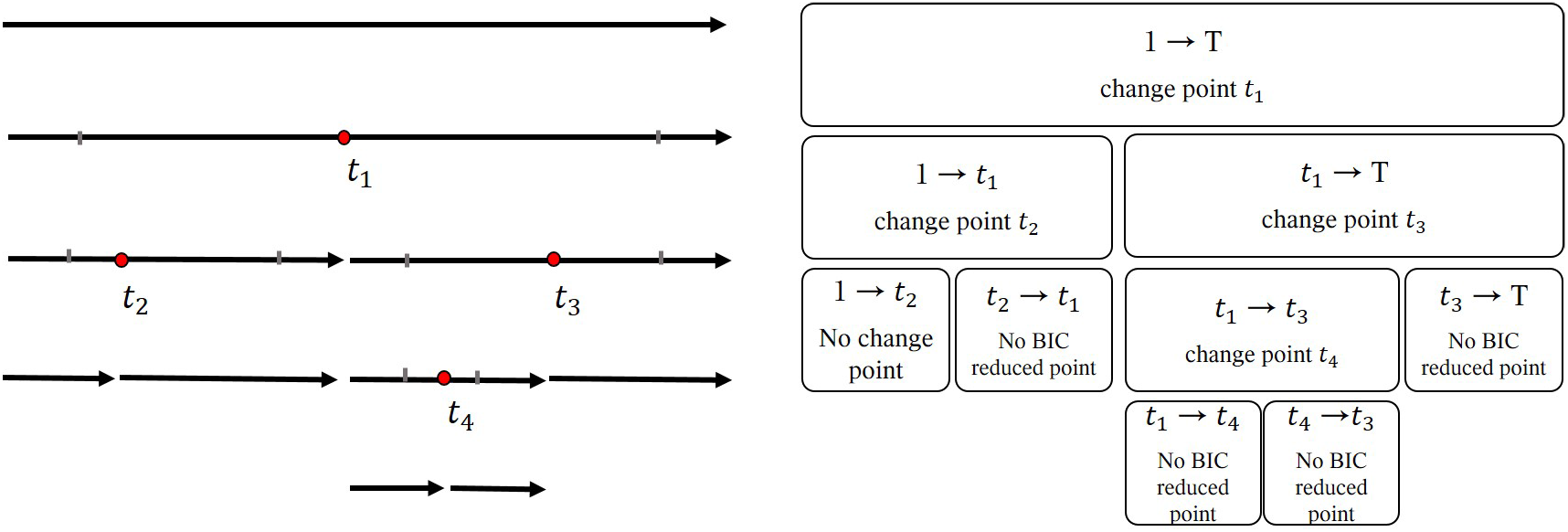
An example of Binary Segmentation (BS). Here, *t*_1_ corresponds to the first candidate change point. It is deemed a change point, hence the data is then split into two sub-intervals. On each sub-segment, we restart a similar search to further segment the data.

VCCP also includes more recent segmentation methods, Wild Binary Segmentation (WBS: Fryzlewicz, 2014) and moving sum (MOSUM: Eichinger and Kirch, 2018). Briefly, WBS starts by randomly drawing *S* sub-samples 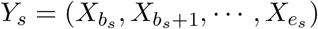, where *b_s_*, *e_s_* and *s* are integers such that 1 ⩽ *b_s_ < e_s_* ⩽ *T*, *s* = 1, …, *S*. Then it estimates candidate change point 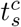 on each sub-sample *Y_s_* based on a single binary search (see *t*_1_ in Figure 2) and a corresponding statistic, 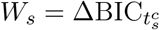. After, WBS selects the candidate change point as

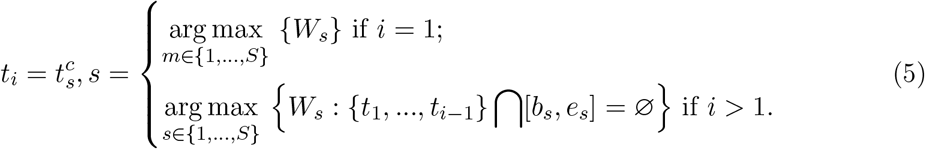

This segmentation method exhausts when *t_i_* is not a change point or each random interval ([*b_s_, e_s_*]) covers at least one detected change point. To simplify computation, we remove sub-intervals that contain less than 2*δ* time points before computing the BIC reduction. By ‘randomly localizing’ the range of BIC reductions, we overcome the issue of greedy searching for certain configurations of multiple change points. Also, since intervals are drawn randomly, we avoid issues such as choosing the appropriate window size prevalent in other segmentation methods. We set *S* as the minimum number of random draws needed to ensure that the bound on the speed of convergence for the estimated number and the location of change points is suitably small. According to Theorem 3.2 in Fryzlewicz (2014), under some restriction on the magnitudes of the jump between BIC in different intervals and the minimum spacing between change points, we have 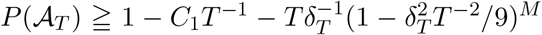 for certain positive *C*_1_, where 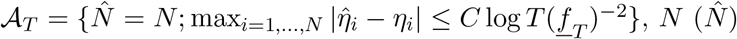 the true (estimated) number of change points, 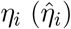 the location of *i^th^* true (estimated) change point and 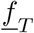 the lower bound for the BIC jump. In order to match the rate of the term *C*_1_*T* the upper bound for 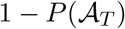, we need to ensure in 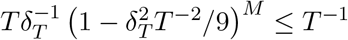 which is equivalent to:

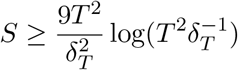

where *δ_T_* is the minimum distance between two change points.

MOSUM (Eichinger and Kirch, 2018) detects change points in the mean and variance based on the moving sum statistics for a univariate time series. For multivariate time series with a changing network structure, it is not possible to directly apply MOSUM. Hence, we adapt MOSUM for VCCP as follows. First, similar to NBS, we set an auxiliary time point *i* that shifts from [*δ* + 1] to [*T* − *δ* − 1]. For every fixed *i*, three vine copula specifications are estimated on the data: between time points [*i*−1−*δ* : *i*−1], [*i* : *i*+*δ*] and [*i*−1−*δ* : *i*+*δ*]. For MOSUM, the two intervals have the same length (*δ*), while for NBS the window size varies. The reduced BIC sequence {BICr_*i*_}_*i*=1*,…,T −*2*δ*_ can be easily incorporated into a MOSUM model. For the univariate time series BICr, consider the following moving sum statistic:

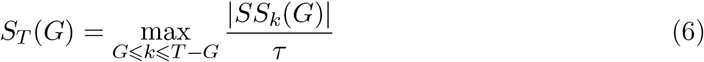

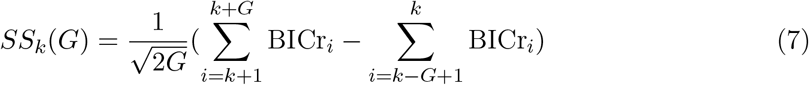

where *G* represents the bandwidth that is selected beforehand. *SS_k_*(*G*) compares the mean of the sub-sample BICr_*k−G*_, …, BICr_*k*_ with the mean of the sub-sample BICr_*k*+1_, …, BICr_*k*+*G*_, where a large difference indicates the presence of a candidate change point at time point *k*. Hence, we adapt MOSUM into our VCCP model as follows:

1. Calculate the reduced BIC sequence within a local range [*i* − *δ* + 1 : *i* + *δ*].
2. Apply MOSUM to the time series, resulting in several boundary points (*m*_1_, …, *m_k_*).
3. Separate the whole time axis into intervals [*m_i_, m_i_*_+1_], *i* = 0, …, *k, m*_0_ = 1, *m_k_*_+1_ = *T* and record time points *t*_1_, …, *t_k_* corresponding to the local maximal reduced BIC in the different intervals.

### 2.4 Inference

To perform inference on the candidate change points, VCCP uses the Vuong test (Vuong, 1989) or the stationary bootstrap (SB: Politis and Romano, 1994). The Vuong test is used to compare non-nested models. In our R-vine setting, non-nested models can refer to different copulas for edges of the same tree structure, different vine structures, or a combination of the two. For a random variable *X*, the Vuong test uses the Kullback-Leibler information criterion to compare two models *F_β_* = *f* (*X*|*β*) and *G_γ_* = *g*(*X*|*γ*). Suppose *β*_∗_ and *γ*_∗_ are the true parameters of the two candidate models and 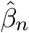 and 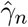 are their corresponding maximum likelihood estimates. Then the null hypothesis is

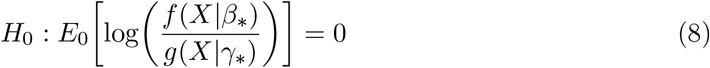

where *E*_0_ is the expectation under *H*_0_, and the likelihood ratio test statistic is

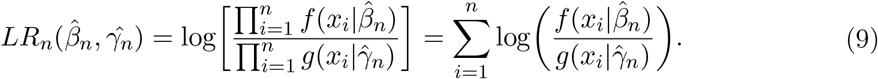

Vuong (1989) proved that this test statistic is asymptotically normal, which leads to the asymptotic test, 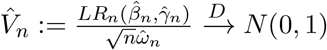 as *n* → ∞ and

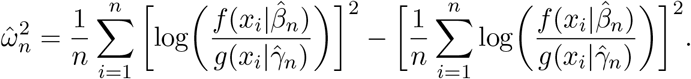

If we do not reject *H*_0_ under significance level *α*, the conclusion is that both models fit the data equally well. On the contrary, if we were to reject *H*_0_ and the statistic 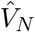 is positive, the test suggests that model *F_β_* on the numerator of the likelihood ratio statistic is superior, while a negative value supports model *G_γ_* on the denominator. Similar to BIC, Vuong (1989) included the following a penalty term in the likelihood ratio statistic, 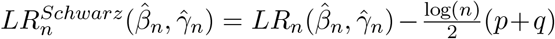, where *p* and *q* are the number of parameters in models *F_β_* and *G_γ_*, respectively.

The SB is an adaptation of the block bootstrap, which allows us to deal with the auto-correlation in time series by retaining the dependence structure of the data. Unlike resampling blocks of data with a fixed block size in the block bootstrap, the SB randomly specifies each block size. For any strictly stationary time series {*X_t_, t* = 1, …, *T*}, the SB procedure consists of generating pseudo-samples 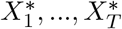 from the sample *X*_1_, …, *X_T_* by taking the first *T* elements from 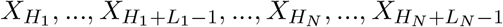 where *H_i_* is an independent and identically distributed sequence of random variables uniformly distributed on {1,…,T} and *L_i_* is an i.i.d. sequence of geometrically distributed random variables with *P* (*L_i_* = *h*) = *q*(1 − *q*)^*h−*1^, *h* = 1, 2, …, for some *q* ∈ (0,1). Note that the mean block size is 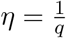, where the choice of *η* is similar to the choice of block length in the block bootstrap.

After the change points have been detected, in order to account for multiple comparisons, we perform a Benjamini Hochberg correction (Benjamini and Hochberg, 1995) to the *p*-values.

## 3 Simulated and fMRI data descriptions

In order to show the performance of our VCCP methodology we use several simulated data sets, and two task-based fMRI data sets. To carry out inference on these data sets, we have to adjust the Vuong test (Section 2.4) when we combine it with the segmentation methods. For NBS and MOSUM, the first step is to detect all candidate change points 1 < *t*_1_ < *… < t_k_ < T*. For interval *I_i_* = [*t_i−_*_1_, *t_i_*_+1_], three vine copulas are estimated based on data in the intervals [*t_i−_*_1_, *t_i_*_+1_], [*t_i−_*_1_, *t_i_* −1], [*t_i_, t_i_*_+1_], which we denote by *V C*_0_, *V C_l_* and *V C_r_*. In each interval, the data are assumed to be i.i.d and that there are no other change points between time points *t_i−_*_1_ and *t_i_*_+1_ except *t_i_*. If *H*_0_: *t_i_* is not a change point in [*t_i−_*_1_, *t_i_*_+1_] is rejected, then constructing two VC models in the two partitions separately provides a superior fit to the data than a VC model using the entire data set. Hence, we perform a Vuong test on *V C*_0_ and *V C_l_* based on data in [*t_i−_*_1_, *t_i_* − 1] and a Vuong test on *V C*_0_ and *V C_r_* using data in the interval [*t_i_, t_i_*_+1_]. If both tests show that *V C*_0_ is worse than either *V C_l_* or *V C_r_*, we conclude that *t_i_* is a change point. For OBS, the only difference in the above process is that the left and right end of the intervals are the endpoints of the partitioned data in each BS search, since OBS runs detection and inference sequentially. For WBS, the intervals are based on the endpoints of randomly chosen sub-samples. The rest of their procedures are the same as NBS and MOSUM. The only difference between SB and the Vuong test is that instead of conducting likelihood-based inference, SB creates SB pseudo-samples on [*t_b_, t_e_*] and compares them to the original BIC reduction value, 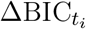, where *t_i_* is the candidate change point, [*t_b_, t_e_*] = [*t_i−_*_1_, *t_i_*_+1_] for NBS and MOSUM, 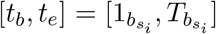 for OBS and 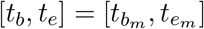 for WBS.

### 3.1 Simulated data

We include three data types: multivariate normal (MVN) data, data from a vector autoregression (VAR) model, and non-Gaussian (or Vine copula) data. The descriptions (and results) of the first two data types are contained in the Supplementary Materials. We run each simulation using 100 iterations. We now describe the non-Gaussian (or vine copula) simulations. To illustrate the matrix representation of simulation settings with tail dependence change points, we take as a toy example a 4-layer D-vine copula. In Figure 3(a), we depict a 4-layer D-vine as a nested set of connected trees. As storing the nested set of trees is too expensive, we instead can use a convenient matrix representation that encodes this D-vine (or R-vine) structure. An edge in the *k^th^* layer can be represented by ‘*i, j*|*m*_1_, …, *m_k−_*_1_’, where {*m*_1_, …, *m_k−_*_1_} denotes the conditioned set, {*i, j*} the conditioning set, and ({*i, j*}|*m*_1_, …, *m_k−_*_1_) is one element in the constraint set. For more detailed definitions of the conditioned, conditioning and constraint sets, see Dissmann et al. (2013). Since there is a one-to-one correspondence between the constraint set and the nested set of connected trees for a vine, the idea here is to store the constraint set of the D-vine in columns of an *d*-dimensional lower triangular matrix *M* = {*m_ij_*}_*i,j*=1*,…,d*_.

**Figure 3:**
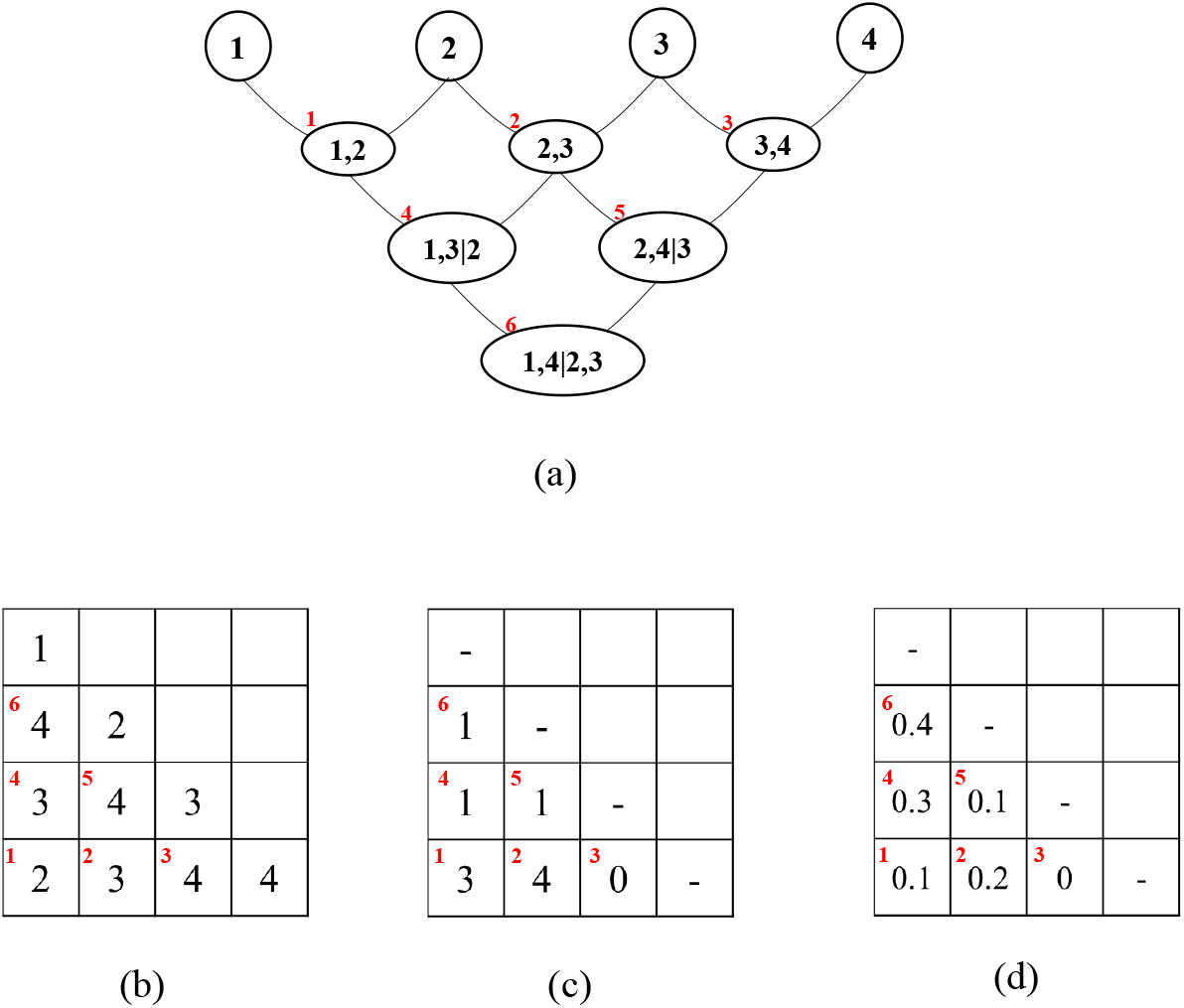
A toy example to explain the simulation settings. (a) A 4-layer D-vine depicted as a nested set of connected trees; (b) The corresponding vine copula structure matrix for (a); (c) The copula family matrix for (a); (d) The parameter matrix for (a).)

#### Definition 2 (Matrix constraint set)

*Let M* = {*m*_*ij*_}_*i,j*=1,…,*d*_ *be a lower triangular matrix. The i-th constraint set for M is C_M_* (*i*) = {({*m_i,i_, m_k,i_*, }, *D*)|*k* = *i* + 1, …, *d, D* = {*m_k_*_+1*,i*,…,*m*_*d,i* }} *for i* = 1, …, *d* − 1. *If k* = *d we set D* = ∅. *The constraint set for matrix M is the union CM* = *C_M_* (1)⋃ … ⋃ *C_M_* (*d* − 1). *For the elements of the constraint set* ({*m_i,i_, m_k,i_*}, *D*) ∈ *CM, we call* {*m_i,i_, m_k,i_*} *the conditioned set and D the conditioning set*.

In other words, every element of the constraint set is made up of an diagonal entry *m_i,i_*, an entry in the same column below the diagonal *m_k,i_* and all the elements following in that column {*m_k_*_+1*,i*_, …, *m_d,i_*}. As long as we specify one element on the lower triangle matrix *m_k_*_+1*,i*_, based on its column number we can find the diagonal entry *m_i,i_* as another node in the conditioned set and all its following elements {*m_k_*_+1*,i*_, …, m_d,i_} as the conditioning set. Therefore, *m_k_*_+1*,i*_ for *i* = 1, …, *d* − 1 and *k* = *i, …, d* − 1 can represent all edges in a D-vine. If we need to assign attributes to pairs of nodes in a D-vine, to build the vine copula model, we can refer to the lower triangle matrix and put any related values of {*m_i,i_, m_k_*_+1*,i*_|(*m_k_*_+2*,i*_, …, m_*d,i*_)} to *m_k_*_+1*,i*_. To complete this, we need to specify the copula family (e.g., Figure 3)(c)) and copula parameter(s) (e.g., Figure 3)(d)) for each edge. For example, the 1, 3|2 node on the D-vine in Figure 3)(a), we can see that it corresponds to element *m*_3,1_ in the matrix. From Figure 3(c) we know then that the copula type for 1, 3|2 is the Gaussian copula and from Figure 3(d) the parameter is 0.3, given element *m*_3,1_ in the copula family matrix is 1 and *m*_3,1_ = 0.3 in the parameter matrix.

With this understanding, Figure 4 provides a visual display of Simulation 1 (no change point in the vine copula structure). In this simulation there are *P* = 10 time series and *T* = 140 time points. Here, the D-vine structure, Kendall’s *τ*, the upper and lower tail dependence structures, the copula type and the parameter values are constant over time.

**Figure 4:**
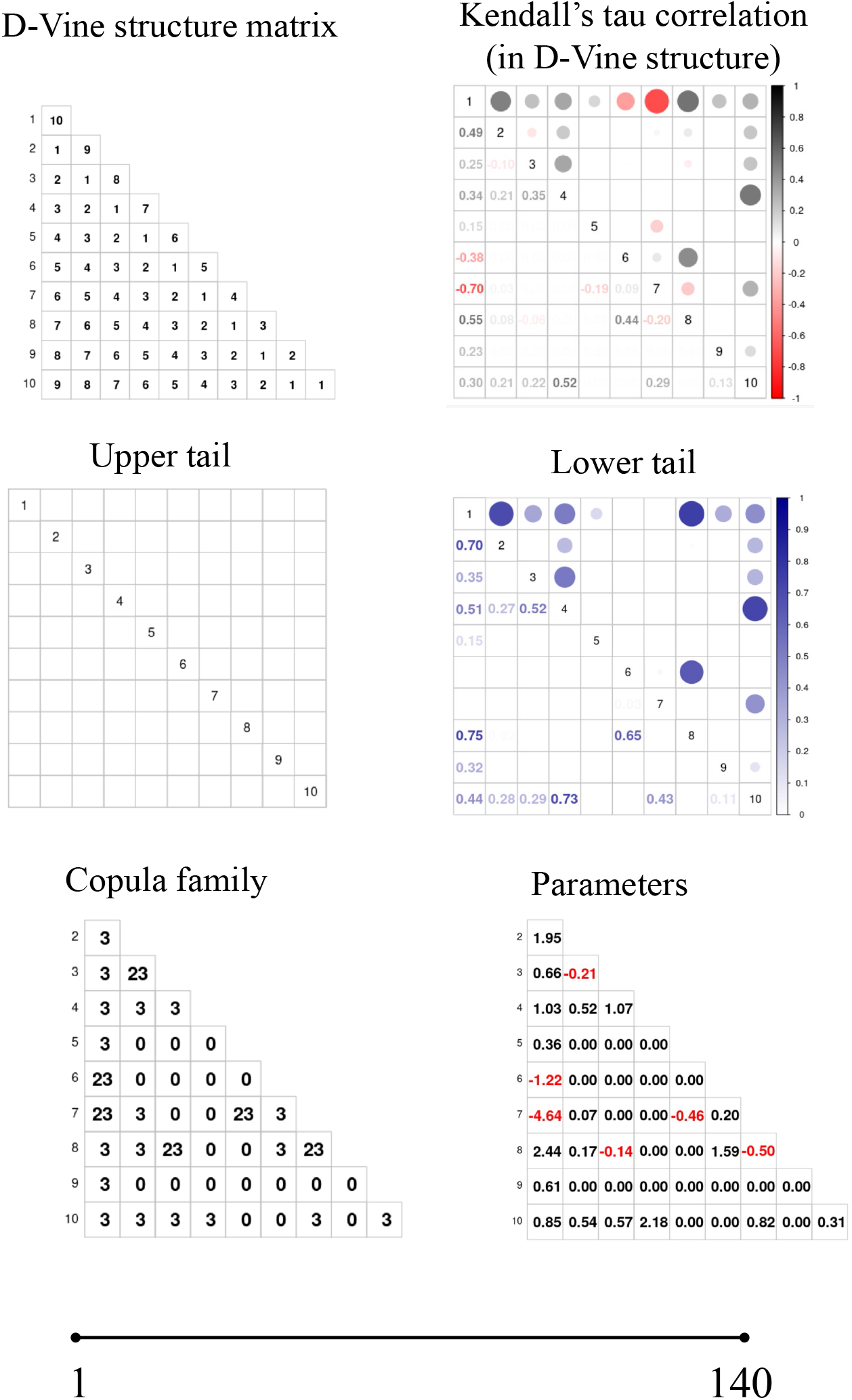
A visual display of Simulation 1 with no change point in the vine copula structure. The first row depicts the D-vine structure and corresponding Kendall’s *τ* across the entire time series. The second row depicts the upper and lower dependence structures. The third row depicts the copula type and the parameter values. For the copula type, 0 represents independence, 13 represents the clayton copula rotated 180 degrees, 23 represents the clayton copula rotated 90 degrees, 14 represents the gumbel copula rotated 180 degrees, and 24 represents the gumbel copula rotated 90 degrees.

Figure 1 in the Supplementary Materials provides a visual display of Simulation 2 (one change point in the vine copula structure). In this simulation, *P* = 10 and *T* = 140, with the change point occurring at time *t* = 99. The left column represents the structure for time points (1:99), while the right column represents the structure for time points (100:140). The change point coincides with a change in the tail dependency structure.

Figure 2 in the Supplementary Materials provides a visual display of Simulation 3 (two change points in the vine copula structure). In this simulation, *P* = 10 and *T* = 210, with the change points occurring at times *t* = 71, 141. The first, second and third columns represent the structure for time points (1:71), (72:141), and (142:210), respectively. The change points coincide with changes in the tail dependency structure.

Figure 3 in the Supplementary Materials provides a visual display of Simulation 4 (three change points in the vine copula structure). In this simulation, *P* = 10 and *T* = 280, with the change points occurring at times *t* = 71, 141, 211. The first, second, third, and fourth columns represent the structure for time points (1:71), (72:141), (142:210), and (211:280), respectively. The change points coincide with changes in the tail dependency structure, with an on-off-on-off type structure that is common in neuroimaging studies.

Unless stated otherwise, we have the following settings for the simulations. The minimum distance between change points is set to *δ* = 30. The tree structure is D-vine. The candidate bivariate copula models were Gaussian, *t*, Clayton, Gumbel, and Frank (see the Supplementary Materials for definitions). The maximum likelihood inference is numerically difficult when non-uniform marginal distributions are involved. In addition, our focus is on the network structure instead of marginal distributions. Therefore, we employ the rank transformations and normalization as follows before any vine copula construction to obtain marginally uniform data similar to Genest et al. (1995):

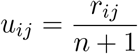

where *r_ij_* is the rank of *x_ij_* among *x_kj_, k* = 1, …, *n*. As long as the transformation is monotonic, it does not change Kendall’s *τ*, which we use to select the copula type.

The significance level of the Vuong test is set to *α* = 0.1, as two tests (left and right partitions) have to be rejected simultaneously before identifying a change point. The average block size and the number of resamples in the SB is 1/0.3 (or *p* = 0.3) and 100, respectively. For MOSUM, we set the bandwidth to *G* = 0.1 ∗ *T* and the number of random intervals to *M* = [9 ∗ log(*T*)] in WBS, where [*x*] denotes the maximal integer smaller than *x*.

We compare the True Positive rate (TP: the ability to detect the true change points), False Positive rate (FP: detecting incorrect change points) and False Negative rate (FN: failing to detect the true change points). As a measure of the accuracy of the detected locations in time compared to the location of the true change points, we also provide the scaled Hausdorff distance,

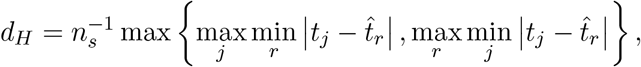

where *n_s_* is the length of the largest segment, 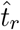 are the estimated change points and *t_j_* are the true change points. The optimal model obtains a minimum scaled Hausdorff distance. The bias standard we use is 10 time points, hence detected change points located no more than 10 units away from the true change points are considered a correct detection.

Many of the methods mentioned in the introduction assume Gaussianity and linear dependence for data between the change points, which is equivalent to using a Gaussian copula in our simulations and fMRI data sets. However, we also compare VCCP to 5 non-parametric methods, e.divisive (Matteson and James, 2014), e.cp3o, e.cp3o_delta, ks.cp3o and ks.cp30_delta (Zhang et al., 2017), implemented in the ecp package (James and Matteson, 2015). We ignore the e.agglo method since it requires an initial segmentation of the data. The five methods were built for multivariate time series data and multiple change points detection. For e.divisive, we keep *min.size* (the minimum number of observations between change points) parameter to be the default value 30, which is equivalent to *δ* = 30 in our VCCP model. The moment index *α* used for determining the distance between and within segments can be 1 or 2, and the number of change point locations to estimate is based on inference test in each turn. The e.cp3o and kp.cp3o methods use the E-statistic and the Kolmogorov-Smirnov statistic for the goodness-of-fit measure, respectively. Both of these methods require users to input *Kmax*, the maximum number of possible change points and *min.size*. E.cp3o_delta and ks.cp3o_delta narrow down the window size used to calculate the complete portion of the approximate test statistic, significantly reducing the computation time. Likewise, it requires *Kmax*, *δ* (window size) and *α*. In simulations 2, 3, 4 *Kmax* = 1, 2, *Kmax* = 2, 4, *Kmax* = 3, 6, respectively.

### 3.2 fMRI data

We also apply our new VCCP methodology to two task-based fMRI data sets. The first fMRI data set was acquired from 9 right-handed native English speakers (5 females and 4 males) aged 18–40 years, while they read Chapter 9 of *Harry Potter and the Sorcerer’s Stone* (one subject’s data was discarded due to artifacts). All subjects had read the Harry Potter book series, or seen the movie series prior to participating in the experiment. All the subjects therefore were familiar with the characters and the events of the book, and were reminded of the events leading up to Chapter 9 before the experiment. This chapter was chosen because it involves many characters and spans multiple locations and scenes. The chapter was read using Rapid Serial Visual Presentation (RSVP): the words of the chapter were presented one by one in the center of the screen, for 0.5 s each. The sampling rate of fMRI acquisition was 2 seconds per observation, hence, four consecutive words were read during the time it took to scan the whole brain once. The 45-minute experiment was divided into four runs, each starting with 20 seconds (= 10 TRs) of rest, and ending with 10 seconds (= 5 TRs) of rest. After discarding data collected during the resting periods, we obtained 1290 scans, during which approximately 5200 words were presented. Figure 5 shows the main plotline within each run. Data was preprocessed as in Wehbe et al. (2014). Fourteen ROIs defined using the Automated Anatomical Atlas (AAL: Tzourio-Mazoyer et al., 2002) were extracted from the data set (Table 2 in the Supplementary Materials). These regions contain a variety of voxels that can distinguish between the literary content of two novel text passages based on neural activity while these passages are being read and are important for understanding reading processes (Wehbe et al., 2014). In this work, we focus on exploring whether the FC network of the 14 ROIs changes as the subjects read. We focus on the varying textual features about story characters (e.g., emotion, motion and dialog) that the dynamic networks encode.

**Figure 5:**
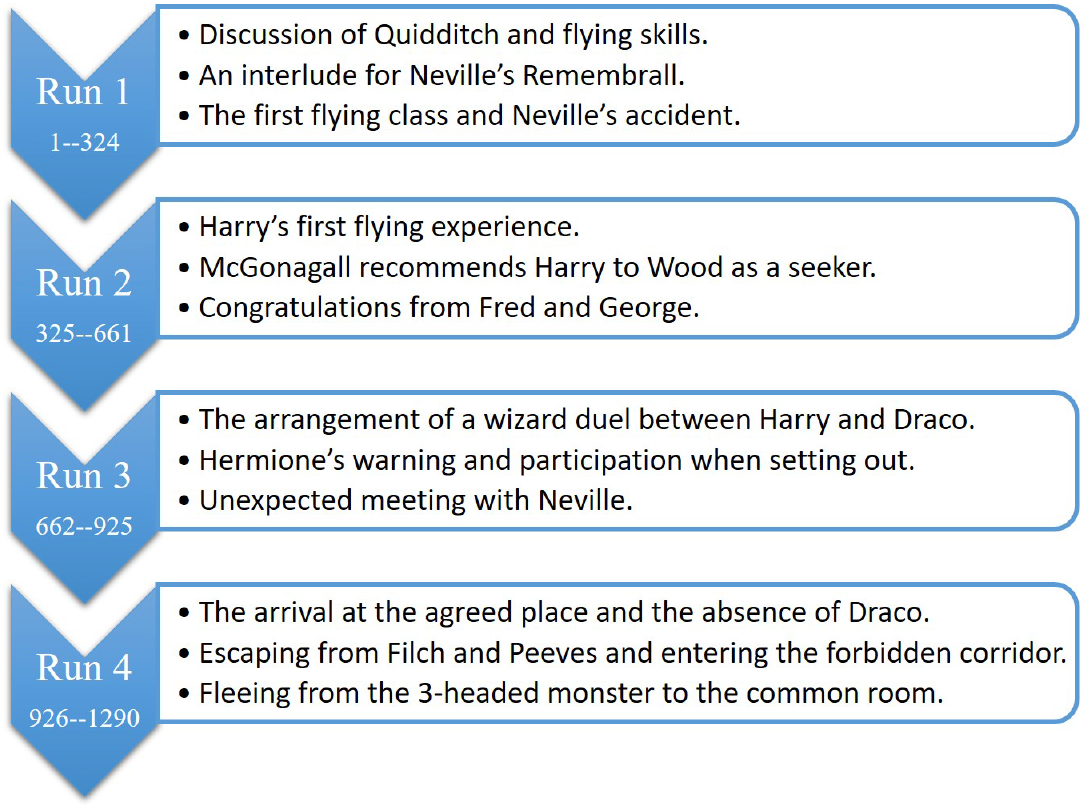
The main plot lines within each run of Chapter 9 of *Harry Potter and the Sorcerer’s Stone*.

**Table 2:** The results from applying all variations of VCCP (a) with 5 different copula types (Gaussian, *t*, Clayton, Gumbel and Frank copulas) and (b) with the Gaussian copula only to Simulation 1 (no change points) over 100 simulated data sequences. The first 8 r ows d enote t he a verage c hange p oints detected in various time intervals. ‘Cand.num’ is the averaged number of candidates while ‘Num.det’ refers to the average number of detected change points after the inferential step. N, O, M, and W denote the adapted binary segmentation, old binary segmentation, moving sum, and the wild binary segmentation methods, respectively. ‘5’ denotes the 5 family copula types; ‘1’ denotes the Gaussian copula; ‘BS’ and ‘V’ denote the stationary bootstrap and the Vuong test.

|  | N.5.BS | N.5.V | O.5.BS | O.5.V | M.5.BS | M.5.V | W.5.BS | W.5.V |
| --- | --- | --- | --- | --- | --- | --- | --- | --- |
| 31-60 | 0.04 | 0.02 | 0.04 | 0.02 | 0.02 | 0.01 | 0.12 | 0.02 |
| 61-90 | 0.02 | 0.02 | 0.02 | 0.02 | 0 | 0.01 | 0.19 | 0.06 |
| 91-120 | 0 | 0 | 0 | 0 | 0.01 | 0 | 0.27 | 0.06 |
| 121-150 | 0.01 | 0 | 0.01 | 0 | 0.01 | 0 | 0.22 | 0.1 |
| 151-180 | 0.01 | 0 | 0.01 | 0 | 0.01 | 0.01 | 0.22 | 0.09 |
| 181-210 | 0 | 0 | 0 | 0 | 0.01 | 0.01 | 0.23 | 0.1 |
| 211-240 | 0.02 | 0.02 | 0.02 | 0.02 | 0 | 0 | 0.15 | 0.04 |
| 241-270 | 0.08 | 0.03 | 0.08 | 0.03 | 0.01 | 0 | 0.16 | 0.05 |
| Cand.num | 0.18 | 0.18 | 0.18 | 0.18 | 2.92 | 2.92 | 1.82 | 1.32 |
| Num.det | 0.18 | 0.09 | 0.18 | 0.09 | 0.07 | 0.04 | 1.58 | 0.53 |
| time (s) | 328.37 | 306.6 | 328.79 | 290.98 | 303.48 | 86.55 | 2171.21 | 2075.89 |

(b)
|  | N.1.BS | N.1.VD | O.1.BS | O.1.VD | M.1.BS | M.1.VD | W.1.BS | W.1.VD |
| --- | --- | --- | --- | --- | --- | --- | --- | --- |
| 31-60 | 0.01 | 0.01 | 0.01 | 0.01 | 0.04 | 0.01 | 0.11 | 0 |
| 61-90 | 0 | 0 | 0 | 0 | 0.06 | 0.03 | 0.2 | 0.01 |
| 91-120 | 0 | 0 | 0 | 0 | 0.02 | 0.03 | 0.2 | 0.01 |
| 121-150 | 0 | 0 | 0 | 0 | 0.05 | 0.02 | 0.14 | 0.01 |
| 151-180 | 0 | 0 | 0 | 0 | 0.03 | 0.01 | 0.17 | 0 |
| 181-210 | 0 | 0 | 0 | 0 | 0.05 | 0.03 | 0.22 | 0.03 |
| 211-240 | 0.02 | 0.02 | 0.02 | 0.02 | 0.03 | 0.03 | 0.21 | 0 |
| 241-270 | 0 | 0 | 0 | 0 | 0.05 | 0.05 | 0.11 | 0 |
| Cand.num | 0.03 | 0.03 | 0.03 | 0.03 | 3.01 | 3.01 | 1.68 | 0.96 |
| Num.det | 0.03 | 0.03 | 0.03 | 0.03 | 0.33 | 0.21 | 1.37 | 0.06 |
| time (s) | 91.58 | 90.46 | 90.88 | 89.87 | 143.36 | 49.79 | 846.91 | 808.29 |

A description of the second fMRI data set (SET) can be found in the Supplementary Materials. For both fMRI data sets, we used the same settings as the simulations (Section 3.1), except for the first fMRI data set, we set *δ* = 40. In addition, since the experiment lasted 45 minutes, and given the computational burden for BS, we run VCCP on each of the four runs separately.

## 4 Results

### 4.1 Simulations

The results for the multivariate normal (MVN) data and the Vector Autoregression (VAR) simulations are displayed in the Supplementary Materials. We now describe the results from the non-Gaussian data. In general, we compare our VCCP model to the two best performing ecp methods in the following tables.

Table 2(a) shows the results from applying all variations of VCCP with 5 different copula types to Simulation 1 (no change points) over 100 simulated data sequences. For comparison purposes, we compare the results to all variations of VCCP with only the Gaussian copula (Table 2(b)). Using only the Gaussian copula, the size of the test is closer to the nominal rate. For VCCP with 5 different copula types, MOSUM in combination with the Voung test (MOSUM.V) has the best results. OBS and NBS with the Voung test are next best. WBS with the SB appears too liberal, it detects more than one change point. In general, the Vuong test with all the segmentation methods is more conservative than the SB and closer to the nominal rate.

Table 3(a) shows the results from applying all variations of VCCP with 5 different copula types to Simulation 2 (one change point) over 100 simulated data sequences. For comparison purposes, we compare the results to all variations of VCCP with only the Gaussian copula (Table 3(b)). As expected, given the change occurs in the tail dependency structure, VCCP with 5 different copula types outperforms the VCCP with only the Gaussian copula. For VCCP with 5 different copula types, NBS, OBS and WBS detects the change point with TP rates larger than 0.9. Overall, NBS.SB stands out for its low FP and high TP rates. It also has the lowest Hausdorff distance. Overall, the binary segmentation methods outperform MOSUM and WBS in this simulation. We also compared our VCCP model to all the non-parametric methods in the ecp R package but only show the two best performing methods in Table 3(a): ecp.1 and ecp.2 denote the e.cp3o_delta_2K and ks.cp3o_delta_2K methods, respectively, with ecp.1 having the highest TP rate and ecp.2 detecting the most accurate number of change points. Both methods, while computationally fast, provide low TP rates and high FP rates.

**Table 3:** The results from applying all variations of VCCP (a) with 5 different copula types (Gaussian, *t*, Clayton, Gumbel and Frank copulas) and (b) with the Gaussian copula only to Simulation 2 (one change point) over 100 simulated data sequences. TP, FN, FP, Cand.num, Num.det, and *dH* denote the true positive rate, false negative rate, false positive rate, number of candidate change points, number of change points detected, scaled Hausdorff distance, respectively. N, O, M, and W denote the adapted binary segmentation, old binary segmentation, moving sum, and the wild binary segmentation methods, respectively. ‘5’ denotes the 5 family copula types; ‘1’ denotes the Gaussian copula; ‘BS’ and ‘V’ denote the stationary bootstrap and the Vuong test. ecp.1 and ecp.2 refer to two best performing methods from the ecp R package.

| (a) |  |  |  |  |  |  |  |  |  |  |
| --- | --- | --- | --- | --- | --- | --- | --- | --- | --- | --- |
|  | N.5.BS | N.5.V | O.5.BS | O.5.V | M.5.BS | M.5.V | W.5.BS | W.5.V | ecp.1 | ecp.2 |
| TP_98 | 0.91 | 0.77 | 0.91 | 0.76 | 0.57 | 0.39 | 0.95 | 0.83 | 0.25 | 0.29 |
| FN_98 | 0 | 0.14 | 0 | 0.15 | 0.35 | 0.52 | 0.01 | 0.13 | 0.41 | 0.33 |
| FP_1-97 | 0.15 | 0.08 | 0.15 | 0.08 | 0.13 | 0.06 | 0.29 | 0.11 | 0.74 | 0.72 |
| FP_98-139 | 0.05 | 0.04 | 0.05 | 0.04 | 0.02 | 0.02 | 0.02 | 0.02 | 0.03 | 0 |
| Hausdorff | 0.16 | 0.33 | 0.16 | 0.34 | 0.73 | 0.98 | 0.27 | 0.34 | 0.50 | 0.56 |
| Cand.num | 1.07 | 1.07 | 1.07 | 1.04 | 1.71 | 1.71 | 1.34 | 1.33 | 2.53 | 2.09 |
| Num.det | 1.06 | 0.85 | 1.06 | 0.84 | 0.71 | 0.45 | 1.24 | 0.94 | 1 | 1 |
| time (s) | 144.5 | 72.22 | 145.81 | 68.81 | 127.1 | 29.72 | 273.74 | 211.68 | 0.01 | 0.00 |

| (b) |  |  |  |  |  |  |  |  |
| --- | --- | --- | --- | --- | --- | --- | --- | --- |
|  | N.1.BS | N.1.V | O.1.BS | O.1.V | M.1.BS | M.1.V | W.1.BS | W.1.V |
| TP_98 | 0.12 | 0.04 | 0.12 | 0.04 | 0.13 | 0.04 | 0.26 | 0.16 |
| FN_98 | 0 | 0.08 | 0 | 0.08 | 0.33 | 0.4 | 0.02 | 0.1 |
| FP_1-97 | 0.15 | 0.12 | 0.15 | 0.12 | 0.22 | 0.09 | 0.48 | 0.13 |
| FP_98-139 | 0.01 | 0 | 0.01 | 0 | 0.01 | 0 | 0.01 | 0 |
| Hausdorff | 1.32 | 1.48 | 1.32 | 1.48 | 1.27 | 1.47 | 0.77 | 1.23 |
| Cand.num | 0.28 | 0.28 | 0.28 | 0.27 | 1.48 | 1.37 | 0.87 | 0.81 |
| Num.det | 0.28 | 0.15 | 0.28 | 0.15 | 0.37 | 0.13 | 0.78 | 0.29 |
| time (s) | 36.6 | 27.56 | 36.02 | 25.84 | 62.14 | 17.51 | 119.78 | 100.01 |

Table 4(a) shows the results from applying all variations of VCCP with 5 different copula types to Simulation 3 (two change points) over 100 simulated data sequences. For comparison purposes, we compare the results to all variations of VCCP with only the Gaussian copula (Table 4(b)). Similar to Simulation 2, given the changes occur in the tail dependency structure, VCCP with 5 different copula types outperforms the VCCP with only the Gaussian copula. None of the VCCP variations with the Gaussian copula were able to identify the two change points with a TP rate higher than 0.25 except the WBS.BS combination. For VCCP with 5 different copula types, the best method is WBS.BS which provides high TP rates, low FP rates and small Hausdorff distances results for this multiple change point simulation. However, NBS.BS, OBS.BS and WBS.V also perform well in terms of high TP rates, low FP rates and small Hausdorff distances. In terms of computation time, MOSUM in combination with the Voung test performs best and has low FP rates, however, we do not recommend it due to its over-conservative detection (high FN rates). The two best performing non-parametric methods from the ecp R package were e.cp3o_delta_2K (lowest Hausdorff distance) and ks.cp3o_delta_2K (most accurate number of change points detected). Both methods, while computationally fast, provide low TP rates and very high FP (e.cp3o_delta_2K had a FP rate of 0.68 for one interval).

**Table 4:** The results from applying all variations of VCCP (a) with 5 different copula types (Gaussian, *t*, Clayton, Gumbel and Frank copulas) and (b) with the Gaussian copula only to Simulation 3 (two change points) over 100 simulated data sequences. TP, FN, FP, Cand.num, Num.det, and *dH* denote the true positive rate, false negative rate, false positive rate, number of candidate change points, number of change points detected, scaled Hausdorff distance, respectively. N, O, M, and W denote the adapted binary segmentation, old binary segmentation, moving sum, and the wild binary segmentation methods, respectively. ‘5’ denotes the 5 family copula types; ‘1’ denotes the Gaussian copula; ‘BS’ and ‘V’ denote the stationary bootstrap and the Vuong test. ecp.1 and ecp.2 refer to two best performing methods from the ecp R package.

| (a) |  |  |  |  |  |  |  |  |  |  |
| --- | --- | --- | --- | --- | --- | --- | --- | --- | --- | --- |
|  | N.5.BS | N.5.V | O.5.BS | O.5.V | M.5.BS | M.5.V | W.5.BS | W.5.V | ecp.1 | ecp.2 |
| TP_71 | 0.54 | 0.38 | 0.52 | 0.25 | 0.27 | 0.1 | 0.77 | 0.51 | 0.35 | 0.26 |
| TP_141 | 0.73 | 0.66 | 0.72 | 0.45 | 0.55 | 0.33 | 0.9 | 0.81 | 0.4 | 0.21 |
| FN_71 | 0 | 0.16 | 0 | 0.08 | 0.23 | 0.39 | 0 | 0.19 | 0.06 | 0.08 |
| FN_141 | 0.01 | 0.08 | 0.01 | 0.24 | 0.1 | 0.29 | 0 | 0.08 | 0.12 | 0.13 |
| FP_1-70 | 0.17 | 0.09 | 0.18 | 0.05 | 0.12 | 0.03 | 0.13 | 0.08 | 0.6 | 0.52 |
| FP_71-140 | 0.15 | 0.07 | 0.15 | 0.02 | 0.12 | 0.06 | 0.13 | 0.09 | 0.68 | 0.66 |
| FP_141-209 | 0.25 | 0.19 | 0.25 | 0.13 | 0.24 | 0.12 | 0.09 | 0.03 | 0.55 | 0.52 |
| Hausdorff | 0.6 | 0.93 | 0.62 | 1.4 | 1.12 | 1.66 | 0.24 | 0.66 | 0.45 | 1.01 |
| Cand.num | 1.91 | 1.91 | 1.88 | 1.42 | 2.78 | 2.54 | 2.03 | 1.92 | 6.04 | 4.04 |
| Num.det | 1.82 | 1.37 | 1.8 | 0.88 | 1.29 | 0.64 | 1.96 | 1.52 | 2.52 | 2.42 |
| time (s) | 335.32 | 206.54 | 354.31 | 173.32 | 219.42 | 54.94 | 875.55 | 772.78 | 0.06 | 0.01 |
| (b) |  |  |  |  |  |  |  |  |  |  |
|  | N.1.BS | N.1.V | O.1.BS | O.1.V | M.1.BS | M.1.V | W.1.BS | W.1.V |  |  |
| TP_71 | 0.16 | 0.08 | 0.16 | 0.04 | 0.25 | 0.04 | 0.65 | 0.23 |  |  |
| TP_141 | 0.01 | 0.01 | 0.01 | 0 | 0.07 | 0.02 | 0.27 | 0.12 |  |  |
| FN_71 | 0 | 0.08 | 0 | 0.11 | 0.21 | 0.36 | 0.01 | 0.37 |  |  |
| FN_141 | 0 | 0 | 0 | 0.01 | 0.24 | 0.27 | 0.05 | 0.08 |  |  |
| FP_1-70 | 0.07 | 0 | 0.07 | 0 | 0.13 | 0.03 | 0.08 | 0.02 |  |  |
| FP_71-140 | 0.06 | 0.04 | 0.06 | 0.04 | 0.15 | 0.04 | 0.33 | 0.06 |  |  |
| FP_141-209 | 0.03 | 0.01 | 0.03 | 0.01 | 0.05 | 0.03 | 0.23 | 0.04 |  |  |
| Hausdorff | 2.01 | 2.19 | 2.01 | 2.25 | 1.76 | 2.19 | 0.75 | 1.8 |  |  |
| Cand.num | 0.33 | 0.33 | 0.33 | 0.29 | 2.23 | 1.96 | 1.63 | 1.25 |  |  |
| Num.det | 0.33 | 0.14 | 0.33 | 0.09 | 0.65 | 0.16 | 1.51 | 0.47 |  |  |
| time (s) | 73.57 | 62.01 | 73.05 | 55.09 | 102.45 | 32.69 | 410.29 | 371.8 |  |  |

Table 5(a) shows the results from applying all variations of VCCP with 5 different copula types to Simulation 4 (three change points) over 100 simulated data sequences. The change points coincide with changes in the tail dependency structure, with an on-off-on-off type structure that is common in neuroimaging studies. For comparison purposes, we compare the results to all variations of VCCP with only the Gaussian copula (Table 5(b)). Similar to Simulations 2 and 3, but even more dramatic, VCCP with 5 different copula types outperforms the VCCP with only the Gaussian copula. OBS and NBS in combination with only the Gaussian copula detects almost no change points, MOSUM has low TP and high FN rates, while WBS has low TP and high FP rates. For VCCP with 5 different copula types, the NBS.BS, NBS.V, WBS.V, and WBS.BS combinations provide high TP rates, low FP rates and small Hausdorff distances results for the multiple change point simulation. In terms of computation time, MOSUM in combination with the Voung test performs best and has low FP rates, however, we do not recommend it due to its over-conservative detection (high FN rates). WBS has the largest computation time in terms of segmentation, while the SB is computationally more intense then the Vuong test. In contrast, the TP rates of the two best non-parametric methods (e.cp3o_delta_2K had the lowest Hausdorff distance while ks.cp3o_2K had the most accurate number of change points detected) from the ecp R package are poor with high FP rates (e.cp3o_delta_2K had a 0.91 FP rate).

**Table 5:** The results from applying all variations of VCCP (a) with 5 different copula types (Gaussian, *t*, Clayton, Gumbel and Frank copulas) and (b) with the Gaussian copula only to Simulation 4 (three change points) over 100 simulated data sequences. TP, FN, FP, Cand.num, Num.det, and *dH* denote the true positive rate, false negative rate, false positive rate, number of candidate change points, number of change points detected, scaled Hausdorff distance, respectively. N, O, M, and W denote the adapted binary segmentation, old binary segmentation, moving sum, and the wild binary segmentation methods, respectively. ‘5’ denotes the 5 family copula types;‘1’ denotes the Gaussian copula; ‘BS’ and ‘V’ denote the stationary bootstrap and the Vuong test. ecp.1 and ecp.2 refer to two best performing methods from the ecp R package.

| (a) |  |  |  |  |  |  |  |  |  |  |
| --- | --- | --- | --- | --- | --- | --- | --- | --- | --- | --- |
|  | N.5.BS | N.5.V | O.5.BS | O.5.V | M.5.BS | M.5.V | W.5.BS | W.5.V | ecp.1 | ecp.2 |
| TP_71 | 0.81 | 0.73 | 0.82 | 0.24 | 0.55 | 0.34 | 0.86 | 0.66 | 0.16 | 0.45 |
| TP_141 | 0.86 | 0.84 | 0.86 | 0.13 | 0.75 | 0.52 | 0.96 | 0.94 | 0.26 | 0.23 |
| TP_211 | 0.77 | 0.66 | 0.78 | 0.18 | 0.52 | 0.31 | 0.83 | 0.67 | 0.17 | 0.09 |
| FN_71 | 0.01 | 0.09 | 0 | 0.22 | 0.1 | 0.3 | 0 | 0.14 | 0.17 | 0.3 |
| FN_141 | 0 | 0.02 | 0 | 0.13 | 0.04 | 0.27 | 0 | 0.02 | 0.27 | 0.35 |
| FN_211 | 0.03 | 0.14 | 0.02 | 0.29 | 0.11 | 0.32 | 0 | 0.1 | 0.19 | 0.16 |
| FP_1-70 | 0.12 | 0.05 | 0.14 | 0.03 | 0.13 | 0.06 | 0.09 | 0.03 | 0.54 | 0.68 |
| FP_71-140 | 0.1 | 0.05 | 0.14 | 0 | 0.16 | 0.08 | 0.06 | 0.04 | 0.91 | 0.67 |
| FP_141-210 | 0.2 | 0.14 | 0.2 | 0 | 0.22 | 0.11 | 0.05 | 0.02 | 0.89 | 0.31 |
| FP_211-279 | 0.17 | 0.09 | 0.17 | 0.02 | 0.17 | 0.13 | 0.05 | 0.02 | 0.44 | 0.1 |
| Hausdorff | 0.5 | 0.75 | 0.49 | 2.84 | 0.85 | 1.67 | 0.25 | 0.68 | 0.71 | 2.26 |
| Cand.num | 3.2 | 3.2 | 3.19 | 1.47 | 3.86 | 3.81 | 2.94 | 2.79 | 8.4 | 7.14 |
| Num.det | 2.98 | 2.52 | 3.06 | 0.6 | 2.49 | 1.54 | 2.9 | 2.38 | 3.46 | 3.33 |
| time (s) | 661.14 | 457.27 | 738.09 | 302.06 | 313.19 | 80.33 | 1787.47 | 1641.97 | 0.1 | 0.89 |
| (b) |  |  |  |  |  |  |  |  |  |  |
|  | N.1.BS | N.1.V | O.1.BS | O.1.V | M.1.BS | M.1.V | W.1.BS | W.1.V |  |  |
| TP_71 | 0 | 0 | 0 | 0 | 0.04 | 0.02 | 0.27 | 0.08 |  |  |
| TP_141 | 0 | 0 | 0 | 0 | 0.02 | 0.02 | 0.16 | 0.05 |  |  |
| TP_211 | 0.01 | 0.01 | 0.01 | 0.01 | 0.06 | 0.03 | 0.23 | 0.09 |  |  |
| FN_71 | 0 | 0 | 0 | 0 | 0.23 | 0.25 | 0.01 | 0.09 |  |  |
| FN_141 | 0 | 0 | 0 | 0 | 0.21 | 0.17 | 0.03 | 0.04 |  |  |
| FN_211 | 0 | 0 | 0 | 0 | 0.28 | 0.3 | 0.02 | 0.11 |  |  |
| FP_1-70 | 0.03 | 0 | 0.03 | 0 | 0.02 | 0.02 | 0.08 | 0.02 |  |  |
| FP_71-140 | 0.01 | 0.01 | 0.01 | 0.01 | 0.14 | 0.07 | 0.34 | 0.09 |  |  |
| FP_141-210 | 0 | 0 | 0 | 0 | 0.1 | 0.04 | 0.39 | 0.1 |  |  |
| FP_211-279 | 0.01 | 0 | 0.01 | 0 | 0.05 | 0.02 | 0.19 | 0.03 |  |  |
| Hausdorff | 3.46 | 3.49 | 3.46 | 3.49 | 3 | 3.19 | 1.48 | 2.9 |  |  |
| Cand.num | 0.06 | 0.06 | 0.06 | 0.06 | 2.88 | 2.78 | 1.82 | 1.25 |  |  |
| Num.det | 0.06 | 0.02 | 0.06 | 0.02 | 0.42 | 0.23 | 1.65 | 0.46 |  |  |
| time (s) | 89.3 | 87.02 | 88.48 | 83.85 | 139.28 | 47.06 | 778.91 | 735.63 |  |  |

### 4.2 Harry Potter task-based fMRI data

In order to show that other forms of dependence exist beyond linear dependence for fMRI data, we compare the detected change points found by the NBS.V combination of VCCP using only the Gauss copula to NBS.V using 5 types of copulas (Gauss, t, Clayton, Gumbel, and Frank copula). Figure 6 shows the detected change points. As expected, by considering various forms of dependence and copulas beyond Gaussian, more change points are detected (1/3 more detected). For example, let’s consider the detected change points for subject 7. Using NBS.V with the Gaussian copula, time points (*p*-values in parenthesis) *t* = 43 (*<* .001), 235 (< .001), 282 (< .001), 372 (0.002), 436 (0.003), 704 (0.003), 1003 (< .001), 1099 (*<* .001), 1149 (0.003), 1247 (< .001) are identified as change points, while time points *t* = 44 (< .001), 241 (< .001), 282 (< .001), 390 (< .001), 704 (0.003), 746 (< .001), 792 (*<* .001), 867 (0.002), 998 (< .001), 1038 (.005), 1096 (< .001), 1168 (< .001), 1247 (.001), are identified as change points for NBS.V with the 5-copula family. Now let’s focus our attention on change points at *t* = 998, 1038 and *t* = 1096. The first and last change points are detected by both methods while the middle change point is missed by NBS.V with the Gaussian copula. Figure 7 displays the estimated Kendall’s *τ*, the upper tail dependence, the lower tail dependence, and the optimal copula function between pairs of ROIs for data between change points. From segment 867 − 997 to segment 998 − 1037, the Kendall’s *τ* network becomes more sparse, while the lower tail network (especially within the IFG sub-network) and the upper tail network become more dense. As a result, the dominant copula type turns out to be the Clayton copula during the second segment (998 − 1037). As the Gaussian copula model is unable to model tail and other forms of dependence, the change point at *t* = 998 is detected only because of the change in Kendall’s *τ*. At time point *t* = 1038, there is not an obvious strong discernible change in Kendall’s *τ*, hence, the Gaussian copula fails to find the related change point. Only models free of the Gaussian assumption are capable of discovering this change point. NBS.V with the 5-copula family has this flexibility. From segment 1038 − 1095 to segment 1096 − 1167, negative edges between nodes IT, IFG1, IFG3 and MT switch to positive, while a new sub-network between F, SM, PCG and IT appears with most of its edges being negative in the network of Kendall’s *τ*. The density of edges also alters. In addition, the lower tail dependence weakens so that only edges between TP and other nodes (TP becomes a hub node) remain, which results in the Clayton copula not being the dominant copula type. Furthermore, there is little change in the upper tail dependence network. Hence, both NBS.V with the Gaussian copula and the 5-copula family are able to detect the change point at *t* = 1096. To understand the rationale behind the change point at *t* = 1038, the last sentence given to readers before this time is, ‘They ripped through a tapestry and found themselves in a hidden passageway, … which they knew was miles from the trophy room.’, and the first sentence after *t* = 1038 is ‘ “I think we’ve lost him,” Harry panted, leaning against the cold wall and wiping his forehead.’ This passage occurs when Harry and the other three characters run away from Filch. The text during this period includes changes in locations, actions and emotions.

**Figure 6:**
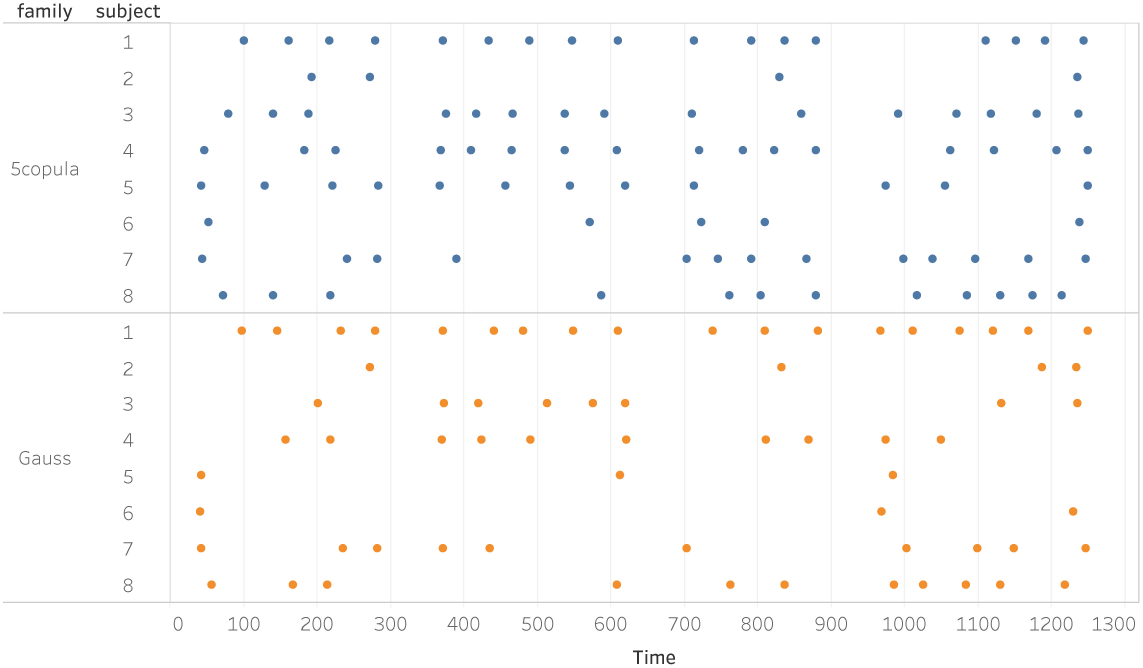
The top and bottom panels represent the change points from NBS.V using the 5-copula family and only the Gaussian copula, respectively, from the Harry Potter task-based fMRI study.

**Figure 7:**
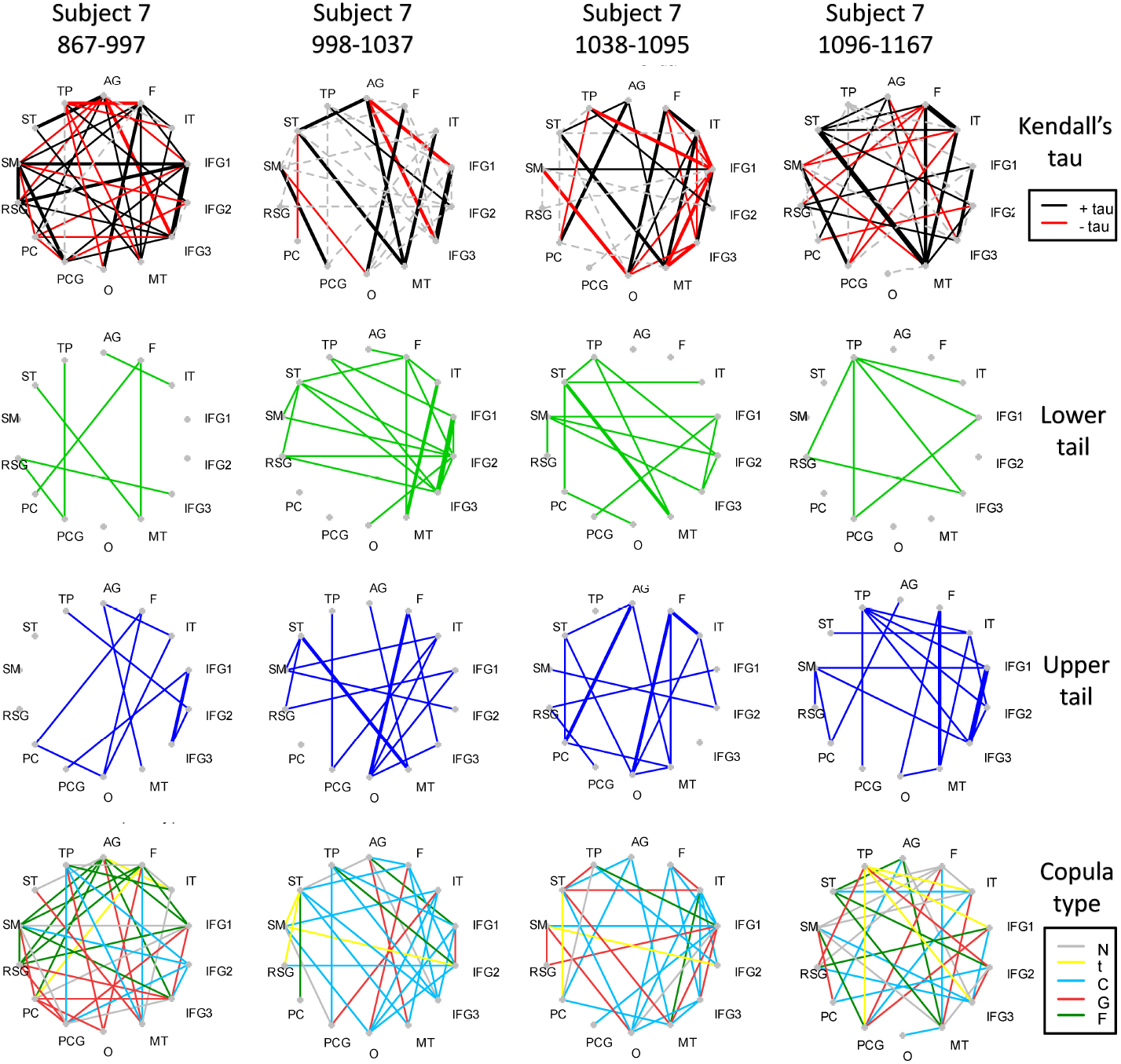
The connectivity graphs across 4 intervals, time points 867 997, 998 1037, 1038 1095, and 1096 1167 (the columns), for subject 7 in the Harry Potter task-based fMRI data set using the combination NBS.V of VCCP. The first, second, third and fourth rows represent the estimated Kendall’s *τ*, the lower tail dependence, the upper tail dependence, and the optimal copula function, for each edge between pairs of nodes. Black (red) lines in the graph on the first row represent positive (negative) Kendall’s *τ* correlation coefficients. Dashed lines indicate edges between nodes with tail dependence but statistically insignificant Kendall’s *τ*. Green and blue edges in the graphs on the second and the third rows represent the lower and upper tail dependence, respectively. The various colored lines on the bottom row indicate the optimal copula family for edges, with grey, yellow, blue, red and green indicating the Gaussian, *t*, Clayton, Gumbel and Frank copula, respectively.

For a complete understanding of the changing brain networks when reading, further exploration of the simultaneous changes in the text becomes necessary. Taking advantage of the rich text information available, we can assign attributes to every word in the text. In particular, we focus on whether the word is a character’s name (Char), implies a kind of emotion (Emo), refers to a motion (Mo), or contains a specific verb (Verb). Subdivisions of the four main word types, with their frequency among the 5200 words in the text, are shown in Table 6. As the ‘Motion’ attribute takes place over the course of a sentence, Wehbe et al. (2014) created two features: a punctual feature and a “sticky” feature. The punctual feature coincides when the verb of the motion is mentioned, and the sticky feature is active for the duration of the motion (i.e., the sentence). The same applies to the ‘Emotion’ attribute, where the punctual features coincide when the emotion is explicitly mentioned, and sticky features align is active for the duration of the emotion felt. Here, we concentrate on the sticky features and changes in them. First, we specify an interval parameter (*Int*) indicating the range in which text changes take place, for example, [*t* − *Int, t* + *Int* − 1]. The range covers two whole sentences to display the complete textual background of the change. Since the upper quantile of the sentence length equals 17 words and 4 words are displayed within 1 TR (2 sec), *Int* is equal to 4 TR. We define three types of changes in text features: ‘off-on’, ‘on-off’, and ‘mixed’ patterns. If the identifier of a feature remains constant on one side of a point *t* (e.g., [*t* − *Int, t* − 1]), but contains a heterogeneous element on the other side (e.g., [*t, t* + *Int* − 1]), we then call it a ‘off-on’ ([0 → 0,1] or [0,1 → 1]) or ‘on-off’ ([1 → 1,0] or [1,0 → 0]) change type. If both sides include heterogeneous elements (i.e., [0,1 → 0,1]), we call it a mixed type.

**Table 6:**
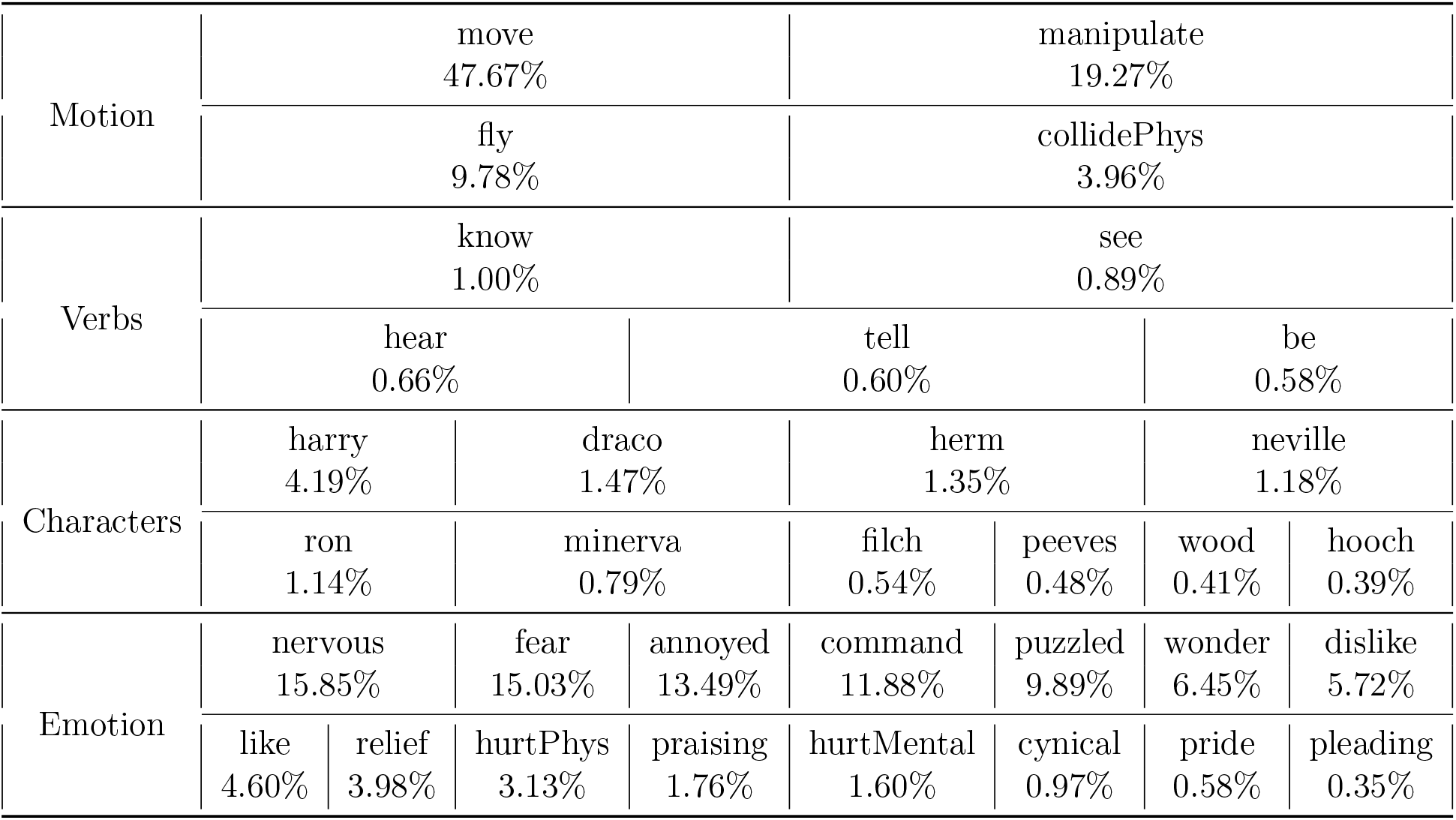
The frequency of words for the following four attributes: a character’s name (Char), emotion (Emo), motion (Mo), or a specific verb (Verb) in Chapter 9 of *Harry Potter and the Sorcerer’s Stone*.

To understand the difficulty of our analysis; coinciding changes in brain networks and the variation of a specific text feature, we explore the latent text changes, and in particular we consider the features before and after a particular time point in the text, time point *t* = 432. Figure 8 shows the text changes: three motion features (collide, fly, manipulate) experience ‘on-off’ changes; one verb (move), one emotion (nervous), and one character (Minerva) encounter ‘off-on’ patterns; and a character (Harry) feature has a mixed variation. Hence, for the FC change point at this time point, the corresponding text may involve changes in multiple features, which increases the difficulty of assigning the change points detected to one specific text feature.

**Figure 8:**
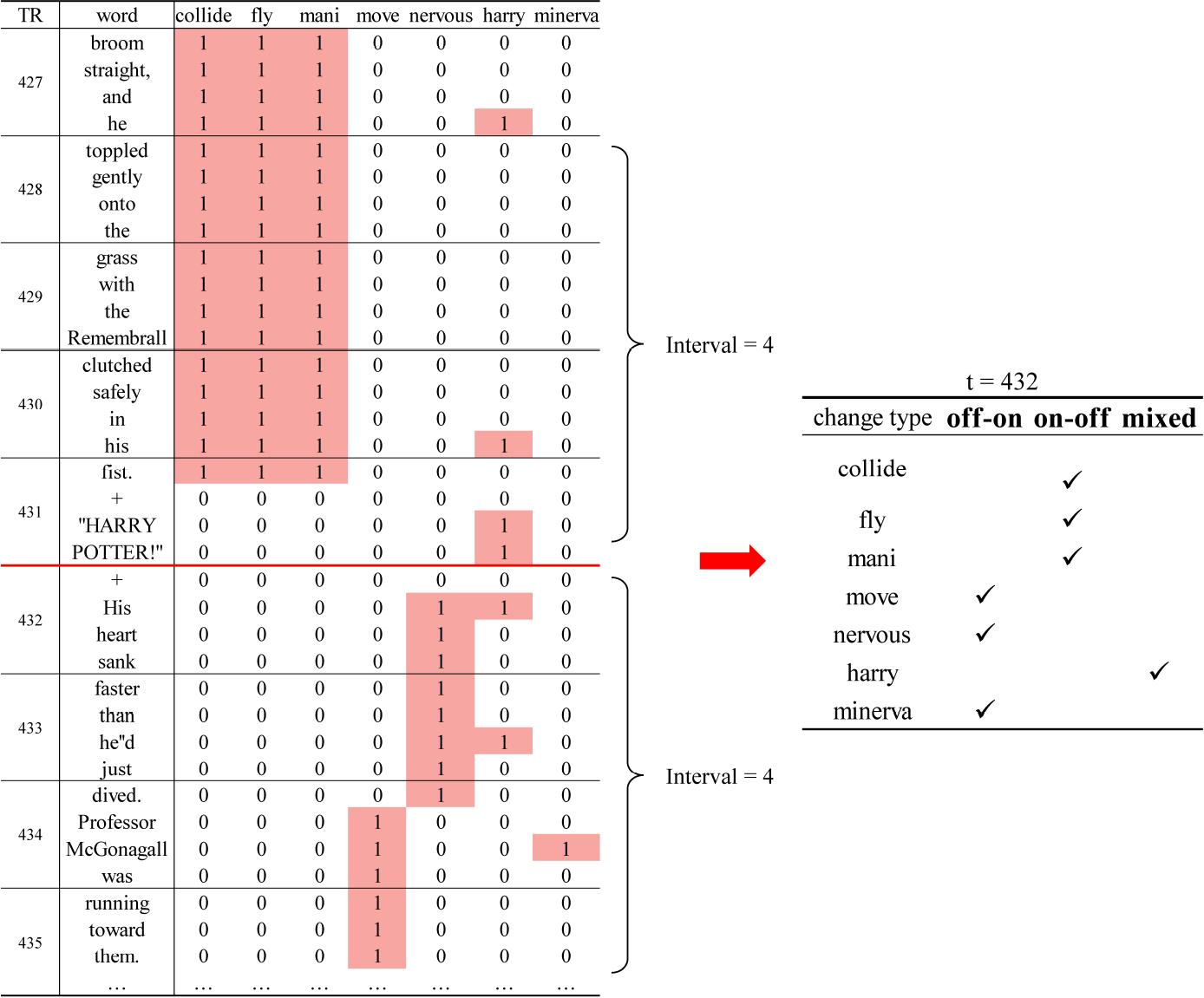
An example of three types of text changes at time point *t* = 432 in Chapter 9 of *Harry Potter and the Sorcerer’s Stone*, with the interval parameter fixed at 4 TRs.

In Figure 9 (top panel), we sum the frequency of attributes (Char, Emo, Mo, Verb) that correspond to change points detected by NBS.V with the 5-copula family for each subject. For example, the 94% in the first cell implies that amongst all the change points for subject 1, 94% are located at time points that coincide with character switching. Overall, there is a great deal of subject-level heterogeneity. For example, despite there being frequent character switches in the text, the dominant attributes at FC change points for subject 5 is motion. Further, Subjects 3, 4 and 8 react actively to emotion and verb attributes. Considering that pre-specified verbs occur infrequently in Table 6, this large number of change points (e.g., 13 out of 16 change points for subject 4) strongly implies that some subjects are very sensitive to changing verbs. We also display the locations of the detected change points with the attribute underpinning them Figure 9 (bottom panel). These plots further illustrate the complexity of textual changes. In summary, more than 90% of the change points are related to at least two types of attributes simultaneously. For these attributes, the emergence, rather than the disappearance more frequently leads to a change point, as red segments are more evident than yellow ones.

**Figure 9:**
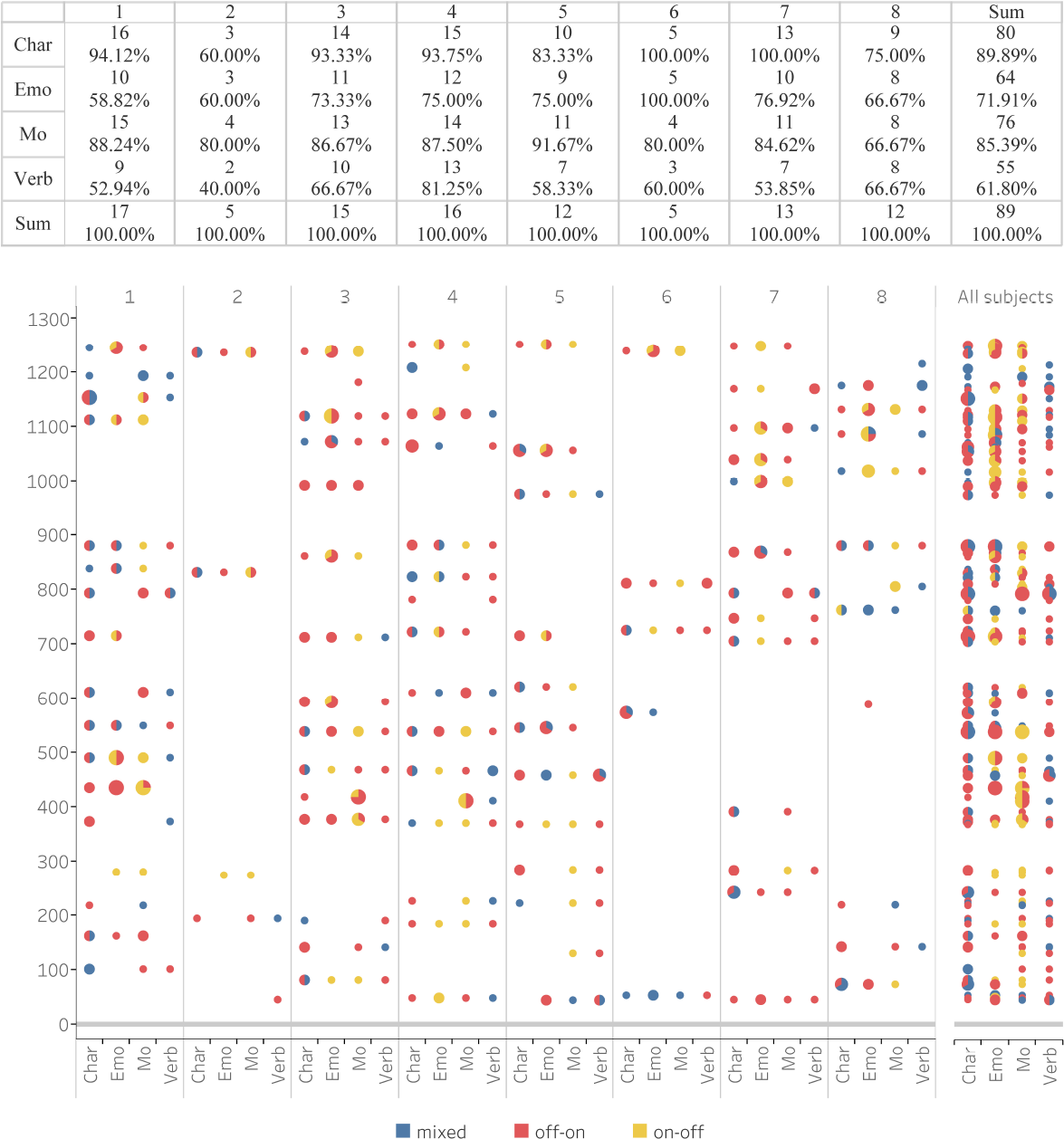
The top panel is the summation of the frequency of attributes (Char, Emo, Mo, Verb) that correspond to change points detected by NBS.V with the 5-copula family for each subject. The bottom panel displays the locations of change points corresponding to textual changes for each subject (left) and all subjects (right). Colors denote the three types of patterns.

In Figure 10, we plot the percentage of changing subdivisions (off-on) in their corresponding attributes at the change points detected by NBS.V for each subject. For each attribute, the first row (Freq) records the relative frequency of subdivisions in the original text in decreasing order, which we then compare with the frequency of change points for the same subdivision. For example, from the top left graph (Character: off-on), we can conclude that the appearance of characters such as ‘Hermione’, ‘Neville’, ‘Minerva’, and ‘Peeves’ in the text closely relate to the change points across some of the subjects (match the block size in Freq to the blocks in each of the subjects). The appearance of ‘Harry’ is not as consistent across subjects. The composition of the emotion graph is even more complex, where subjects are generally susceptible to different emotional changes of characters. Among the fifteen different emotions, ‘puzzlement’ is the most likely to be associated with changes in the brain networks across the 8 subjects. Even though positive feelings such as ‘like’, ‘relief’ and ‘pride’ seldom appear in the text, subjects 4, 5 and 7 are still sensitive to them. On the contrary, emotions such as ‘nervousness’ and ‘fear’ that cover a large portion of the text only stimulate a subset of the subjects (1, 3, 5, 7 and 8), while other subjects seem unaffected by the characters evoking such emotions. The reaction to the ‘annoyance’ of characters was the most inconsistent. Subjects 2, 3 and 6 experienced more frequent change points in terms of emotional transition to annoyance, while subjects 1, 4 and 7 share a moderate proportion of change points related to annoyance. Subjects 5 and 8, however, did not exhibit any reaction when confronting the presence of annoyance in the text. Such inconsistency could be partly due to the insufficient number of change points detected for certain subjects (e.g., subjects 2 and 6), and perhaps partly to the real heterogeneity of dynamic brain networks towards a certain emotion experienced by characters. The proportion of motion subdivisions in the text is consistent with that of subdivision change points for subjects 3, 4 and 7. The other five subjects did not react to the beginning of the characters flying nor to the colliding in the story. Further research could be carried out by dividing the ‘manipulate’ and ‘move’ motions into more detailed subdivisions. The heterogeneity across subjects becomes more evident when it comes to the variation in verbs: subjects 1 and 4 are very sensitive to the verb ‘know’; the FC of subject 8 changes intensively when the characters ‘hear’ a sound; subjects 5, 7 and 8 experience switching patterns in their brain network at the time of speaking; subject 2 fails to give any reaction to the appearance of the five types of verbs. Interestingly, brain networks across all 8 subjects remain in a relatively stable status when characters ‘tell’ something unexpected in Chapter 9. In other words, other than visual description, auditory stimulation to characters can generate more change points in the brain networks for readers.

**Figure 10:**
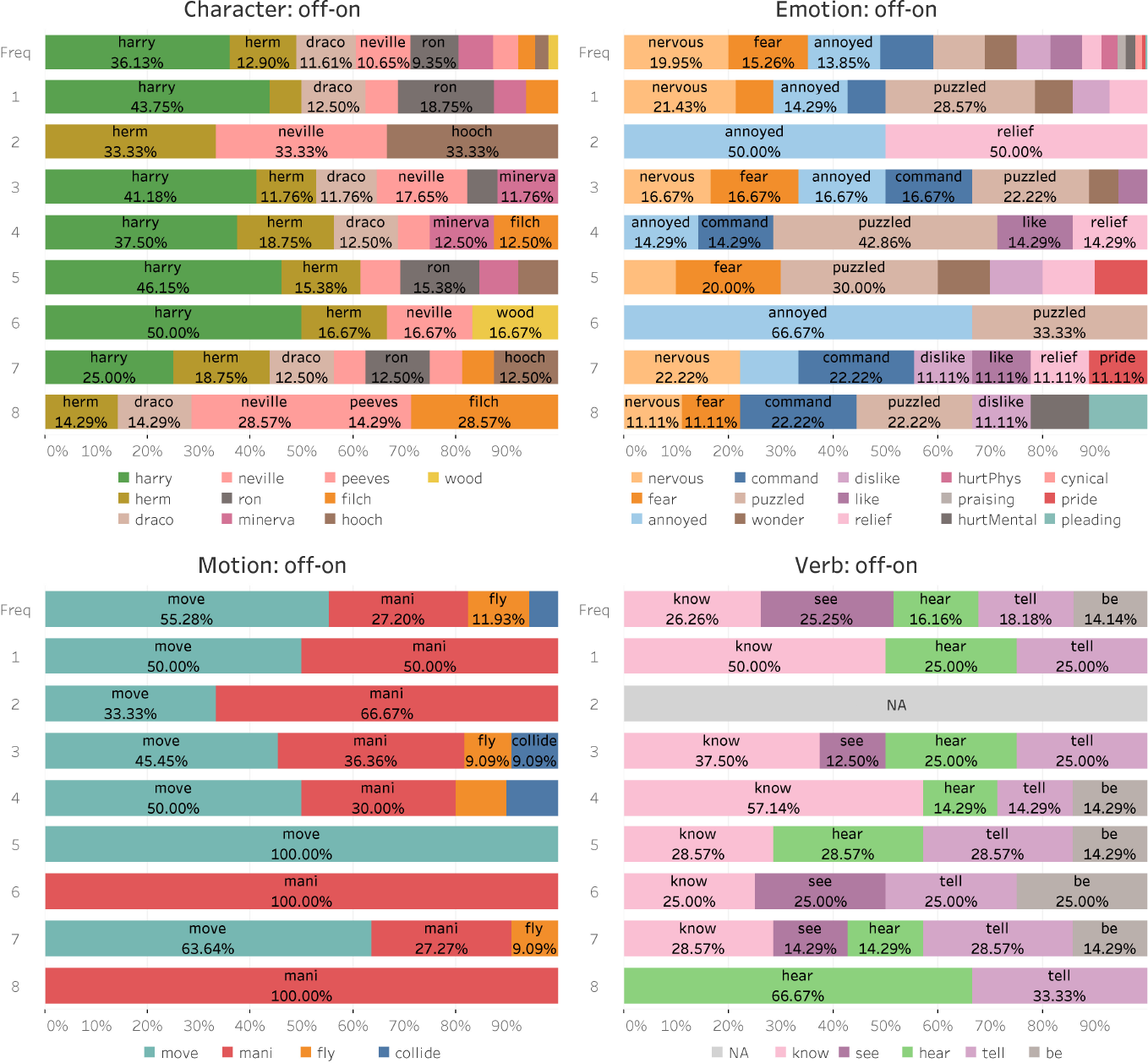
The percentage of changing subdivisions (off-on) in their corresponding attributes at the change points detected by the NBS.V combination of VCCP for each subject. For each attribute, the first row (Freq) records the relative frequency of subdivisions in the original text in order (of time).

### 4.3 SET fMRI data

The results for the SET data set can be found in the Supplementary Materials. To illustrate the importance of diverse copulae and possible tail dependence between the selected 5 ROIs, we compare the performance of the Gaussian copula against the 5-copula family in our VCCP model. We also study the effect of the choice of *δ*.

## 5 Discussion

### 5.1 Computational speed

Each table in the simulation section shows the average computation in seconds. Among the four segmentation methods, MOSUM is the most computationally efficient as it reduces the problem of a multivariate time series to a univariate time series problem. WBS is relatively slow as the computational efficiency depends heavily on the number of random draws, *M*, which should be large enough to guarantee the detection of the candidate change points. NBS and OBS have a similar moderate speed since both apply the idea of BS. With respect to inferential methods, SB depends on the number of resamples: as *N* increases, SB slows, but the result becomes more reliable. The Vuong test is computationally efficient since it has a known asymptotic distribution. For VCCP, the number of ROIs included has a large influence on its computational speed. As the number of ROIs increases, the MST methods (Cormen et al., 2009) have to consider more edges and hence more time is consumed finding the optimal vine. By specifying the vine structure (D-vine or C-vine), MST only needs to set the linking order or the central node, which can increase computational efficiency. Similarly, if we restrict the candidate family set of copulas (e.g., only the Gaussian copula) for each edge, we can decrease speed. Finally, as the experimental time course increases, the search space has a wider range, which leads to an increased computing time.

### 5.2 Extensions

The asymptotic theory for general maximum likelihood estimators is well developed (see for example, Lehmann and Casella 2006) and can be applied in the parametric vine copula case (see Chapter 7 of Czado 2019). In addition, the theoretical justification for using the Vuong test for comparing different regular vine copula models can also be found in Czado (2019). In terms of robustness, (Haff, 2013) observes that the stepwise semiparametric estimator for vine copulas (the sequential estimating algorithm in our work) is not robust to misspecification of the pairwise copulas. However, we address this issue by specifying multiple copula candidates for each edge and choose the copula type that maximizes the log-likelihood. Finally, to the best of our knowledge, there is no theoretical work on dynamic vine copula models or vine copula change point models. We leave the consistency and robustness of our model and the estimated number as well as location of change points for future work.

To deal with a large number of ROIs, it is possible to combine VCCP with dimension reduction tools such as Principal Component Analysis (PCA), Linear Discriminant Analysis (LDA) or Feature Annealed Independence Rule (Fan and Fan, 2008). In addition, the distribution of the value of BIC reduction at certain time points could also be explored under the vine copula specification. Here the classic hypothesis test could be applied instead of the SB, in order to improve computation. Furthermore, similar to MOSUM, we could consider other methods that directly explore the local maximums of the BIC reduction time series, especially if we believe there are small and subtle dynamic FC patterns in the data.

VCCP can also be applied to multi-subject data (panel data). The only distinct difference between single-subject and multi-subject data is that the sample size in each interval increases as a result of stacking different subjects’ data in the same period. Accordingly, more signal is available. For more details, see Xiong and Cribben (2021).

Applying VCCP to the Harry Potter fMRI data set, we discovered that including using more copula families in the building blocks of VCCP results in more change points detected. To further explore the effect of copula families on the performance of VCCP, we extend our choice to other copulas including Joe, Clayton-Gumbel (BB1), Joe-Gumbel (BB6), Joe-Clayton (BB7) and the Joe-Frank (BB8) copula, and re applied the change point analysis with *δ* = 40 on the Harry Potter fMRI data (Figure 11). In comparison to the single Gaussian copula (see Figure 6), copulas with heavy tail dependence allow VCCP to possibly detect more change points. This was also evident when we add t, Clayton, Gumbel, Frank (the previous 4 copulas) to the family or add Joe, BB1, BB6, BB7 and BB8 (the new 5 copulas). When we combined all the 9 non-gaussian copulas with the Gaussian copula (last panel), the number of detected change points exceeds the other cases, although not as evident in moving from Gaussian to 5 family copulae. That is, the number of change points detected by VCCP is not strictly linearly correlated with the number of copula families.

**Figure 11:**
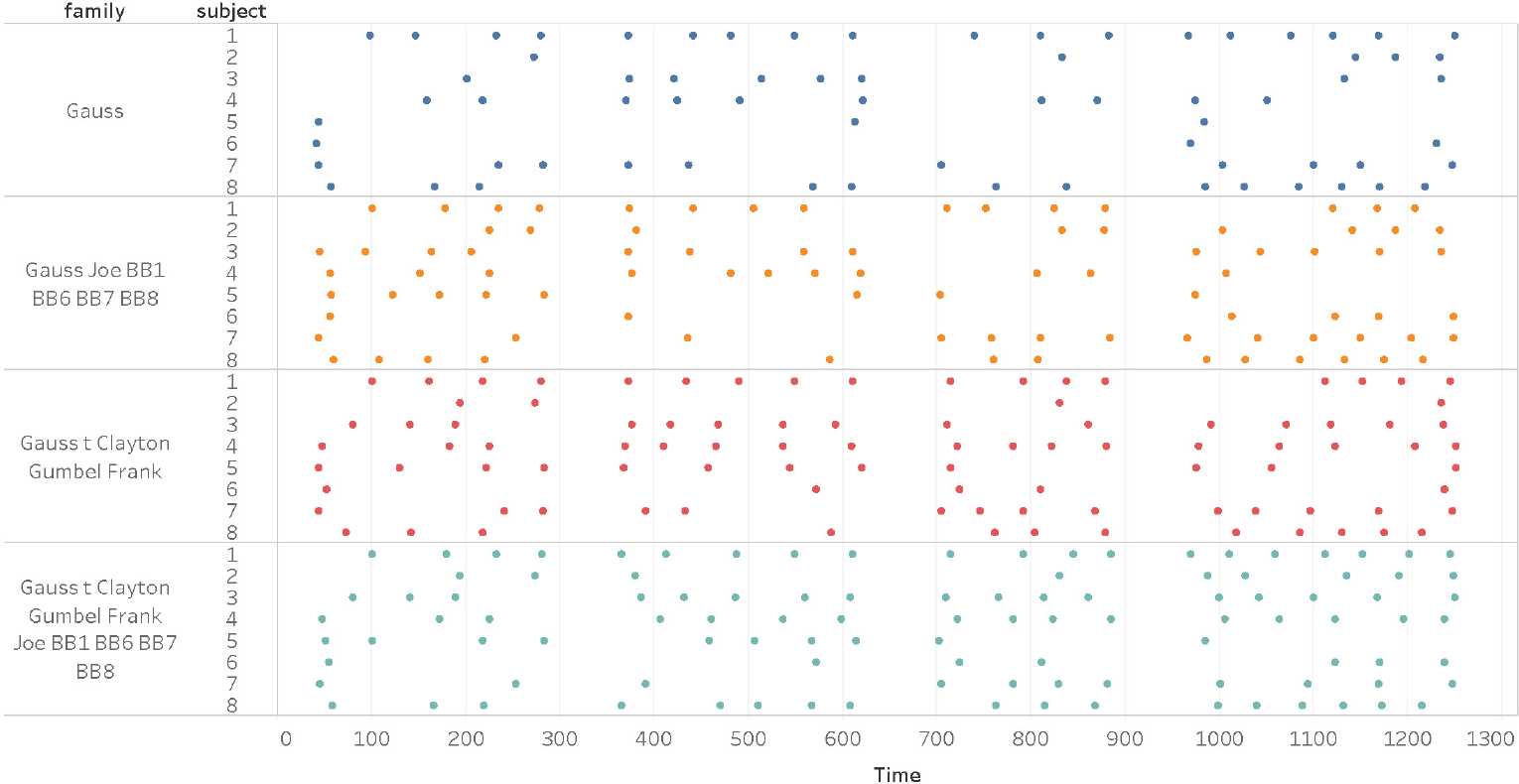
The four panels represent the change points from NBS.V using different copula family combinations with *δ* = 40, from the Harry Potter task-based fMRI study.

### 5.3 Limitations

While different combinations of segmentation and inference methods in VCCP can be used in different settings, some combinations have shortcomings under certain circumstances. For example, both OBS and NBS segmentation methods have at their core BS, which is limited in that it can only find one change point at a time and then recursively splits the data. They are also restricted by *δ*, the minimum between change points and hence the total number of possible change points. It is important that VCCP remains robust to alternative choices to this parameter and the practitioner has the option to obtain more change points by decreasing this value. We studied the robustness of VCCP to changing *δ* for the SET fMRI set in the Supplementary Materials. Overall, we find that the method finds consistency in the location of the change points but obviously the number of change points alter for different *δ*. For a theoretical perspective on the minimum between change points, *δ*, in binary segmentation, see Fryzlewicz (2014).

The MOSUM and WBS methods allow us to bypass specifying this minimum distance between change points. However, MOSUM is excessively conservative when changes in the FC are subtle (change in edge strength, see Simulation 6). This may due to MOSUM’s indirect detection in this setting, which reduces the problem to a univariate time series (in our case the BIC reduction series) and then chooses the local maximums as candidate change points. For resting-state fMRI data sets, where subjects smoothly transit between states, MOSUM might confuse the change points with random variations or might amalgamate several moderate peaks into one change point, which ultimately leads to fewer change points. Finally, for the task based fMRI data sets, we obtained conservative results for WBS. This is understandable given WBS stops if the point detected is not a change point or no random intervals remains. If the number of random intervals is not sufficient and the first candidate change point is close to the middle of the time course, the second condition is more likely to occur. In future work, we intend to explore other change point segmentation methods such as isolate detect (Anastasiou et al., 2022), that allow for dense change points and change points within short intervals.

## 6 Conclusion

In this paper, we introduce a new methodology, called Vine Copula Change Point (VCCP), that estimates change points in a vine copula structure. It is motivated by the fact that no other change point method, to the best of our knowledge, considers possible non-linear and non-Gaussian dynamic functional connectivity. Furthermore, the layers of a vine can be cut, which leads to a simplified and sparse network, and requires less computation time when the dimension of the data set expands.

We also compared different combinations of segmentation methods (NBS, OBS, MOSUM and WBS) and inference methods (Vuong test and the stationary bootstrap) in VCCP. Overall, different combinations serve different needs. NBS with the Vuong test provides the most robust change points for the task-based fMRI data sets, requires fewer inputs, and is more computationally efficient. Although not studied here, perhaps WBS is more suitable for resting-state fMRI data sets as it is sensitive to dense but weak change points in the network. Overall, we found that OBS (used by most previous change point methods in the neuroscience literature) failed to perform well. SB and the Vuong test are both suitable for inference, but Vuong is computationally faster.

We showed using an extensive simulation study that VCCP performs well on non-Gaussian data as well as multivariate normal data and on Vector autoregression (VAR) data, even in challenging scenarios where the subject alternates between task and rest. The MVN data can be considered whitened data, while the VAR data can be considered un-whitened data. However, the simulation results show the deterioration in performance is not drastic. In addition, the task-based fMRI experimental results clearly indicate the presence of FC change points that are due to non-linear dependence, which has not been explored in a dynamic fashion before in the analysis of neuroimaging data.

While VCCP was applied to task based fMRI data, it could seamlessly be applied to resting-state fMRI data, Electroencephalography (EEG) or Magnetoencephalography (MEG), and electrocorticography (ECoG) data or other time series applications where the network structure is changing. The outputs of VCCP could also be used as an input into a classification model for predicting brain disorders. Finally, the R package **vccp** (Xiong and Cribben, 2021) implementing the methodology from the paper is available from CRAN.

## Acknowledgments

X. Xiong was supported by a China Council Scholarship and the work of I. Cribben was supported by the Natural Sciences and Engineering Research Council (Canada) grant RGPIN-2018-06638 and the Xerox Faculty Fellowship, Alberta School of Business.

## Supplemental Materials description

### R Code and Data

The supplemental files for this article include files containing R code and data for reproducing the simulation study and some of the fMRI results in the paper. Please see the README.txt file for specific details on how to run the code.

### Supplementary materials

The supplemental files for this article also include the following: (i) definitions of Kendall’s *τ* and copula formulae, (ii) additional information on the Harry Potter fMRI data set, (iii) additional information on the anxiety fMRI data set, (iv) figures explaining the setup of the non-gaussian simulations in the main paper (v) descriptions of the setup of the multivariate normal (MVN) and the vector autoregression (VAR) simulations, (vi) additional results from the Harry Potter and the anxiety fMRI data sets, and (vii) simulation results from the the MVN and the VAR simulations

## Supplementary Materials

### 1 Kendall’s *τ* and copula formulae

**Definition 1** (**Kendall’s** *τ*) *Let (X*_1_, *Y*_1_*) and (X*_2_, *Y*_2_) *be two independent pairs of random variables with joint cdf F and marginal distributions F_X_ and F_Y_. Kendall’s τ is given by*

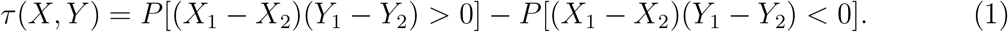

*The estimate of Kendall’s τ is given by :*

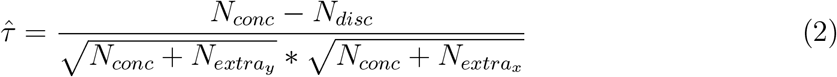

*where N_conc_ is the number of concordant pairs (the orderings of the two Xs is the same as the ordering of the two Ys), N_disc_ is the number of discordant pairs (the ordering of the two Xs is opposite from the ordering of the two Ys) and* 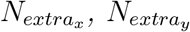 *are the numbers of extra-X pairs* (*i, j*) *such that X_i_* = *X_j_, i* ≠ *j and extra-Y pairs* (*i, j*) *such that Y_i_* = *Y_j_, i* ≠ *j, respectively*.

The formulas for the copulae we consider in this work are given by:

#### 1. Gaussian Copula

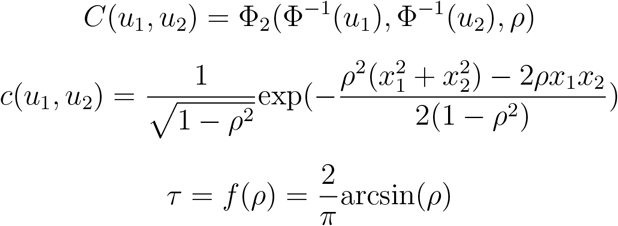

where *x*_1_ = Φ^*−*1^(*u*_1_), *x*_2_ = Φ^*−*1^(*u*_2_), and Φ_2_(*., ., ρ*) is the bivariate normal cdf with two standard univariate marginal cdfs and a Pearson correlation coefficient, *ρ*.

#### 2. *t* Copula

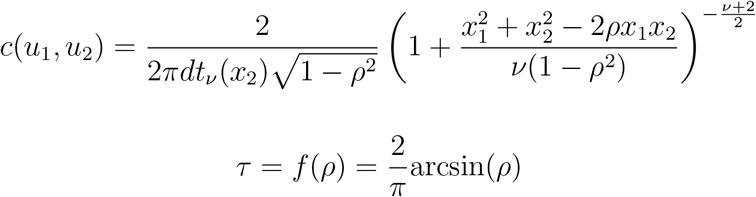

where 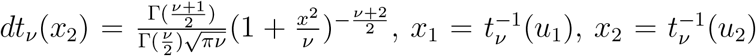 and 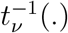 is the inverse cdf of univariate *t* distribution with *ν* degrees of freedom.

#### 3. Frank Copula

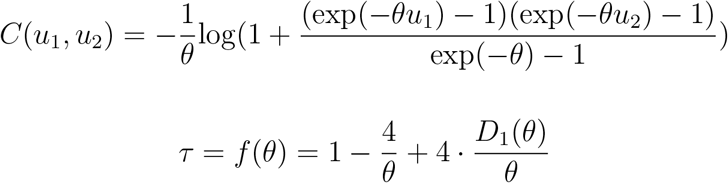

where 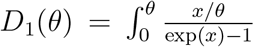 and *θ* ∈ (−∞, +∞). As *θ* → −∞, the two variables linked by the Frank copula become more negatively correlated. As *θ* → +∞, the two variables become more positively correlated.

#### 4. Clayton Copula

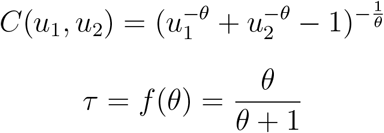

where *θ* ∈ (−1, ∞)\0. Clayton copulas are only used to describe positive dependence. As *θ* → 0, the two variables become more independent, and as *θ* → ∞, the two variables become more positively correlated.

#### 5. Gumbel Copula

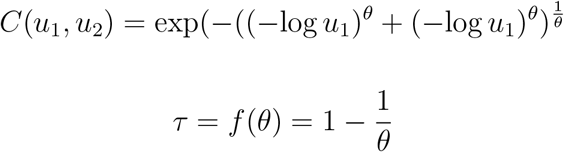

where *θ* ∈ (1, ∞). Gumbel copulas are only used to describe positive dependence. When *θ* = 1, the two variables are independent, and as *θ* → ∞, the two variables become more positively correlated.

### 2 Harry Potter fMRI data set

Table 1 provides information on the 14 ROIs from the AAL atlas extrfrom subjects in the Harry Potter task-based fMRI data set.

**Table 1:** Information on the 14 ROIs from the AAL atlas extracted from subjects in the Harry Potter task-based fMRI data set.

| ROI id | Regions | number of voxels | Label |
| --- | --- | --- | --- |
| 1 | Angular gyrus | 838 | AG |
| 2 | Fusiform gyrus | 1447 | F |
| 3 | Inferior temporal gyrus | 2013 | IT |
| 4 | Inferior frontal gyrus, opercular part | 747 | IFG 1 |
| 5 | Inferior frontal gyrus, orbital part | 1008 | IFG 2 |
| 6 | Inferior frontal gyrus, triangular part | 1355 | IFG 3 |
| 7 | Middle temporal gyrus | 2795 | MT |
| 8 | Occipital lobe | 6371 | O |
| 9 | Precentral gyrus | 2030 | PCG |
| 10 | Precuneus | 2014 | PC |
| 11 | Supplementary motor area | 1322 | SM |
| 12 | Superior temporal gyrus | 1635 | ST |
| 13 | Temporal pole | 1353 | TP |
| 14 | Supramarginal gyrus | 550 | SG.R |

### 3 Anxiety fMRI data set

This fMRI data set (SET data) was taken from an anxiety-inducing experiment. Before the start of the experiment, participants were told that they would be given 2 minutes in the scanner to present a 7-minute speech, though there was a small chance that they might not be randomly selected to give the speech. Once the fMRI acquisition began, subjects rested for 2 min and then an instruction slide showing the topic – ‘Why you are a good friend’ could be seen for 15 s. When the slide disappeared, subjects went through a 2min silent brainstorming which was interrupted by another instruction slide that informed participants they would not have to give the speech. Then they rested for 2min which completed the functional run. Data was acquired and preprocessed the same way as in the previous work (Wager et al., 2009; Cribben et al., 2012, 2013). The experimental data set consists of 5 ROIs: (1) visual cortex, (2) left superior temporal sulci, (3) ventral striatum, (4) right superior sulci and (5) ventromedial PFC that were created by averaging the voxel time series across the entire region. *N* = 26 subjects were involved in the experiment, which lasted for *T* = 215, with TR = 2 s. Our VCCP methodology weas applied to individual subjects to learn if the stressor onset triggered a change point in the FC network between the 5 ROIs.

### 4 Simulation setup

#### 4.1 Non-Gaussian simulations

Figures 1, 2, and 3 are visual displays of the non-Gaussian simulations in the main paper.

**Figure 1:**
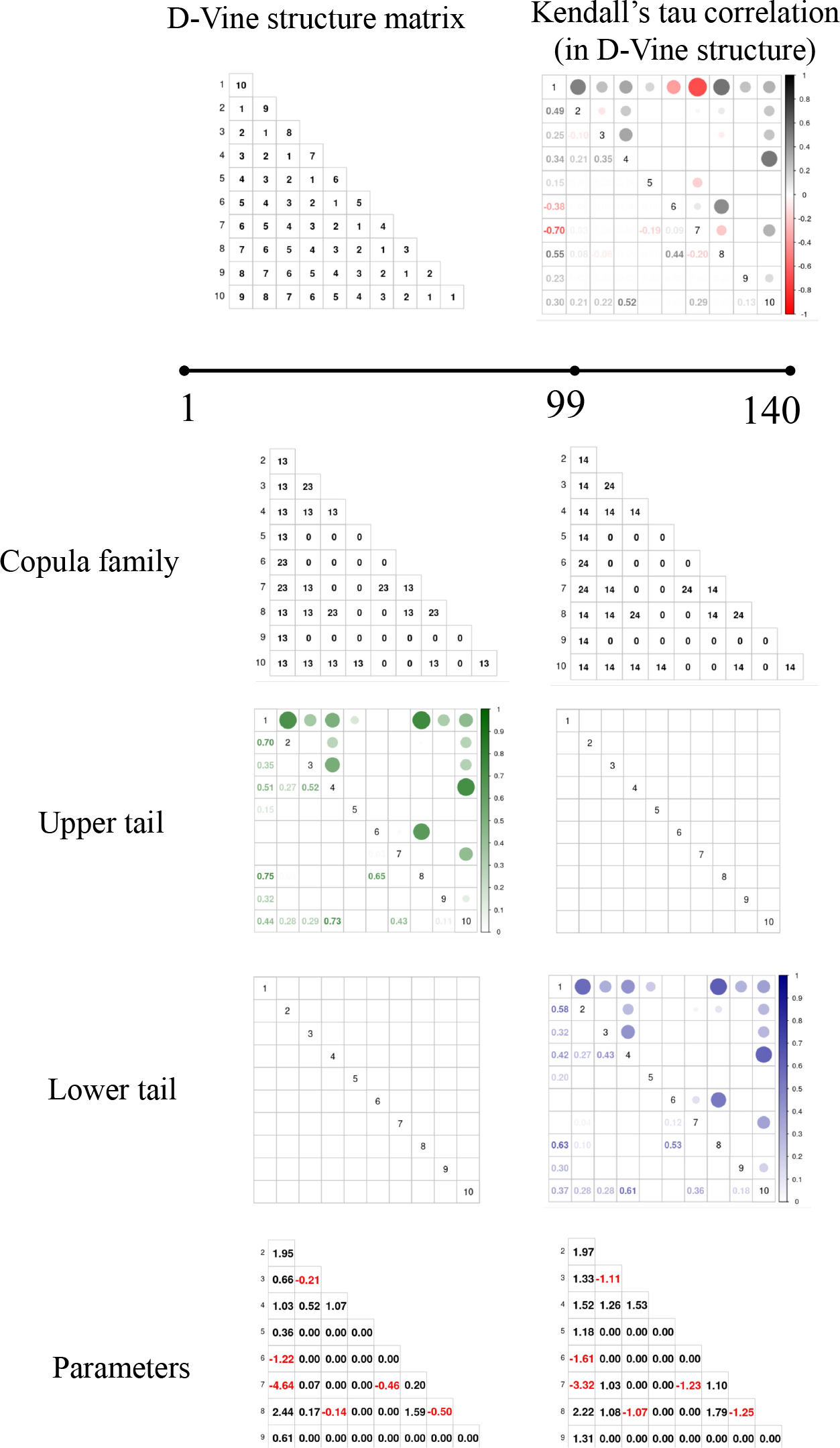
A visual display of Simulation 2 with one change point in the vine copula structure at time point *t* = 99. The left column represents the structure for time points (1:99), while the right column represents the structure for time points (100:140). The first row depicts the D-vine structure and corresponding Kendall’s *τ* which are constant across the time series. The second row depicts the copula type. Here 0 represents independence, 13 represents the clayton copula rotated 180 degrees, 23 represents the clayton copula rotated 90 degrees, 14 represents the gumbel copula rotated 180 degrees, and 24 represents the gumbel copula rotated 90 degrees. The third and fourth rows depict the upper and lower tail dependence structures, respectively, for time points (1:99) and (100:140). The fifth row depicts the changes in the parameters values.

**Figure 2:**
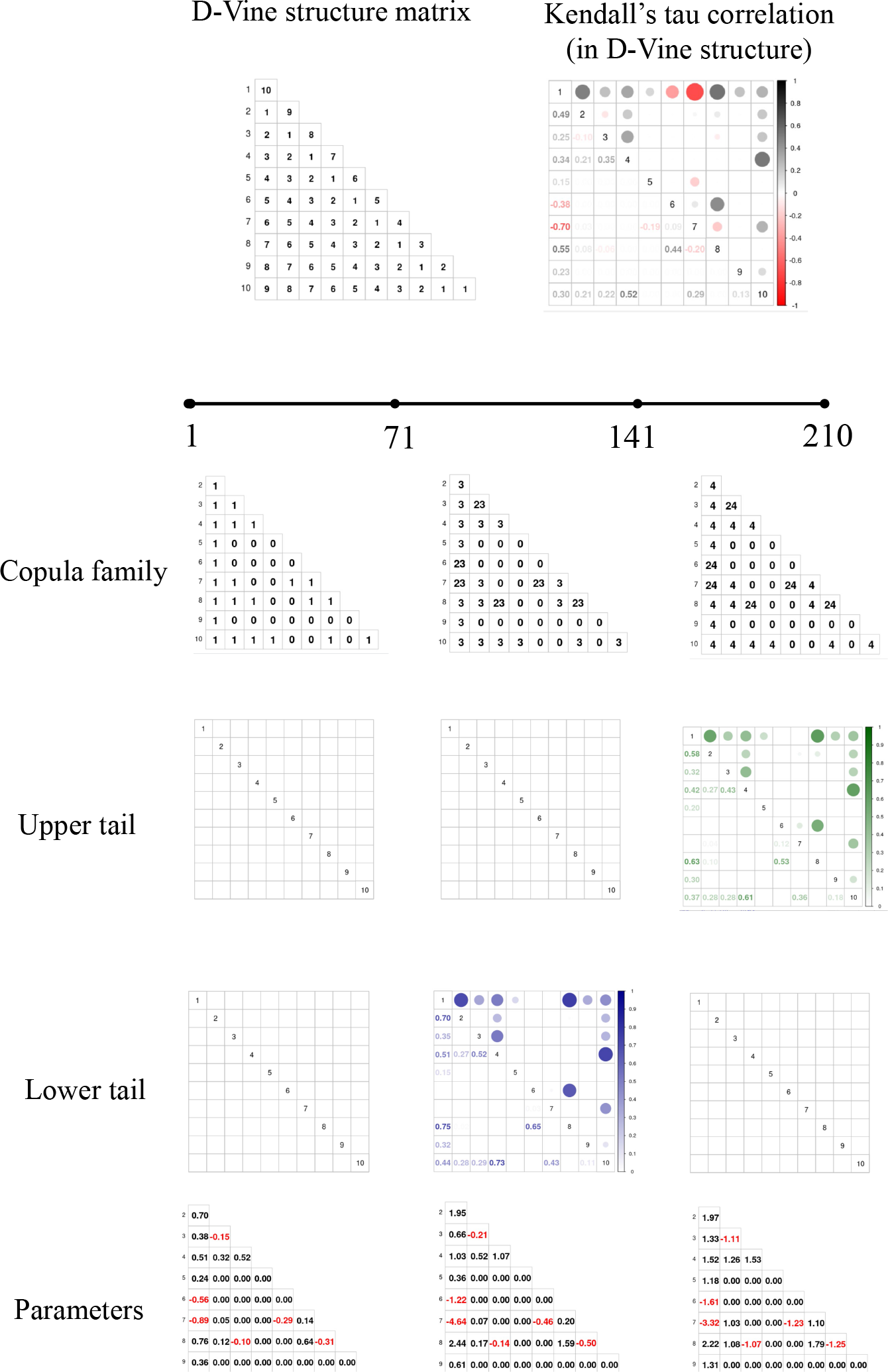
A visual display of Simulation 3 with two change points in the vine copula structure at time points *t* = 71, 141. The first, second and third columns represent the structure for time points (1:71), (72:141), and (142:210), respectively. The first row depicts the D-vine structure and corresponding Kendall’s *τ* which are constant across the time series. The second row depicts the copula type. Here 0 represents independence, 13 represents the clayton copula rotated 180 degrees, 23 represents the clayton copula rotated 90 degrees, 14 represents the gumbel copula rotated 180 degrees, and 24 represents the gumbel copula rotated 90 degrees. The third and fourth rows depict the upper and lower tail dependence structures, respectively, for time points (1:71), (72:141), and (142:210). The fifth row depicts the changes in the parameters values.

**Figure 3:**
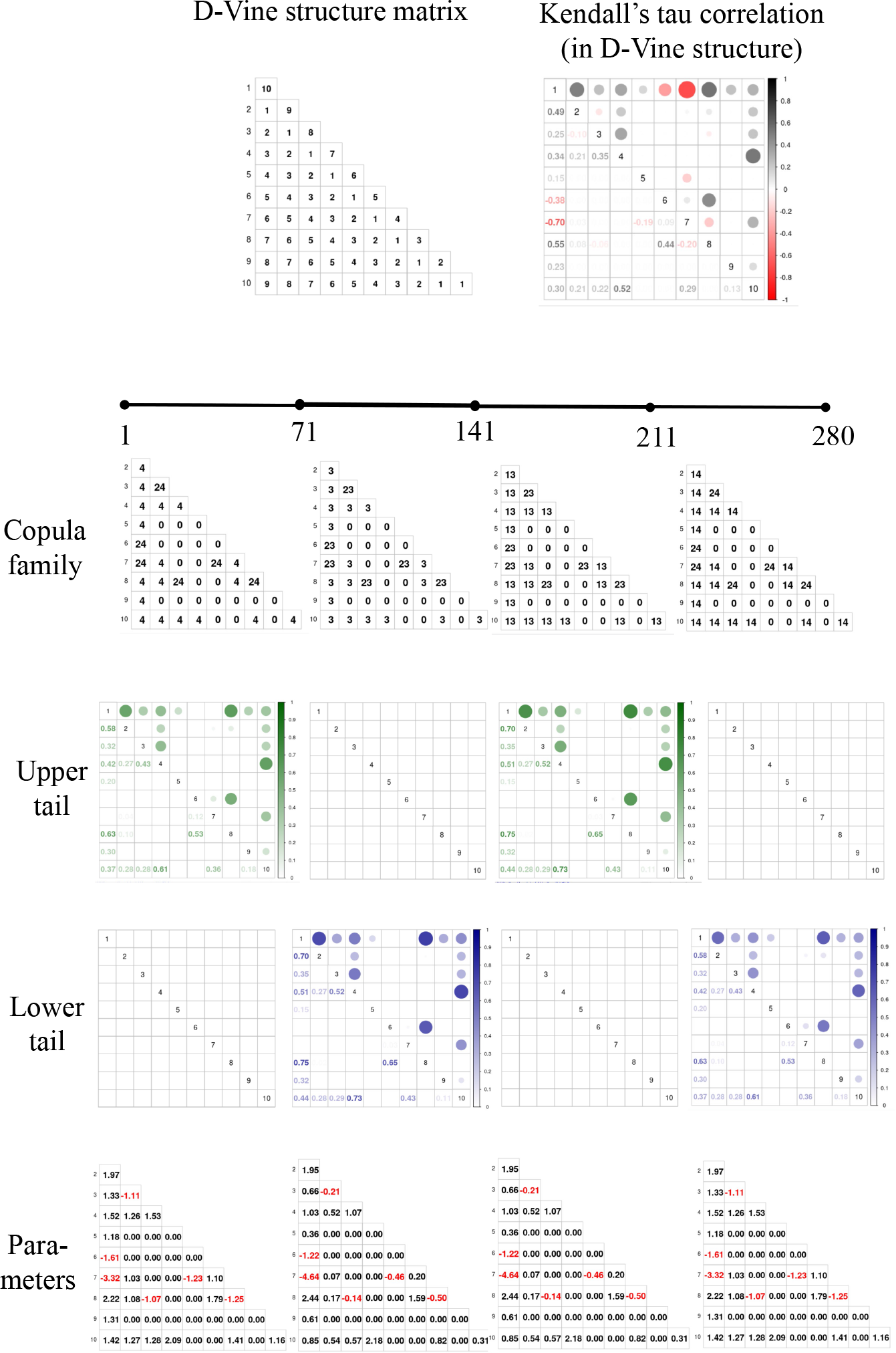
A visual display of Simulation 4 with three change points in the vine copula structure at time points *t* = 71, 141, 211. The first, second, third and fourth columns represent the structure for time points (1:71), (72:141), (142:210), and (211:280) respectively. The first row depicts the D-vine structure and corresponding Kendall’s *τ* which are constant across the time series. The second row depicts the copula type. Here 0 represents independence, 13 represents the clayton copula rotated 180 degrees, 23 represents the clayton copula rotated 90 degrees, 14 represents the gumbel copula rotated 180 degrees, and 24 represents the gumbel copula rotated 90 degrees. The third and fourth rows depict the upper and lower tail dependence structures, respectively, for time points (1:71), (72:141), (142:210), and (211:280). The fifth row depicts the changes in the parameters values.

#### 4.2 MVN and VAR simulations

Table 2 provides a summary of the multivariate normal (MVN) and the Vector Autoregression (VAR) simulations, while Figure 4 provides a visual display of the MVN simulations only (Simulations 5-8). For more details on the strength of the correlation, see the Appendix in Cribben (2019). A VAR model is a generalization of a univariate AR process. Given a *p*-dimensional multivariate time series **X**, the lag-1 vector autoregression model, VAR(1), is defined as **X**_*t*_ = Π_1_**X**_*t−*1_ + ϵ_*t*_, *t* = 2, …, *T*, where Π_1_ is an (*p* × *p*) coefficient matrix and _*t*_ is an (*p* × 1) unobservable mean white noise vector process with time invariant covariance matrix. The VAR model is used to reconstruct the linear inter-dependency element prevalent among multivariate time series applications such as fMRI data. For both data types, the mean vector is always zero since we only focus on change points in connectivity. In the simulations overall, we attempt to emulate the properties in fMRI data. In particular, we consider no change points (null data), multiple change points, general changes in the correlation/network structure, changes in the strength of the connectivity, pre-whitened (MVN) and un-whitened (VAR) data, and inspired by the experimental fMRI data, an off-on-off pattern. We run each simulation using 100 iterations.

**Table 2:** A summary of the simulated data sets. MVN and VAR denote multivariate normal and vector autoregression data, respectively. Weak means that only the edge weights are adjusted between change points with the non-zero edges themselves remaining constant over the entire time course, while small means that only small changes in the networks are present between change points. The ROI connectivity is defined by cliques such as 2-6-9.

| Sim | Data type | Change type | $p$ | Locations of change points | ROI connectivity |
| --- | --- | --- | --- | --- | --- |
| 5 | MVN | Normal | 15 | 61;121;181;241 | 1-60s: 1-3-5-8-10 & 2-6-9;<br>61-120s: 1-4-5-7-9;<br>121-180s: 2-3-4-6-8-10 & 1-7;<br>181-240s: 2-7-9;<br>241-300s: 1-5-6 & 7-8-10. |
| 6 | MVN | Weak | 15 | 61;121;181;241 | 1-300s: 1-4-8-14 & 3-6-9;<br>(Changes in connecting strength) |
| 7 | MVN | Small | 15 | 76;151;226;301;376;451;526 | 1-75s & 301-375s & 526-600s:<br>1-5-8-10-15 & 3-6-13;<br>76-150s: 3-6-13;<br>151-225s: 1-5-8-10 & 3-6-13;<br>226-300s: 1-5-8-10-15;<br>376-450s: 1-5-8-10-15 & 3-6-13;<br>451-525s: 1-5 & 3-6-13. |
| 8 | VAR | Normal | 15 | 61;121;181;241 | 1-60s: 1-3-5-8-14 & 4-15;<br>61-120s: 2-6-10 & 3-9-13;<br>121-180s: 4-6-11-14 & 5-12;<br>181-240s: 1-4-7;<br>241-300s: 8-10-13-15. |
| 9 | VAR | Weak | 15 | 61;121;181;241 | 1-300s: 1-5-10-15 & 2-9-14;<br>(Changes in connecting strength) |
| 10 | VAR | Small | 10 | 76;151;226;301;376;451;526 | 1-75s & 301-375s & 526-600s:<br>1-5-8-10 & 2-7-9;<br>76-150s: 2-7-9;<br>151-225s: 1-5-8 & 2-7-9;<br>226-300s: 2-7-9;<br>376-450s: 1-5-8-10 & 2-7-9;<br>451-525s: 5-8 & 2-7-9. |

**Figure 4:**
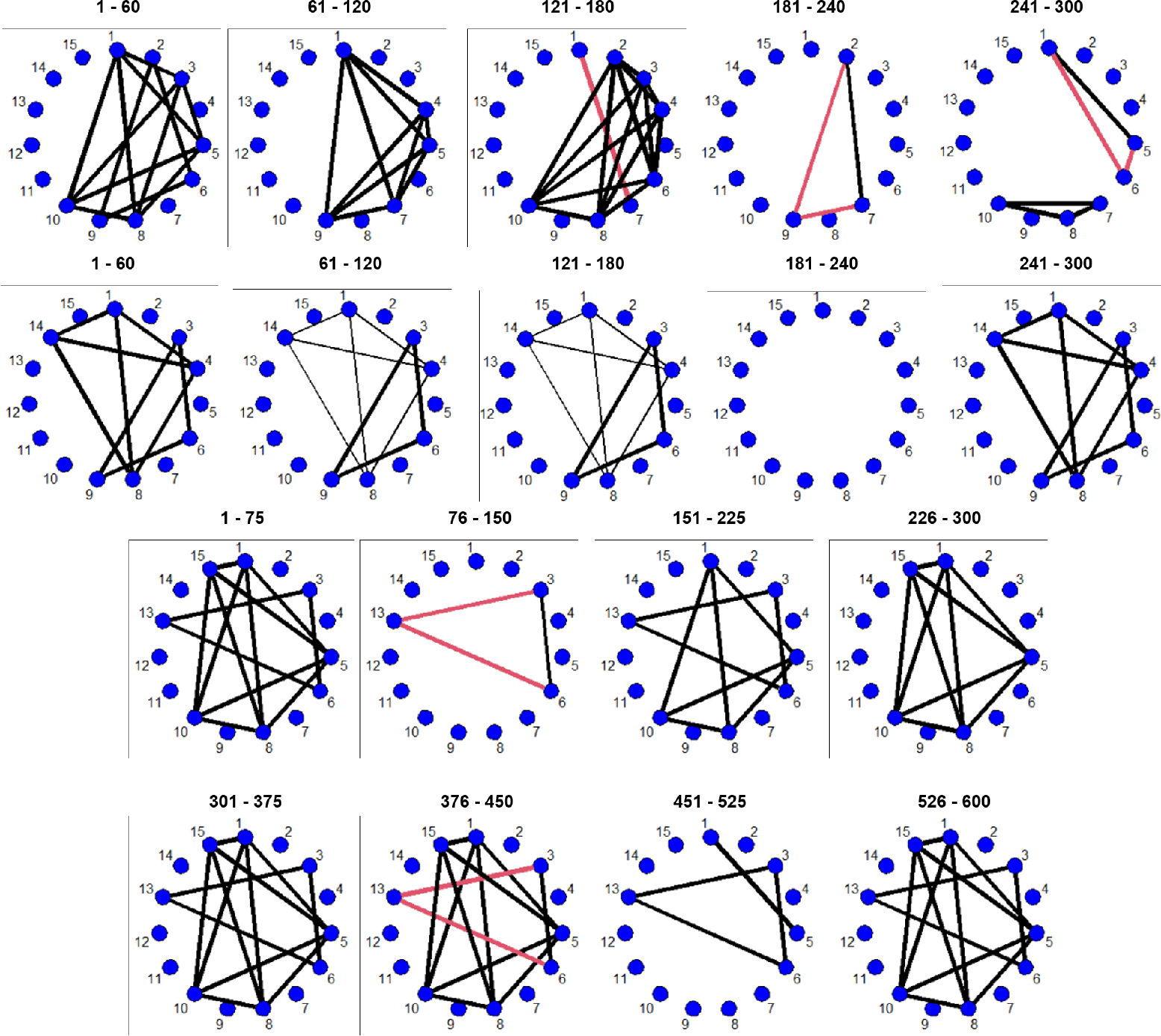
The network structures for the simulations: (first row) MVN data set with four change points (Simulation 5), (second row) MVN data set with four change points of small magnitude (Simulation 6), and (third and fourth rows) MVN data set with seven change points (Simulation 7). The time series are represented by nodes, black (red) edges infer positive (negative) connectivity, and the strength of connection between the regions is directly related to the thickness of the edges, that is, the thicker the edge the stronger the connection.

### 5 Results

### 5.1 Harry Potter fMRI data

If we turn to the ‘on-off’ subdivision pattern in Figure 5, all 8 subjects remain in a brain network longer than the subdivision when confronting the disappearance of a certain verb or the transfer of focus from a certain character. For all attributes, char, Verb, Emo, the ‘on-off’ subdivision pattern are less likely to coincide with change points, however, for attribute Mo, there is correspondence between the subdivision pattern and the detected change points. Yet, for the attribute Emo, the abnormally higher portion of the ‘hurtPhys’ element in subjects 1 and 8, the ‘like’ element in subjects 4 and 5, the ‘cynical’ element in subject 7 still deserves exploration. For all textual features, the functional network of the subjects’ brain varies in accordance with the integrated variation of the text, instead of a single, specific attribute. Considering the complex component of the reading material and the collaborative work of our brain, it is hard to directly relate a connection variation between two nodes to one special type of textual change.

**Figure 5:**
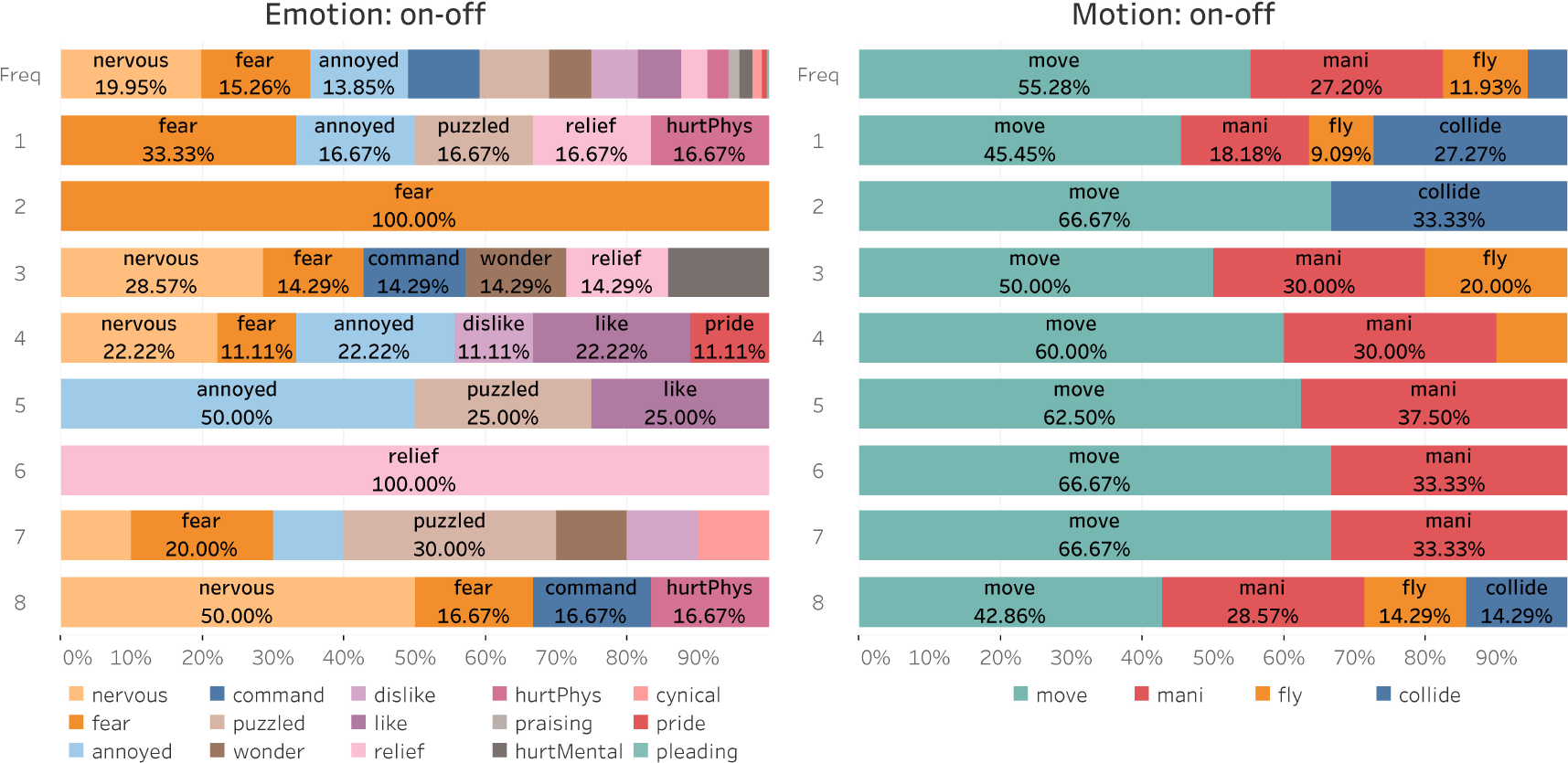
The percentage of changing subdivisions (on-off) in their corresponding textual types at change points for each subject. For each attribute, the first row (Freq) records the relative frequency of subdivisions in the original text in order (of time).

Next, we focus on general change points as a response to plot changes. By selecting a representative set of change points shared by multiple subjects and the contexts read before and after the change, we display the heterogeneity and homogeneity among subjects when confronting the same combined textual variation.

#### The first flying experience

> Blood was pounding in his ears. He mounted the broom and kicked hard against the ground and up, up he soared; air rushed through his hair, and his robes whipped out behind him – and in a rush of fierce joy he realized he’d found something he could do without being taught – this was easy, this was wonderful. |^*s*5^ He pulled his broomstick up a little to make it even higher |^*s*4^ and heard screams and gasps of girls back on the ground |^*s*1^ and an admiring whoop from Ron. |^*s*3^ He turned his broomstick sharply to face Malfoy in midair …

We detect change points in subjects 1, 3, 4 and 5 that coincide with Harry’s first flying experience. The exact locations of the change points are marked as vertical lines in the quotation above, and the corresponding FC networks before and after the change points are provided in Figure 6 (we only plot the positive correlation between nodes). Several positive emotions such as ‘joy’ and ‘admiring’ emerged during this extract, which are rare emotions in Chapter 9. As a reaction to such a ‘positive’ textual change, the Kendall’s *τ* network of all four subjects become more sparse (Figure 6, top left). For subject 1, the close relationship between ROIs IFG (IFG1, IFG2, IFG3), F and RSG disappears; for subject 3, most of the edges disappear while the connections within the sub-network (ST-AG-MT) become stronger. In subject 4’s network, there is disconnection between ST, TP, MT and IFG, with only seven edges remaining. The middle temporal gyrus is sensitive to visual motion (flying), and while traditional language processing areas include the inferior frontal gyrus (Broca’s area), superior temporal and middle temporal gyri, supramarginal gyrus and angular gyrus (Wernicke’s area), there is evidence that structures in the medial temporal lobe have a role in language processing (Tracy and Boswell, 2008). The FC networks of subject 5 before and after the change point are more closely related, compared to the other three subjects, with decreasing Kendall’s *τ* before and after the change point.

Contrary to the decreasing Kendall’s *τ*, the lower tail networks (Figure 6, bottom left) increase in degree edges while the upper tail dependence networks (bottom right) vary across subjects. Based on the increasing number of grey dashed lines in the networks after the change point, we can ascertain that although some edges disappear, some of the tail dependence edges remain or appear. For example, for subjects 1 and 3, Kendall’s *τ* between ROIs TP and IFG3 disappears but the corresponding lower tail dependence appears or stabilizes, respectively. The temporal pole has been associated with several high-level cognitive processes: visual processing for complex objects and face recognition, naming and word-object labelling, semantic processing in all modalities, and socio-emotional processing (Herlin et al., 2021). Accordingly, changes in the connection (strength) of the tail dependence has less to do with the variation in Kendall’s *τ*. Rich colors in the top right sub-graphs for each subject represent the diversity in the best fitted copula types for edges between each pair of nodes. As Harry is experiencing the joy of his first flying experience, the dominant copulas shift to the Clayton and Gumbel copula after the change point, which is due to the fact that Clayton and Gumbel copulas provide a better fit to the lower and upper tail dependence than the other three available copulas. This is best illustrated by subject 5, where both types of tail dependence networks become denser and the preponderant F copula loses its dominance after time point *t* = 367.

#### Encountering with the three-headed dog

> “What?” Harry turned around – and saw, quite clearly, what. For a moment, he was sure he’d walked into a nightmare – this was too much, on top of everything that had happened so far. |^*s*7^ They weren’t in a room, as he had supposed. They were in a corridor. The forbidden corridor on the third floor. |^*s*8^ And now they knew why it was forbidden. They were looking straight into the eyes of a monstrous dog, |^*s*3^ a dog that filled the whole space between ceiling and floor. It had three heads. Three pairs of rolling, mad eyes; three noses, twitching and quivering in their direction; three drooling mouths, saliva hanging in slippery ropes from yellowish fangs.

This event is the most thrilling in Chapter 9 of *Harry Potter and the Sorcerer’s Stone*. The exact locations of the change points for subjects 3, 7 and 8 are marked as vertical lines in the quotation above, and the corresponding FC networks before and after the change points are provided in Figure 7. Both consistency and heterogeneity exist in their brain networks. For subject 3, the most obvious change after the change point at time *t* = 1179 is the disappearance of the upper tail network between ROIs RSG, IT, IFG1, IFG2 and IFG3. In addition, strong connections between ROIs TP, AG, ST and F show up in Kendall’s *τ* and the upper tail dependence network after the change point. For subject 7, the Kendall’s *τ* network after the change point resembles that of subject 3, especially for sub-networks MT-ST-AG-F-IT and IFG1-SM-RSG-PCG. The first sub-network hub at MT is also present in the networks of subjects 3 and 7 before their change points, though subject 7’s former network had two hub ROIs (MT and ST) and involved more related ROIs (e.g., AG, IFG3). The supramarginal gyrus is essential for visuospatial awareness and it may generate the fictive dream space necessary for the organized hallucinatory experience of dreaming (Pace-Schott and Picchioni, 2017). The sequence of events occurring in the book at this time are dreamlike with the description including the word “nightmare” and the characters moving from a room to a corridor without their own movement. The edge between ROIs SM and IFG1 also shares a reappearing pattern in subjects 7 and 8. Interestingly, subject 8 has a lower tail dependence network hub at ROI ST similar to the Kendall’s *τ* network of subject 7 before the change point. However, the hub node pattern around ROI ST fades and shifts to a stronger Kendall’s *τ* network between SM, RSG, PCG, IFG1 and IFG3 after the change point, which is unique during the post-change period among the three subjects. There are also between-edge, between-subject and between-period variations in the best fitted copula functions, which indicate the heterogeneity in the dependence type.

**Figure 6:**
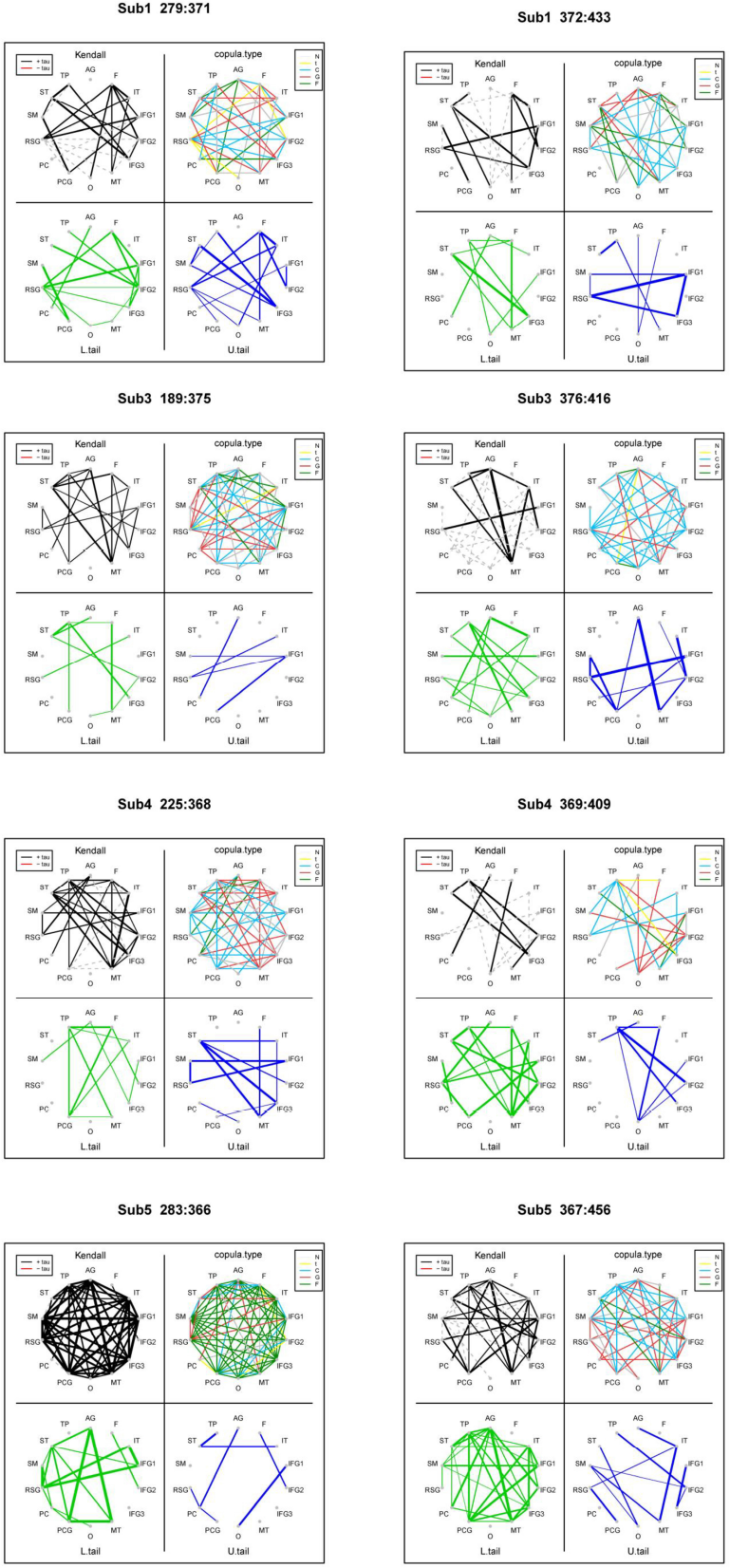
The FC networks estimated using the NBS.V combination of VCCP for subjects 1, 3, 4 and 5 at the change point that coincides with Harry’s first flying experience. The left plots represent the networks before the change point while the right plots represent the networks after the change point. The top left, top right, bottom left and bottom right networks represent the estimated Kendall’s *τ*, the lower tail dependence, the upper tail dependence, and the optimal copula function, for each edge between pairs of nodes, respectively. Black (red) lines in the top left graphs represent positive (negative) Kendall’s *τ* correlation coefficients. Dashed lines indicate edges between nodes with tail dependence but statistically insignificant Kendall’s *τ*. Green and blue edges in the bottom left and right graphs represent the lower and upper tail dependence, respectively. The various colored lines on the top right graphs row indicate the optimal copula family for edges, with grey, yellow, blue, red and green indicating the Gaussian, *t*, Clayton, Gumbel and Frank copula, respectively.

In general, when subjects share common change points, their dynamic reaction reflected in the FC networks embodies not only uniformity towards a certain combination of textual changes, but also individual heterogeneity. However, to further confirm the specific textual feature leading to the change point, more refined experiments controlling other irrelevant variables are required. Otherwise, we can only attribute the change in brain networks to combined textual factors.

### 5.2 Anxiety fMRI data

In Figure 8, we display the results from applying all segmentation methods of VCCP (with the Vuong test) to the SET fMRI data set. The solid vertical lines indicate the times of the showing and of the removal of the visual cues. We compare the change points detected across the segmentation methods using both the Gauss and the 5-copula family. Similar to the simulation results, the best method segmentation method is NBS, finding the most change points that are close to the times of the showing and of the removal of the visual cues (change points are more tightly clustered around the four vertical lines). OBS and MOSUM have a similar performance to NBS, however MOSUM detects a smaller number of change points across all subjects. WBS has the worst performance. The 5-copula family has a marginal superior performance over the Gaussian copula across all segmentation methods of VCCP, but many of the change points align, indicating that the change points are due to changes in the Gaussian copula.

**Figure 7:**
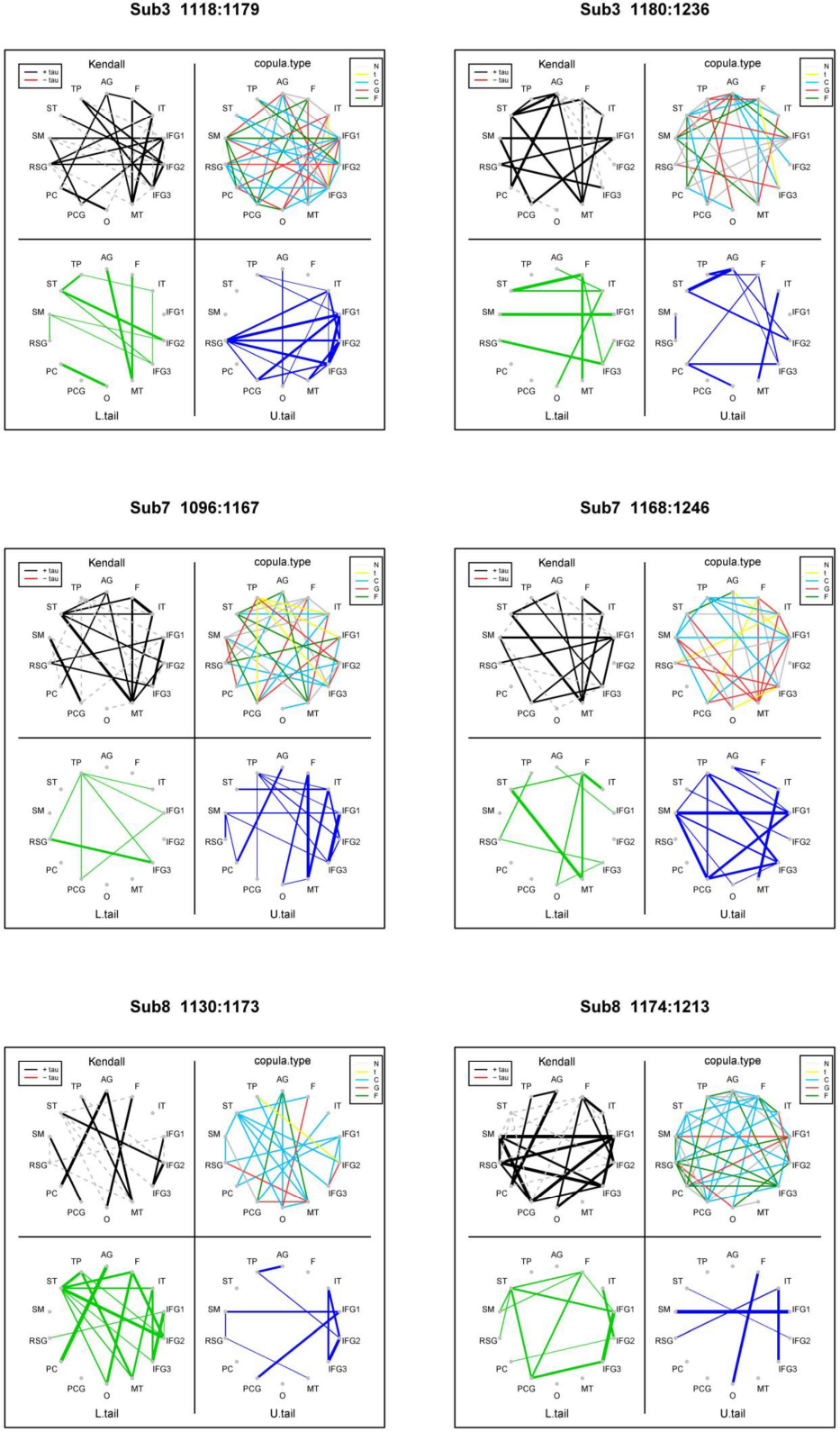
The FC networks estimated using the NBS.V combination of VCCP for subject 3, 7, and 8 at the change point that coincides with the four main characters encountering the three-head dog. The left plots represent the networks before the change point while the right plots represent the networks after the change point. The top left, top right, bottom left and bottom right networks represent the estimated Kendall’s *τ*, the lower tail dependence, the upper tail dependence, and the optimal copula function, for each edge between pairs of nodes, respectively. Black (red) lines in the top left graphs represent positive (negative) Kendall’s *τ* correlation coefficients. Dashed lines indicate edges between nodes with tail dependence but statistically insignificant Kendall’s *τ*. Green and blue edges in the bottom left and right graphs represent the lower and upper tail dependence, respectively. The various colored lines on the top right graphs row indicate the optimal copula family for edges, with grey, yellow, blue, red and green indicating the Gaussian, *t*, Clayton, Gumbel and Frank copula, respectively.

**Figure 8:**
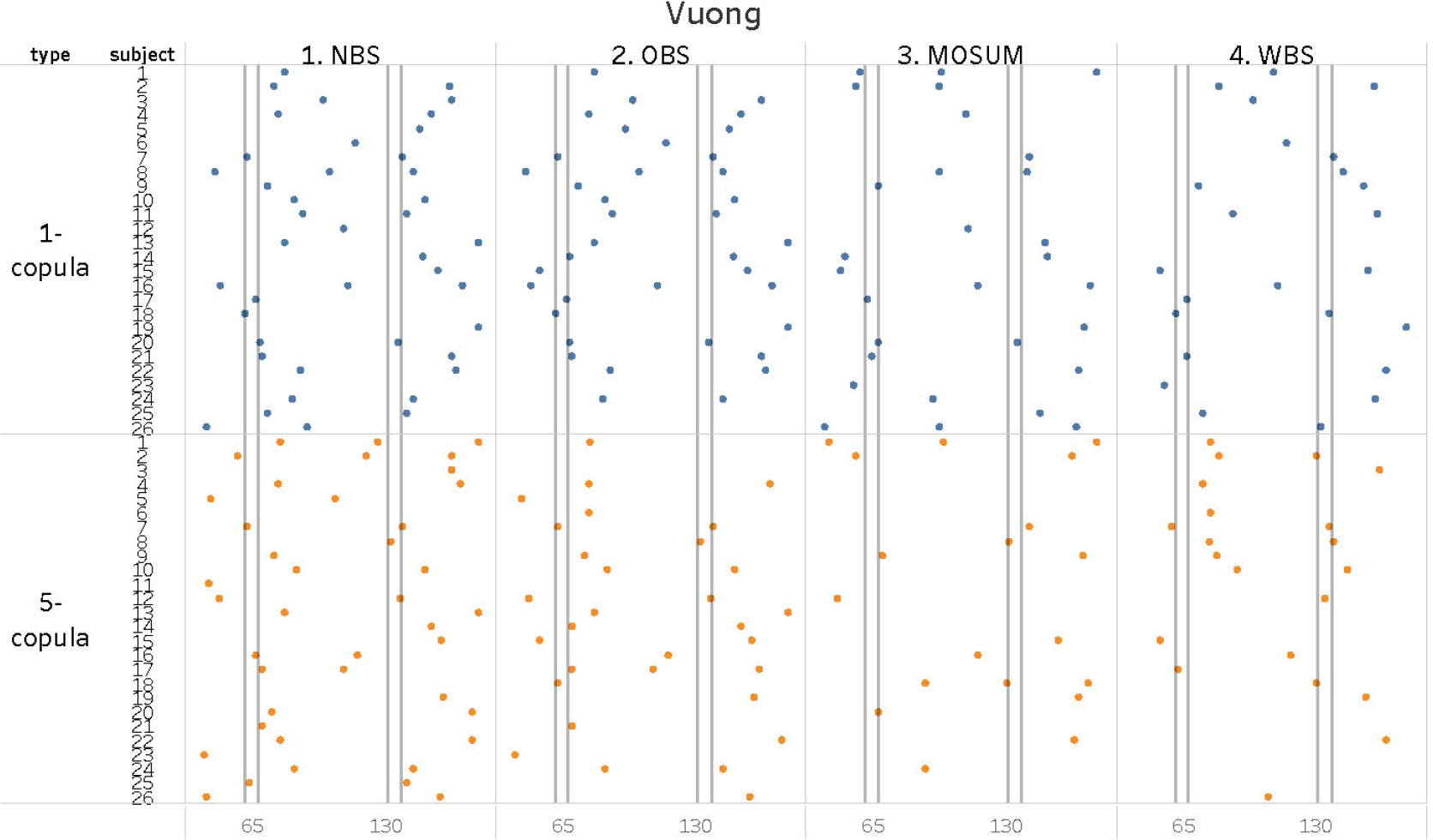
The results from applying all segmentation methods of VCCP (with the Vuong test) to the SET fMRI data set. NBS, OBS, MOSUM, and WBS denote the adapted binary segmentation, old binary segmentation, moving sum, and the wild binary segmentation methods, respectively. We set the minimum distance between change points to be *δ* = 40 and used the Gaussian and the 5-copula family. The solid vertical lines indicate the times of the showing and of the removal of the visual cues.

In Figure 9, we display the results from applying all segmentation methods of VCCP (with the SB) to the SET fMRI data set. The results are very similar to applying the Vuong test (Figure 8). The main difference is that the SB is more computationally intense due to the resampling procedure.

As discussed in Section 2.3 (main file), the parameter *δ*, represents the the minimal distance between two candidate change points. It needs to be sufficiently large enough to estimate a stable D-vine structure, but also small enough in order to not miss candidate change points. In Figure 10, we explore the change points detected from applying the combination NBS.D.V (NBS segmentation, D-vine, and the Vuong test) from VCCP to the SET fMRI data set using values of *δ* = 20, 30, 40. As expected, the number of change points decreases as *δ* increases. This inverse relationship is reasonable as once a candidate point *t*_0_ is detected, we exclude other possible change points in the range [*t*_0_ − *δ, t*_0_ + *δ*] to ensure sufficient sample size during the vine copula construction. Remarkably, even when we explore a limited range (*δ* = 40), NBS can identify change points close to the times of the showing and of the removal of the visual cues. The *δ* = 40 appears to provide and the best and most stable results. The 5-copula family has a superior performance over the Gauss copula across all *δ* values, but many of the change points align, indicating that the change points are due to changes in the Gauss copula. The results from using the SB instead of the Vuong test are very similar.

**Figure 9:**
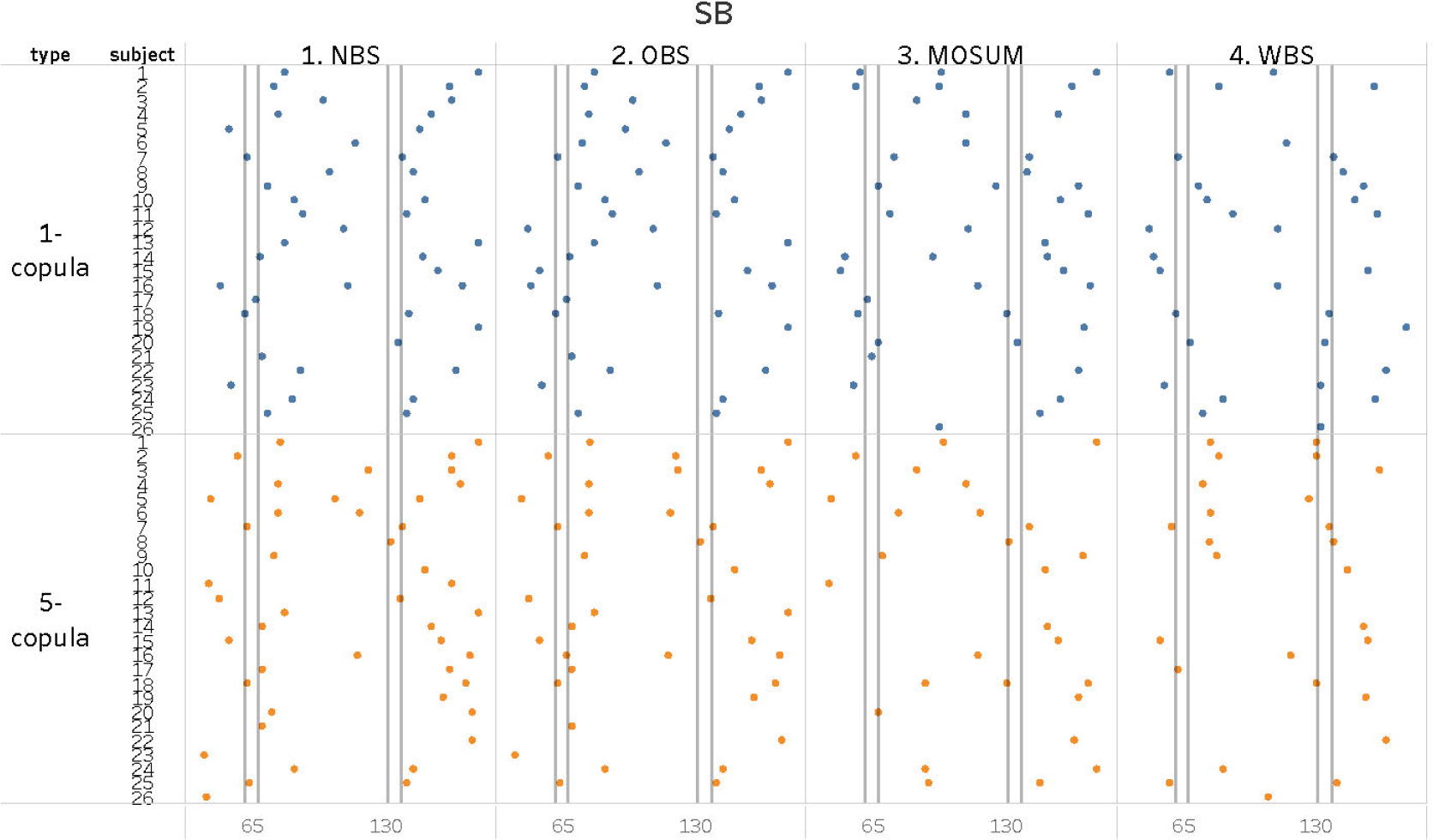
The results from applying all segmentation methods of VCCP (with the SB) to the SET fMRI data set. NBS, OBS, MOSUM, and WBS denote the adapted binary segmentation, old binary segmentation, moving sum, and the wild binary segmentation methods, respectively. We used the Gaussian copula and the 5-copula family. The solid vertical lines indicate the times of the showing and of the removal of the visual cues.

Using the results from Figure 10 (*δ* = 40), we plot the networks for subjects 2, 7 and 25 in the SET data, with change points detected by NBS.D.V and the 5-copula family in Figure 11. The main graph of each partition corresponds to the Kendall’s *τ* network, while the two sub-graphs on the top corners summarize the tail dependence. All partition specific networks (apart from the last partition for subject 7) contain at least an edge with heavy tail dependence, some of them are only displayed in dashed lines in the main graph due to an insignificant Kendall’s *τ*. Since the tail dependence cannot be described by Gaussian models, changes in these non-Kendall-but-heavy-tail correlations may be neglected if researchers build their change point detection model on the Gaussianity assumption. This may be the reason for a sparser change point detection using the Gauss copula than the results from the 5-copula family. For example, in Figure 10 the Gauss copula is only able to detect the two change points for subject 2, while the 5-copula family can detect three change points. For subject 7, the change points using the Gauss copula and the 5-copula family are identical indicating that the change points are due to the Gauss copula with little tail dependence. The middle panel of Figure 11 verifies this.

**Figure 10:**
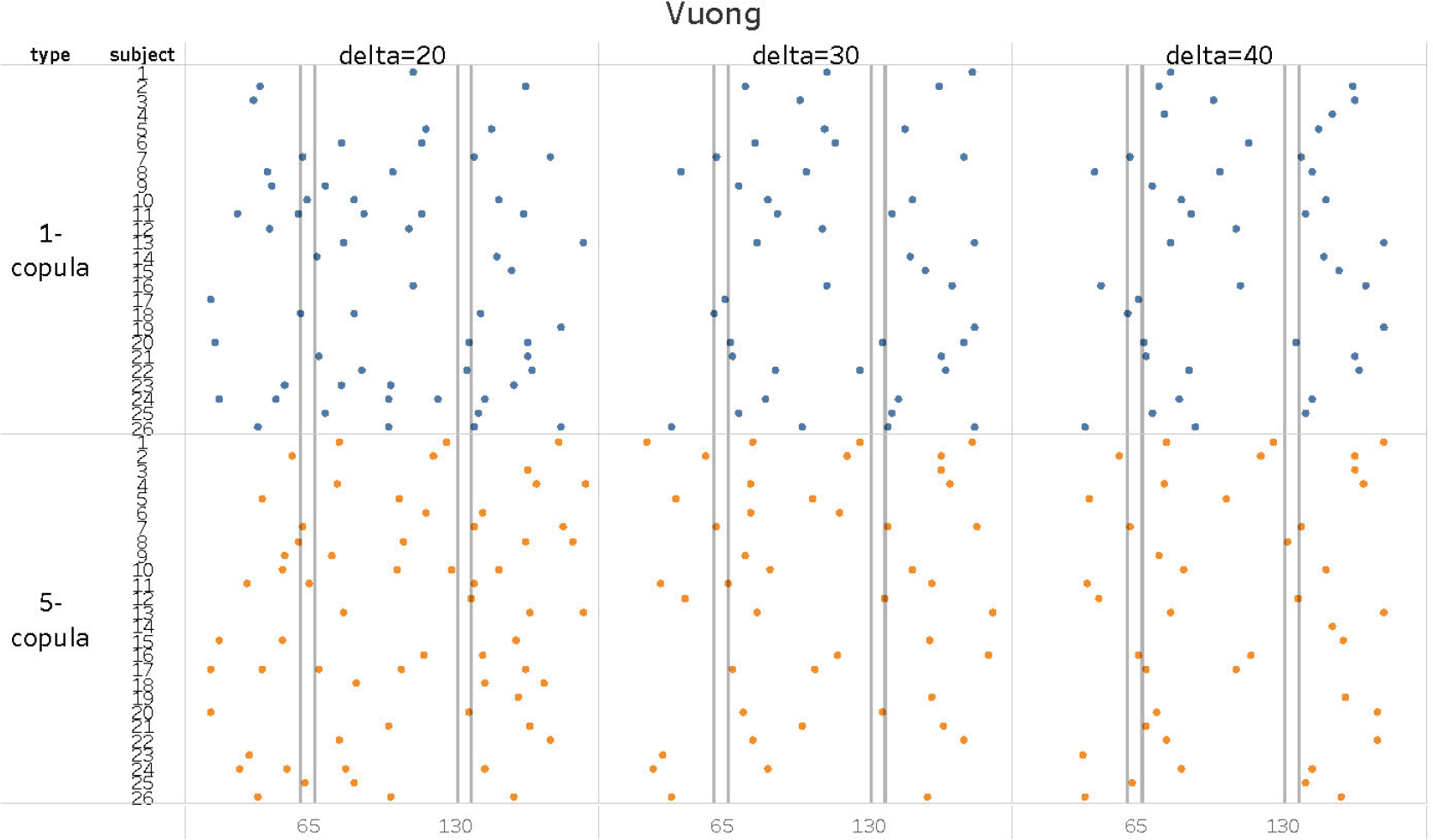
The results from applying the combination NBS.D.V (with Gauss and 5-copula family) from VCCP to the SET fMRI data set. The minimum distance between change points is set to *δ* = 20, 30, 40. The solid vertical lines indicate the times of the showing and of the removal of the visual cues.

In Figure 11, we also label the best copula type for each edge. Only a small portion of them are denoted as Gauss copulas (N), while most of the edges are best fitted with Clayton (C) or Gumbel (G) copulas, consistent with their heavy tail dependence. During different periods, some best-fitting copula types remain the same, such as the F copula linking ventromedial PFC (vPFC) and right superior temporal sulci (R.STS), and Gumbel copula linking the right superior temporal sulci (R.STS) and the left superior temporal sulci (L.STS) in subject 25. However, some edges experience changes not only in connecting strength but also in copula types. The diversity and the variability of copula types in the networks once again prove the superiority of a vine copula model with multiple copula candidates.

**Figure 11:**
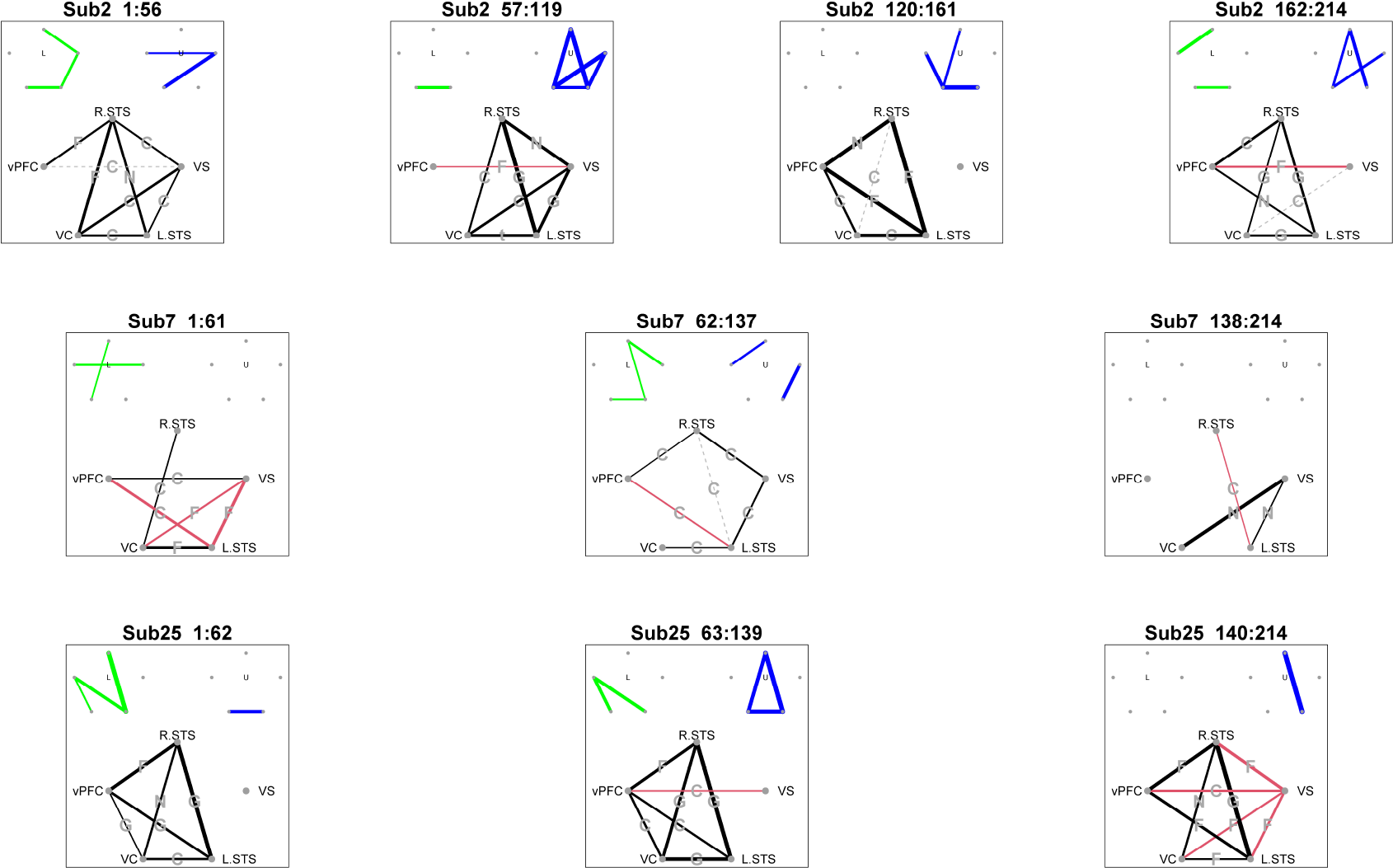
The networks for subjects 2, 7, and 25 in the SET data, with change points detected by NBS.D.V and the 5-copula family. Black and red lines in the main networks correspond to positive and negative Kendall’s *τ*, with a grey label marking the best-fitting copula type for each edge. Only edges that are found to be significant in Kendall’s correlation test or with heavy heavy tail dependence are shown. The green sub-network on the top left corner represents lower tail dependence, and the blue sub-network on the top right corner represents upper tail dependence. Dashed lines indicate ROIs with heavy tail dependence but insignificant Kendall’s *τ*.

Overall, for the anxiety-inducing experiment, the NBS segmentation method performs best with consistent change point detection close to time points (61 − 67.5) and (131 − 137.5). The performance of the SB and the Vuong test are similar. While the 5-copula family allowed for more flexibility in the model and change points in the network, it also increases the computational time.

### 6 Simulation results

#### 6.1 MVN data

Table 3 shows the results from applying all variations of VCCP with 5 different copula types to Simulation 5 (MVN data with 4 change points) over 100 simulated data sequences. Overall, the NBS and OBS segmentation methods shared the best performance with each almost perfectly detecting the four true change points within the bias range for each iteration, especially in combination with SB. NBS.SB and OBS.SB have the smallest scaled Hausdorff distance, indicating the detected change points are located closest to the true change points. MOSUM also present good results, but with a higher FP rate and larger scaled Hausdorff distance. WBS appears to be too conservative with inferior accuracy with both the SB and the Vuong test. Overall, the two best non-parametric methods from the ecp package, e.cp3o and ks.cp3o_delta, have inferior performance to all variations of VCCP with lower TP, higher TN and higher FP (see Table 9). Both e.cp3o and ks.cp3o detected more than 4 change points on average.

Table 4 shows the results from applying all variations of VCCP with 5 different copula types to Simulation 6 (MVN data with 4 weak change points) over 100 simulated data sequences. In this simulation, the nonzero edges are fixed over the entire time course with the edge weights varying between change points. This is a very difficult simulation but for VCCP, the combinations NBS.V, OBS.SB, and WBS.BS perform well, identifying almost all 4 change points across the 100 iterations. Overall, OBS.SB has the best performance, having the smallest scaled Hausdorff distance and the highest TP rate. However, NBS.V has almost the same performance but with a smaller number of FPs. WBS has the next best performance across the segmentation methods, identifying over 3 change points on average, with low FP rates. The MOSUM combinations have the worst performance. In terms of the non-parametric methods, e.cp3o and ks.cp3o_delta perform best (see Table 10). However, they are all inferior to all variations of VCCP, with the FP rates being particularly poor.

We also compare the results in Table 3 (MVN data with 4 change points) with the results in Table 4 (MVN data with 4 weak change points), and find that the results for the latter are inferior due to the less distinctive changing patterns. The number of detected change points declines on average across all variations of VCCP. For the latter data set, the change point located at time point *t* = 61 is the most difficult to detect, where the highest TP rate is 47% from NBS.V. While the FN rate remains low in the former data, it increases for the latter data, especially using MOSUM, where the FN rate for the first partition exceeds 0.3 when combined with both the SB and Voung. Since MOSUM is based on segmenting the reduced BIC series, the soaring FN rate may be due to the selection of the fixed bandwidth *G*. Due to its conservative behavior, WBS performs better in the more difficult simulation. However, it did have low FN rates and high TP rates at the change points at *t* = 121 and *t* = 241 in the latter data set.

**Table 3:** The results from applying all variations of VCCP with 5 different copula types (Gaussian, *t*, Clayton, Gumbel and Frank copulas) to Simulation 5 (MVN data with 4 change points) over 100 simulated data sequences. TP, FN, FP, Cand.num, Num.det, and *d_H_* denote the true positive rate, false negative rate, false positive rate, number of candidate change points, number of change points detected, scaled Hausdorff distance, respectively. N, O, M, and W denote the adapted binary segmentation, old binary segmentation, moving sum, and the wild binary segmentation methods, respectively. ‘5’ denotes the 5 family copula types; ‘B’ and ‘V’ denote the stationary bootstrap and the Vuong test.

|  | N.5.BS | N.5.V | O.5.BS | O.5.V | M.5.BS | M.5.V | W.5.BS | W.5.V |
| --- | --- | --- | --- | --- | --- | --- | --- | --- |
| TP_61 | 1 | 1 | 1 | 0.99 | 0.99 | 0.99 | 0.88 | 0.86 |
| TP_121 | 0.99 | 0.99 | 0.99 | 0.98 | 1 | 1 | 1 | 1 |
| TP_181 | 1 | 1 | 1 | 0.99 | 1 | 1 | 0.97 | 0.97 |
| TP_241 | 1 | 0.98 | 1 | 0.99 | 0.99 | 0.97 | 0.74 | 0.74 |
| FN_61 | 0 | 0 | 0 | 0 | 0 | 0 | 0 | 0.02 |
| FN_121 | 0 | 0 | 0 | 0 | 0 | 0 | 0 | 0 |
| FN_181 | 0 | 0 | 0 | 0 | 0 | 0 | 0 | 0 |
| FN_241 | 0 | 0.02 | 0 | 0.01 | 0 | 0.02 | 0.01 | 0.01 |
| FP_1-60 | 0 | 0.01 | 0 | 0 | 0.01 | 0.07 | 0.02 | 0.02 |
| FP_61-120 | 0.03 | 0.09 | 0.04 | 0.08 | 0.02 | 0.03 | 0.03 | 0.03 |
| FP_121-180 | 0 | 0.05 | 0.01 | 0.04 | 0.01 | 0.01 | 0 | 0 |
| FP_181-240 | 0 | 0.05 | 0 | 0.05 | 0.05 | 0.08 | 0.02 | 0.02 |
| FP_241-299 | 0 | 0.09 | 0.01 | 0.08 | 0.03 | 0.07 | 0.04 | 0.04 |
| Hausdorff | 0.06 | 0.16 | 0.07 | 0.17 | 0.12 | 0.16 | 0.37 | 0.39 |
| Cand.num | 5.24 | 5.24 | 4.86 | 4.92 | 4.31 | 4.31 | 3.75 | 3.75 |
| Num.det | 4.02 | 4.26 | 4.05 | 4.2 | 4.08 | 4.2 | 3.68 | 3.66 |
| time (s) | 1039.97 | 633.6 | 1162.98 | 634.04 | 480.45 | 104.52 | 2774.43 | 2482.63 |

**Table 4:** The results from applying all variations of VCCP with 5 different copula types (Gaussian, *t*, Clayton, Gumbel and Frank copulas) to Simulation 6 (MVN data with 4 weak change points) over 100 simulated data sequences. TP, FN, FP, Cand.num, Num.det, and *d_H_* denote the true positive rate, false negative rate, false positive rate, number of candidate change points, number of change points detected, scaled Hausdorff distance, respectively. N, O, M, and W denote the adapted binary segmentation, old binary segmentation, moving sum, and the wild binary segmentation methods, respectively. ‘5’ denotes the 5 family copula types; ‘B’ and ‘V’ denote the stationary bootstrap and the Vuong test.

|  | N.5.BS | N.5.V | O.5.BS | O.5.V | M.5.BS | M.5.V | W.5.BS | W.5.V |
| --- | --- | --- | --- | --- | --- | --- | --- | --- |
| TP_61 | 0.33 | 0.47 | 0.43 | 0.28 | 0.29 | 0.33 | 0.4 | 0.33 |
| TP_121 | 0.72 | 0.81 | 0.86 | 0.57 | 0.7 | 0.75 | 0.87 | 0.82 |
| TP_181 | 0.38 | 0.45 | 0.63 | 0.33 | 0.41 | 0.46 | 0.54 | 0.43 |
| TP_241 | 0.85 | 0.83 | 0.92 | 0.51 | 0.83 | 0.82 | 0.87 | 0.79 |
| FN_61 | 0.36 | 0.22 | 0.09 | 0.07 | 0.36 | 0.32 | 0.14 | 0.11 |
| FN_121 | 0.18 | 0.09 | 0 | 0.02 | 0.14 | 0.09 | 0 | 0.02 |
| FN_181 | 0.42 | 0.35 | 0.11 | 0.05 | 0.23 | 0.18 | 0.09 | 0.14 |
| FN_241 | 0.09 | 0.11 | 0.01 | 0.2 | 0.07 | 0.08 | 0 | 0.07 |
| FP_1-60 | 0.11 | 0.19 | 0.19 | 0.06 | 0.11 | 0.07 | 0.05 | 0.03 |
| FP_61-120 | 0.11 | 0.24 | 0.26 | 0.08 | 0.05 | 0.1 | 0.14 | 0.1 |
| FP_121-180 | 0.12 | 0.19 | 0.25 | 0.1 | 0.09 | 0.15 | 0.15 | 0.11 |
| FP_181-240 | 0.07 | 0.11 | 0.15 | 0.07 | 0.07 | 0.06 | 0.08 | 0.08 |
| FP_241-299 | 0.08 | 0.13 | 0.11 | 0.04 | 0.06 | 0.14 | 0.01 | 0.01 |
| Hausdorff | 1.29 | 0.85 | 0.63 | 1.98 | 1.27 | 1.08 | 0.75 | 1.1 |
| Cand.num | 5.94 | 5.94 | 4.65 | 2.94 | 4.48 | 4.48 | 3.57 | 3.27 |
| Num.det | 2.7 | 3.32 | 3.69 | 2 | 2.6 | 2.84 | 3.06 | 2.66 |
| time (s) | 838.8 | 457.95 | 846.82 | 332.12 | 396.13 | 80.6 | 1835.96 | 1619.27 |

Table 5 shows the results from applying all variations of VCCP with 5 different copula types to Simulation 7 (MVN data with 7 small change points) over 100 simulated data sequences. For VCCP, NBS.V and OBS.SB have the best performance, with OBS.SB marginally outperforming NBS.V. MOSUM and WBS have a similar performance in terms of scaled Hausdorff distance with MOSUM detecting more change points on average. The change point at time point *t* = 526 and *t* = 301 were difficult to detect. With respect to the non-parametric methods, all variations of VCCP outperform them. In particular, for this data set, the best performing methods, e.cp3o_delta E.cp3o, have low TP, high FN and very high FP rates (see Table 11).

#### 6.2 VAR results

Table 6 shows the results from applying all variations of VCCP with 5 different copula types to Simulation 8 (VAR data with 4 change points) over 100 simulated data sequences. For VCCP, NBS.SB has the best performance, it perfectly recognizes the four true change points within the bias range for each iteration, with a small FP rate. Similar to the corresponding MVN simulation, NBS.SB and OBS.SB also have the smallest scaled Hausdorff distance, indicating the detected change points are located closest to the true change points. They also have low FN and FP rates. The results for the Vuong test and the SB are very similar in combination with all the segmentation methods, with Voung having superior computational speed. Other combinations such as MOSUM present good results, but with slightly higher FP rates. WBS appears to be too conservative with inferior accuracy but smaller FP rates. Overall, e.cp3o and ks.cp3o have inferior performance to all variations of VCCP with lower TP, higher TN and higher FP (see Table 12). e.cp3o and ks.cp3o show similar results, with ecp being more conservative, leading to lower TP as well as lower FP and FN rates. Both detected more than 4 change points on average. If we reduce the *Kmax* to 4 (see Table 12), all methods in ecp detected fewer than 3 change points.

Table 7 shows the results from applying all variations of VCCP with 5 different copula types to Simulation 8 (VAR data with 4 change points) over 100 simulated data sequences. In this simulation, only the edge weights in the network are fluctuating between change points with the non-zero edges themselves remaining fixed over the entire time course. This is a very difficult simulation but for VCCP with 5 different copula types, WBS.BS performs well in terms of detection, low FP rate, and low Hausdorff distance. NBS.V, WBS.B and NBS.BS also have a decent performance. OBS.V has the worst performance, identifying the least number of true change points. Since OBS carries out segmentation and inference sequentially, we attribute its deterioration in performance to this. Once the changes become weak enough, it is likely that the Vuong test reject the first candidate in OBS, which results in it failing to detect the remaining change points. As small changes are common in resting-state fMRI data, it might be inappropriate to apply OBS to this data type. The good performance of WBS may stem from the sub-sampling step in WBS. As stated in Section 2, WBS firstly creates a series of multiple candidates and with their corresponding BIC reductions within each sub-sample. Then an OBS-like inference procedure decides the test order. Compared to the BIC reduction of the original OBS method, the sub-sampling idea in WBS overcomes its drawbacks. In fact, WBS is more similar to the combination of NBS and OBS, which contributes to its unique strength. In terms of the non-parametric methods, e.cp3o and ks.cp3o_delta perform best (see Table 13). However, they are all inferior to all variations of VCCP. In fact, it becomes hard to pick the “best” nonparametric methods for this simulation, given the variation in TP and FP. When *Kmax* = 4, all but e.divisive have low TP and FP rates. When *Kmax* increases to 8, the TP rate grows, but the FP rate of e.cp3o_delta soars to 1 in some intervals, and the same applies to other methods requiring *Kmax* in their detection. This pattern indicates non-robustness of the non-parametric methods. On the contrary, methods depending on the E-statistics are less sensitive with respect to the *α* value.

**Table 5:**
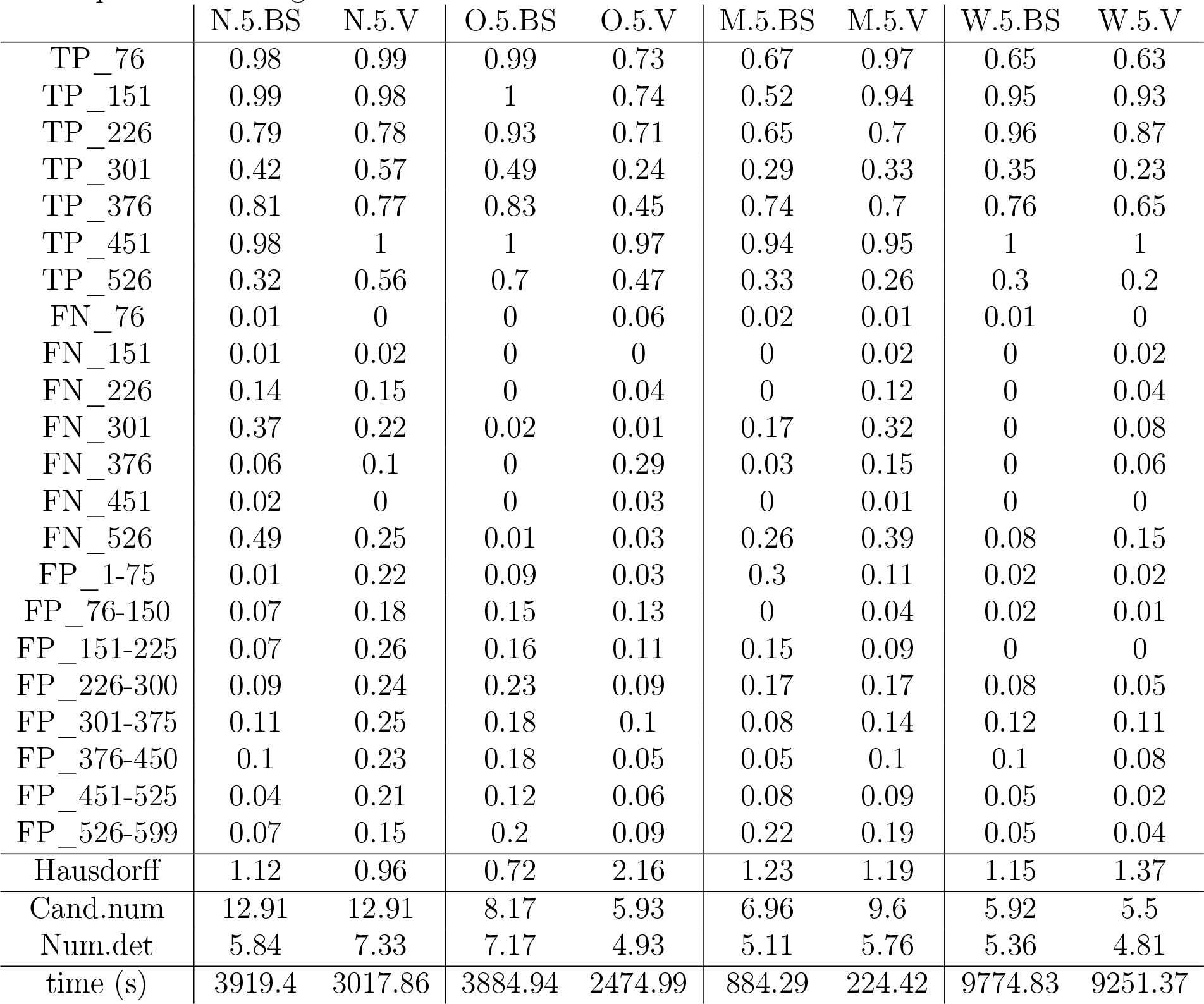
The results from applying all variations of VCCP with 5 different copula types (Gaussian, *t*, Clayton, Gumbel and Frank copulas) to Simulation 7 (MVN data with 7 small change points) over 100 simulated data sequences. TP, FN, FP, Cand.num, Num.det, and *d_H_* denote the true positive rate, false negative rate, false positive rate, number of candidate change points, number of change points detected, scaled Hausdorff distance, respectively. N, O, M, and W denote the adapted binary segmentation, old binary segmentation, moving sum, and the wild binary segmentation methods, respectively. ‘5’ denotes the 5 family copula types; ‘B’ and ‘V’ denote the stationary bootstrap and the Vuong test.

**Table 6:** The results from applying all variations of VCCP with 5 different copula types (Gaussian, *t*, Clayton, Gumbel and Frank copulas) to Simulation 8 (VAR data with 4 change points) over 100 simulated data sequences. TP, FN, FP, Cand.num, Num.det, and *d_H_* denote the true positive rate, false negative rate, false positive rate, number of candidate change points, number of change points detected, scaled Hausdorff distance, respectively. N, O, M, and W denote the adapted binary segmentation, old binary segmentation, moving sum, and the wild binary segmentation methods, respectively. ‘5’ denotes the 5 family copula types; ‘BS’ and ‘V’ denote the stationary bootstrap and the Vuong test.

|  | N.5.BS | N.5.V | O.5.BS | O.5.V | M.5.BS | M.5.V | W.5.BS | W.5.V |
| --- | --- | --- | --- | --- | --- | --- | --- | --- |
| TP_61 | 1 | 0.98 | 1 | 0.99 | 1 | 1 | 0.83 | 0.8 |
| TP_121 | 0.99 | 1 | 1 | 0.99 | 1 | 0.99 | 0.97 | 0.96 |
| TP_181 | 0.98 | 0.95 | 0.99 | 0.98 | 0.94 | 0.95 | 0.9 | 0.86 |
| TP_241 | 0.97 | 0.93 | 0.99 | 0.98 | 0.9 | 0.89 | 0.67 | 0.63 |
| FN_61 | 0 | 0.02 | 0 | 0 | 0 | 0 | 0 | 0.03 |
| FN_121 | 0.01 | 0 | 0 | 0 | 0 | 0.01 | 0 | 0 |
| FN_181 | 0.01 | 0.04 | 0 | 0 | 0.01 | 0 | 0 | 0.03 |
| FN_241 | 0.02 | 0.06 | 0 | 0.01 | 0.03 | 0.04 | 0.1 | 0.12 |
| FP_1-60 | 0.01 | 0.09 | 0.02 | 0.08 | 0.09 | 0.12 | 0.02 | 0.02 |
| FP_61-120 | 0 | 0.06 | 0.01 | 0.05 | 0.01 | 0.01 | 0.05 | 0.03 |
| FP_121-180 | 0.03 | 0.07 | 0.04 | 0.08 | 0.04 | 0.05 | 0.04 | 0.02 |
| FP_181-240 | 0.04 | 0.14 | 0.07 | 0.17 | 0.09 | 0.07 | 0.01 | 0.02 |
| FP_241-299 | 0.01 | 0.02 | 0.01 | 0.04 | 0.05 | 0.08 | 0.01 | 0.03 |
| Hausdorff | 0.12 | 0.28 | 0.11 | 0.23 | 0.21 | 0.24 | 0.53 | 0.63 |
| Cand.num | 5.57 | 5.57 | 5.02 | 5.12 | 4.55 | 4.55 | 3.74 | 3.65 |
| Num.det | 4 | 4.21 | 4.1 | 4.33 | 4.12 | 4.17 | 3.49 | 3.34 |
| time (s) | 866.88 | 506.29 | 959.75 | 504.8 | 414.06 | 84.92 | 2134.22 | 1880.08 |

We also compared the results in Table 6 (VAR data with 4 change points) with the results in Table 7 (VAR data with 4 weak change points), and find that the results for the latter are inferior due to the less distinctive changing patterns. Similar to the MVN data, the performance degraded from the former data set to the latter. The FP rate and the scaled Hausdorff distance increased. Similar to the results in the MVN data with 4 weak change points, the first change point located at time point *t* = 61 is the most difficult to detect, where the highest TP rate was only 49% from WBS.BS.

**Table 7:** The results from applying all variations of VCCP with 5 different copula types (Gaussian, *t*, Clayton, Gumbel and Frank copulas) to Simulation 9 (VAR data with 4 weak change points) over 100 simulated data sequences. TP, FN, FP, Cand.num, Num.det, and *d_H_* denote the true positive rate, false negative rate, false positive rate, number of candidate change points, number of change points detected, scaled Hausdorff distance, respectively. N, O, M, and W denote the adapted binary segmentation, old binary segmentation, moving sum, and the wild binary segmentation methods, respectively. ‘5’ denotes the 5 family copula types; ‘BS’ and ‘V’ denote the stationary bootstrap and the Vuong test.

|  | N.5.BS | N.5.V | O.5.BS | O.5.V | M.5.BS | M.5.V | W.5.BS | W.5.V |
| --- | --- | --- | --- | --- | --- | --- | --- | --- |
| TP_61 | 0.24 | 0.3 | 0.14 | 0.08 | 0.1 | 0.15 | 0.49 | 0.3 |
| TP_121 | 0.46 | 0.51 | 0.4 | 0.21 | 0.3 | 0.36 | 0.69 | 0.52 |
| TP_181 | 0.58 | 0.58 | 0.57 | 0.24 | 0.52 | 0.52 | 0.84 | 0.7 |
| TP_241 | 0.7 | 0.66 | 0.59 | 0.3 | 0.71 | 0.71 | 0.8 | 0.71 |
| FN_61 | 0.3 | 0.24 | 0.14 | 0.11 | 0.55 | 0.5 | 0.1 | 0.15 |
| FN_121 | 0.27 | 0.22 | 0.03 | 0.09 | 0.3 | 0.24 | 0.03 | 0.13 |
| FN_181 | 0.26 | 0.26 | 0.02 | 0.05 | 0.24 | 0.24 | 0.02 | 0.12 |
| FN_241 | 0.11 | 0.15 | 0.04 | 0.11 | 0.11 | 0.11 | 0.03 | 0.07 |
| FP_1-60 | 0.07 | 0.16 | 0.15 | 0.03 | 0 | 0.02 | 0.06 | 0.04 |
| FP_61-120 | 0.15 | 0.18 | 0.14 | 0.09 | 0.08 | 0.12 | 0.15 | 0.09 |
| FP_121-180 | 0.12 | 0.13 | 0.21 | 0.12 | 0.08 | 0.11 | 0.21 | 0.14 |
| FP_181-240 | 0.08 | 0.11 | 0.11 | 0.05 | 0.03 | 0.06 | 0.07 | 0.04 |
| FP_241-299 | 0.17 | 0.21 | 0.16 | 0.07 | 0.1 | 0.1 | 0.02 | 0.03 |
| Hausdorff | 1.58 | 1.3 | 1.85 | 2.84 | 2.06 | 1.77 | 0.71 | 1.34 |
| Cand.num | 5.44 | 5.44 | 3.3 | 2.1 | 4.38 | 4.38 | 3.62 | 3.16 |
| Num.det | 2.52 | 2.78 | 2.42 | 1.15 | 1.89 | 2.14 | 3.24 | 2.49 |
| time (s) | 858.83 | 521.84 | 748.17 | 333.72 | 393.45 | 79 | 1839.76 | 1607.59 |

Table 8 shows the results from applying all variations of VCCP with 5 different copula types to Simulation 10 (VAR data with 7 small change points) over 100 simulated data sequences. For VCCP, NBS.V has the best performance with a high TP rate and the lowest Hausdorff distance. MOSUM and WBS do not perform well in this simulation. OBS.V has the worst performance, identifying the least number of true change points and having the largest Hausdorff distance by some distance.

To explore the reason behind its unusual performance, we notice that in OBS, the inference result at the beginning of the procedure decides whether to conduct later segmentation. If the first candidate change point is at time point *t*, the Vuong test compares the goodness of fit between *V C*_0_ and *V C_l_* using the left part of data (1 : *t* − 1), and *V C*_0_ and *V C_r_* using the right part of data (*t* : *T*). Only when both tests show *V C*_0_ is inferior do we treat *t* as a change point. However, if one candidate is close to the beginning (end) of the timeline and the right (left) part of data contains one or more latent change points, segmentation at time point *t* might not improve the goodness-of-fit of the right (left) part of data. That is to say, neither *V C_r_* (*V C_l_*) nor *V C*_0_ can perfectly describe the distribution of the right (left) part of data with the latent change points undetected. In some cases, the Vuong test even provides a reverse result that *V C*_0_ outperforms the other VC model due to a larger sample size. This happens more frequently when the changes in the structure are less discernible. Therefore, the candidate change point at time point *t* is not considered a change point, and other latent change points remain undiscovered because the segmentation terminates, as shown in the left panel of Figure 12. Here, for one iteration of Simulation 7, OBS.V stops at the second candidate change point (*t* = 76), even though the left part of the test (the dark blue dot) has identified it as a change point. On the contrary, NBS.V’s performance is superior as it performs the Vuong test on all candidate change points after its exhaustive search.

With respect to the non-parametric methods, we observe similar findings. Apart from OBS combined with Vuong, all variations of VCCP outperform them (see Table 14). In particular, for this data set, the performance of e.cp3o is poor. E.cp3o_delta has the best performance for all *α* values, yet it still suffers from high FP and FN rates. Additionally, the predominant effect of *Kmax* in Num.det still exists, deteriorating the model’s robustness. VCCP, on the contrary, does not have this issue since the number of candidate change points is completely data-driven without any prior knowledge.

**Table 8:** The results from applying all variations of VCCP with 5 different copula types (Gaussian, *t*, Clayton, Gumbel and Frank copulas) to Simulation 10 (VAR data with 7 small change points) over 100 simulated data sequences. TP, FN, FP, Cand.num, Num.det, and *d_H_* denote the true positive rate, false negative rate, false positive rate, number of candidate change points, number of change points detected, scaled Hausdorff distance, respectively. N, O, M, and W denote the adapted binary segmentation, old binary segmentation, moving sum, and the wild binary segmentation methods, respectively. ‘5’ denotes the 5 family copula types; ‘BS’ and ‘V’ denote the stationary bootstrap and the Vuong test.

|  | N.5.BS | N.5.V | O.5.BS | O.5.V | M.5.BS | M.5.V | W.5.BS | W.5.V |
| --- | --- | --- | --- | --- | --- | --- | --- | --- |
| TP_76 | 0.95 | 0.93 | 0.96 | 0.22 | 0.67 | 0.96 | 0.59 | 0.57 |
| TP_151 | 0.99 | 0.97 | 0.99 | 0.22 | 0.75 | 0.98 | 0.96 | 0.96 |
| TP_226 | 0.76 | 0.82 | 0.94 | 0.17 | 0.46 | 0.72 | 0.43 | 0.32 |
| TP_301 | 0.71 | 0.77 | 0.89 | 0.18 | 0.5 | 0.62 | 0.59 | 0.54 |
| TP_376 | 0.88 | 0.91 | 0.91 | 0.18 | 0.64 | 0.74 | 0.85 | 0.81 |
| TP_451 | 0.92 | 0.91 | 0.95 | 0.4 | 0.85 | 0.88 | 0.87 | 0.86 |
| TP_526 | 0.62 | 0.68 | 0.81 | 0.25 | 0.44 | 0.45 | 0.3 | 0.25 |
| FN_76 | 0.01 | 0.03 | 0 | 0.19 | 0.01 | 0.03 | 0 | 0.02 |
| FN_151 | 0 | 0.02 | 0 | 0.02 | 0 | 0.02 | 0 | 0 |
| FN_226 | 0.2 | 0.14 | 0.01 | 0.02 | 0.05 | 0.1 | 0.08 | 0.15 |
| FN_301 | 0.23 | 0.17 | 0.01 | 0.02 | 0.05 | 0.17 | 0.06 | 0.06 |
| FN_376 | 0.04 | 0.01 | 0.01 | 0.2 | 0.01 | 0.11 | 0 | 0.01 |
| FN_451 | 0.04 | 0.05 | 0.01 | 0.14 | 0 | 0.04 | 0 | 0.01 |
| FN_526 | 0.23 | 0.17 | 0.01 | 0.11 | 0.12 | 0.24 | 0.14 | 0.1 |
| FP_1-75 | 0.07 | 0.23 | 0.15 | 0.04 | 0.34 | 0.08 | 0.03 | 0.02 |
| FP_76-150 | 0.05 | 0.17 | 0.18 | 0.02 | 0 | 0.03 | 0.04 | 0.01 |
| FP_151-225 | 0.03 | 0.1 | 0.09 | 0.02 | 0.11 | 0.13 | 0.05 | 0 |
| FP_226-300 | 0.03 | 0.17 | 0.17 | 0.03 | 0.1 | 0.16 | 0.09 | 0.08 |
| FP_301-375 | 0.12 | 0.21 | 0.21 | 0.05 | 0.2 | 0.31 | 0.15 | 0.13 |
| FP_376-450 | 0.09 | 0.2 | 0.15 | 0.03 | 0.08 | 0.16 | 0.02 | 0.02 |
| FP_451-525 | 0.16 | 0.29 | 0.28 | 0.05 | 0.09 | 0.11 | 0.05 | 0.05 |
| FP_526-599 | 0.03 | 0.19 | 0.11 | 0.02 | 0.27 | 0.21 | 0.07 | 0.04 |
| Hausdorff | 0.83 | 0.81 | 0.41 | 6.2 | 1.25 | 1.05 | 1.24 | 1.31 |
| Cand.num | 13.59 | 13.59 | 8.76 | 2.88 | 6.62 | 9.61 | 5.63 | 5.34 |
| Num.det | 6.37 | 7.56 | 7.76 | 1.88 | 5.4 | 6.46 | 5.06 | 4.63 |
| time (s) | 3584.47 | 2707.79 | 3694.8 | 1375.93 | 809.62 | 204.73 | 8230.39 | 7811 |

**Figure 12:**
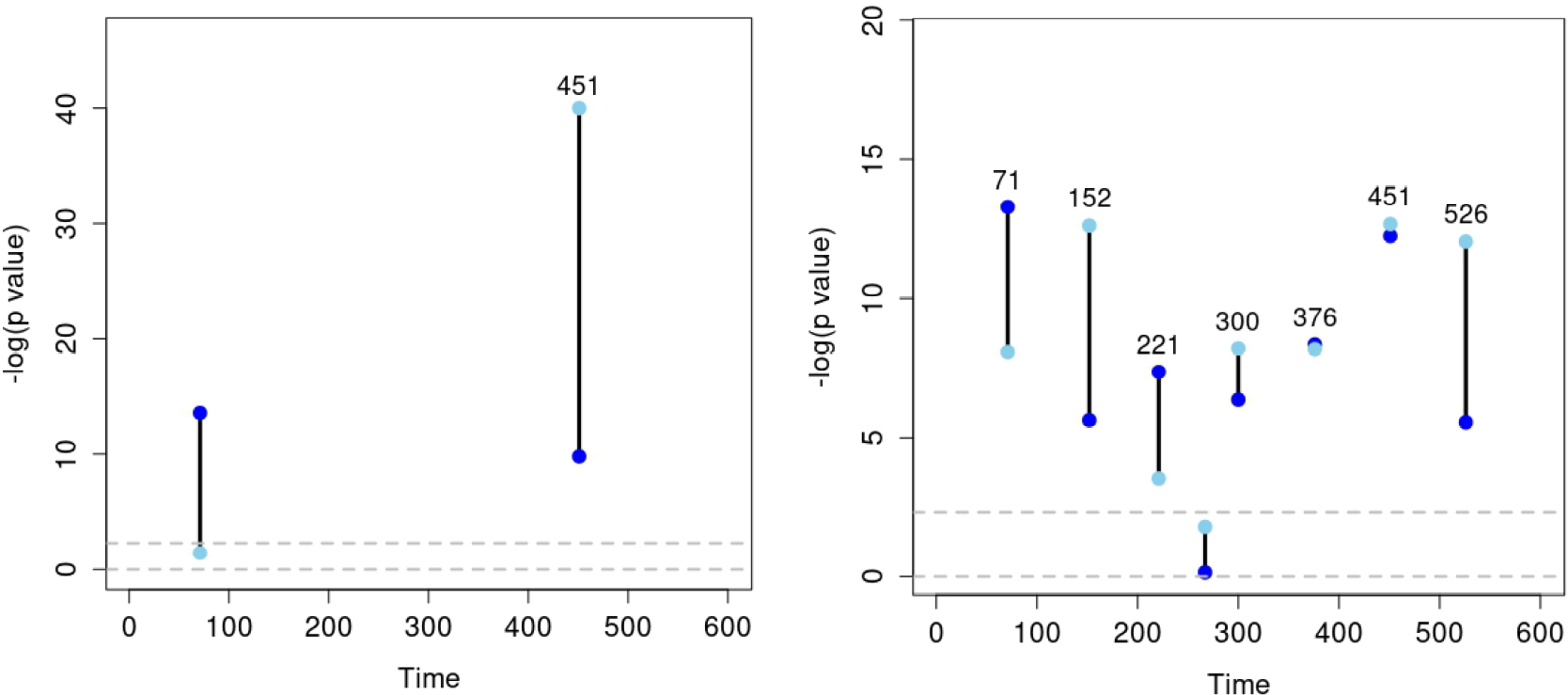
A comparison of the OBS.D.V and NBS.D.V combination methods when applied to one iteration of the VAR data with 7 small change points simulation. The y-axis represents the log(*p*-value) from the Vuong test. The horizontal dashed line represents the significance level, *α* = 0.1^*∗*^log(*p*-value)= 2.303. The dark blue and light blue dots represent the test using the left part and the right part of the data, respectively.

We also put the performance of all the non-parametric methods from Simulation 5 to Simulation 10 in table 9, 10, 11, 12, 13, and 14, respectively.

**Table 9:**
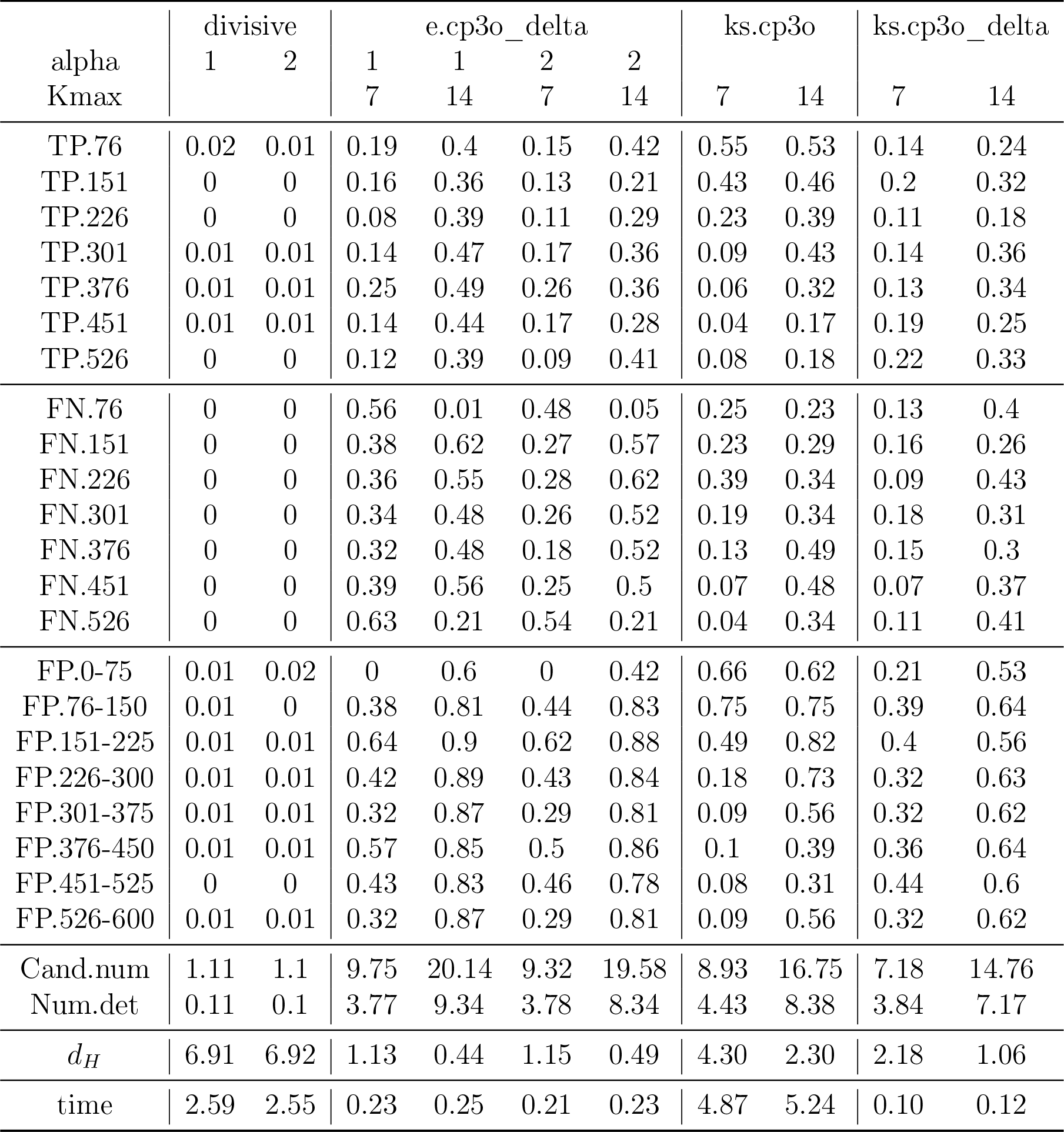
The results from applying 5 non-parametric methods in the **ecp** R package to the MVN data with 4 change points (Simulation 5) over 100 simulated data sequences. TP, FN, FP, Cand.num, Num.det, and *d_H_* denote the true positive rate, false negative rate, false positive rate, number of candidate change points, number of change points detected, scaled Hausdorff distance, respectively.

**Table 10:**
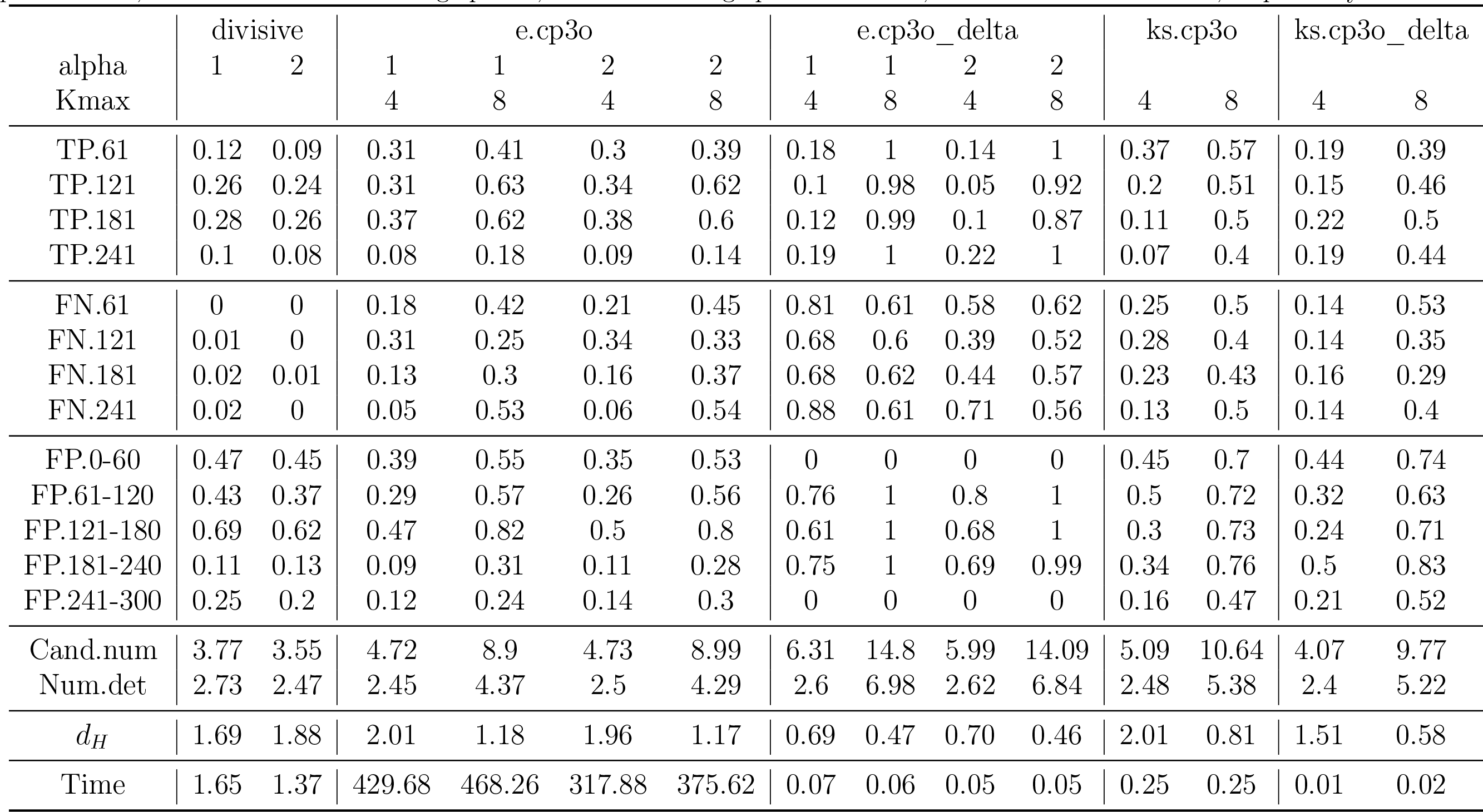
smallThe results from applying 5 non-parametric methods in the **ecp** R package to the MVN data with 4 weak change points (Simulation 6) over 100 simulated data sequences. TP, FN, FP, Cand.num, Num.det, and *d_H_* denote the true positive rate, false negative rate, false positive rate, number of candidate change points, number of change points detected, scaled Hausdorff distance, respectively.

**Table 11:**
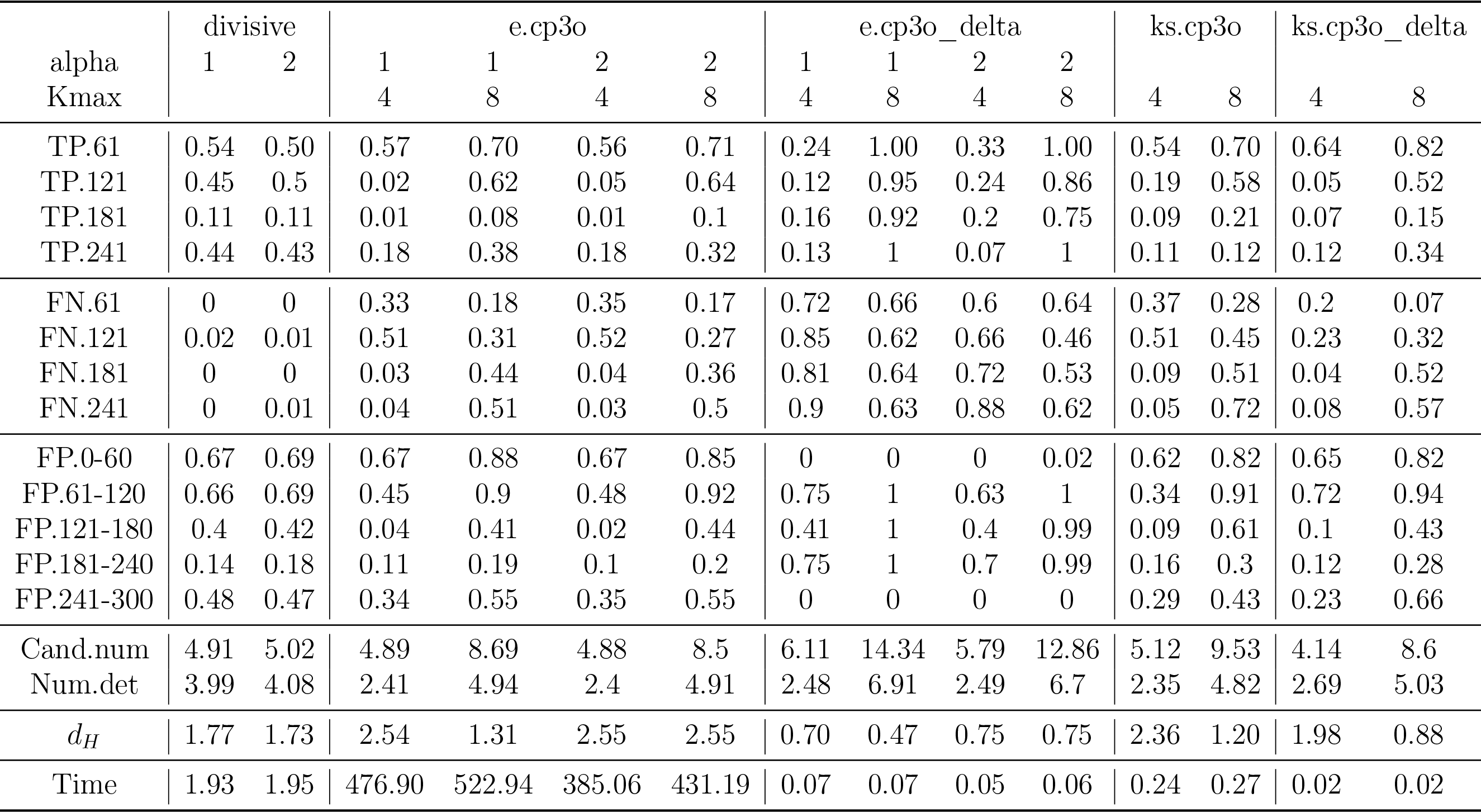
smallThe results from applying 4 non-parametric methods in the **ecp** R package to the MVN data with 7 small change points (Simulation 7) over 100 simulated data sequences. TP, FN, FP, Cand.num, Num.det, and *d_H_* denote the true positive rate, false negative rate, false positive rate, number of candidate change points, number of change points detected, scaled Hausdorff distance, respectively.

**Table 12:**
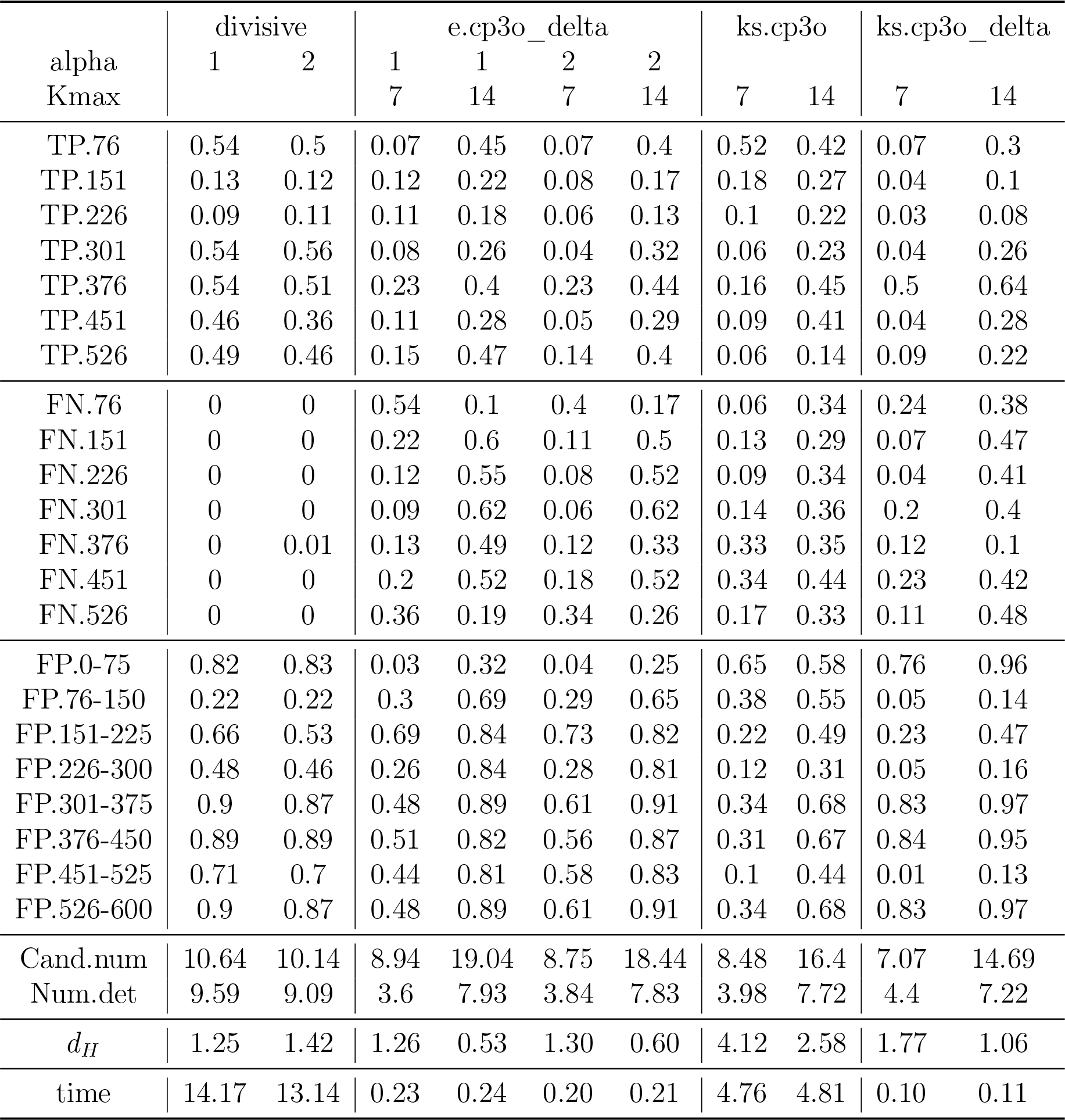
The results from applying 5 non-parametric methods in the **ecp** R package to the VAR data with 4 change points (Simulation 8) over 100 simulated data sequences. TP, FN, FP, Cand.num, Num.det, and *d_H_* denote the true positive rate, false negative rate, false positive rate, number of candidate change points, number of change points detected, scaled Hausdorff distance, respectively.

**Table 13:**
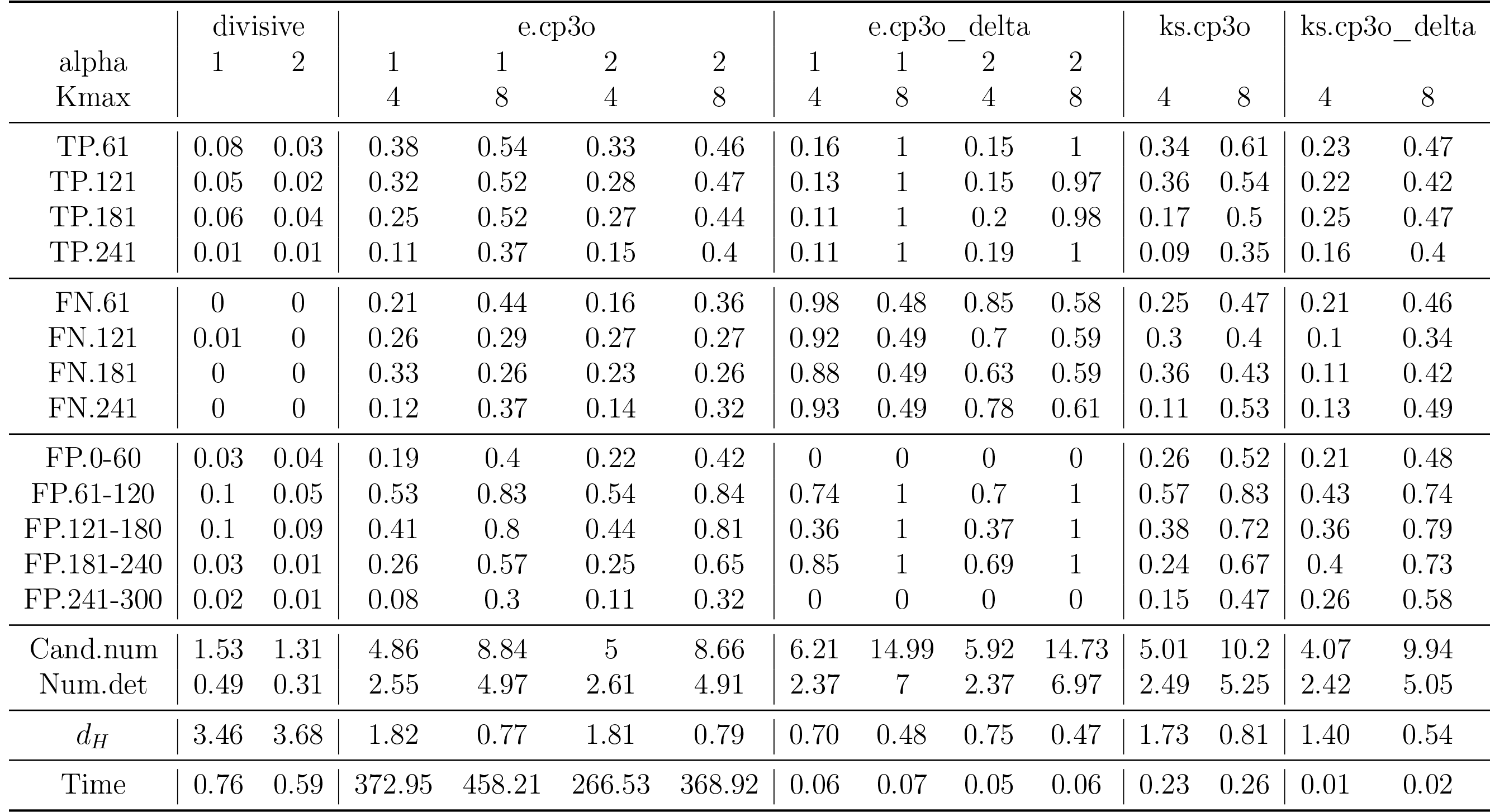
The results from applying 5 non-parametric methods in the **ecp** R package to the VAR data with 4 weak change points (Simulation 9) over 100 simulated data sequences. TP, FN, FP, Cand.num, Num.det, and *d_H_* denote the true positive rate, false negative rate, false positive rate, number of candidate change points, number of change points detected, scaled Hausdorff distance, respectively.

**Table 14:**
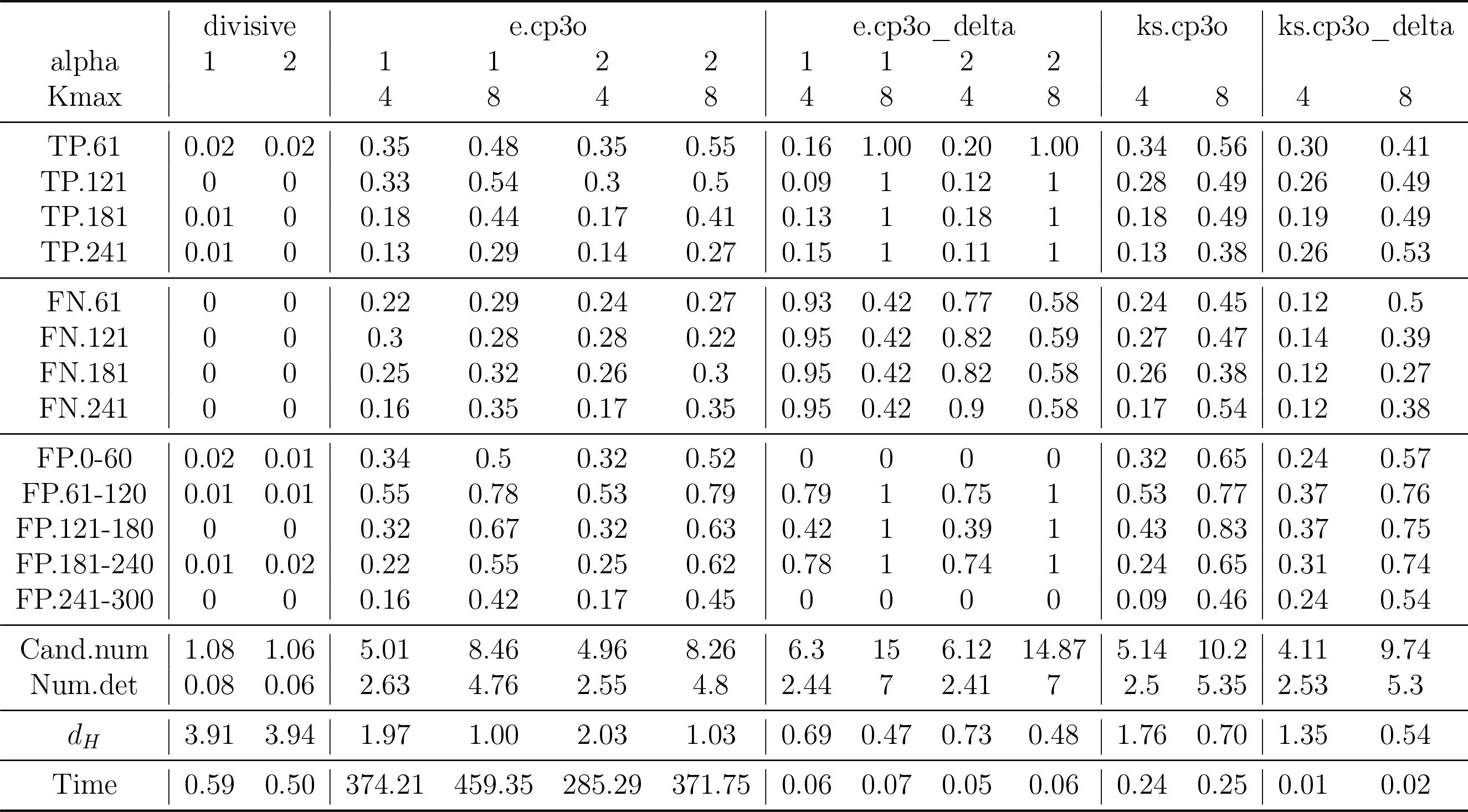
The results from applying 4 non-parametric methods in the **ecp** R package to the VAR data with 7 small change points (Simulation 10) over 100 simulated data sequences. TP, FN, FP, Cand.num, Num.det, and *d_H_* denote the true positive rate, false negative rate, false positive rate, number of candidate change points, number of change points detected, scaled Hausdorff distance, respectively.

